# Functional benefit of CRISPR/Cas9-induced allele deletion for *RYR1* dominant mutation

**DOI:** 10.1101/2024.01.24.576997

**Authors:** Mathilde Beaufils, Margaux Melka, Julie Brocard, Clement Benoit, Nagi Debbah, Kamel Mamchaoui, Norma B. Romero, Anne Frédérique Dalmas-Laurent, Susana Quijano-Roy, Julien Fauré, John Rendu, Isabelle Marty

**Author notes:** Correspondence should be addressed to I.M., Grenoble Institut Neurosciences – Bat EJ Safra – Chemin Fortuné Ferrini -38700 La Tronche - France. equal contribution (John Rendu, Isabelle Marty).

## Abstract

More than 700 pathogenic or probably pathogenic variations have been identified in the *RYR1* gene causing various myopathies collectively known as “*RYR1*-related myopathies”. Currently, there is no treatment for these myopathies, and gene therapy stands out as one of the most promising approaches. In the context of a dominant form of Central Core Disease due to a *RYR1* mutation, we aimed at showing the functional benefit of inactivating specifically the mutated *RYR1* allele by guiding CRISPR/Cas9 cleavages onto frequent single nucleotide polymorphisms (SNPs) segregating on the same chromosome. Whole-genome sequencing was used to pinpoint SNPs localized on the mutant *RYR1* allele and identified specific CRISPR/Cas9 guide-RNAs. Lentiviruses encoding these guide-RNAs and the *SpCas9* nuclease were used to transduce immortalized patient muscle cells, inducing the specific deletion of the mutant *RYR1* allele. The efficiency of the deletion was assessed at both DNA and RNA levels and at the functional level after monitoring calcium release induced by the stimulation of the RyR1-channel. This study provides *in-cellulo* proof of concept regarding the benefits of mutant *RYR1* allele deletion, in the case of a dominant *RYR1* mutation, from both a molecular and functional perspective.

**Graphical abstract:** 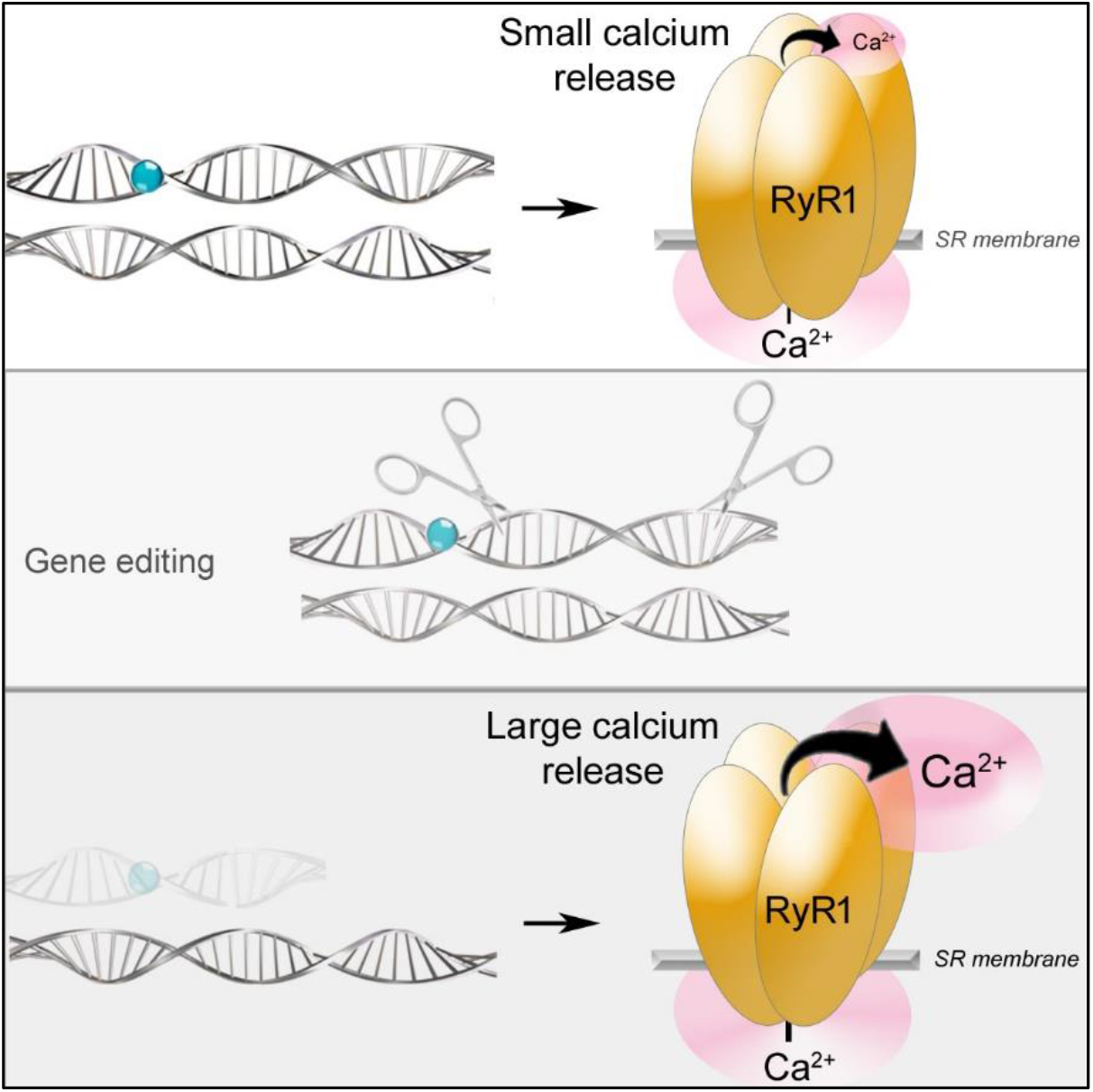

**eTOC synopsis:** Mutations in the *RYR1* gene, encoding a calcium channel required for muscle contraction, cause severe myopathies. In this study, Marty and colleagues demonstrate the functional benefit of suppression of a mutant *RYR1* allele using CRISPR/Cas9, in the case of a dominant mutation, leaving the wild type allele alone.

## Introduction

Muscle stimulation at the neuromuscular junction induces membrane depolarization, activating a macromolecular protein complex known as the calcium release complex (CRC). This complex, composed of the plasma membrane voltage-sensitive calcium channel (dihydropyridine receptor-DHPR) and the intracellular calcium channel (ryanodine receptor-RyR1), allows intracellular calcium release, leading to muscle contraction.^1^ The CRC’s function relies on the cross-talk between its two main components: the DHPR, activated by membrane depolarization, and the RyR1, releasing calcium from the sarcoplasmic reticulum into the cytosol upon stimulation.^1^ Mutations in the gene encoding the skeletal muscle ryanodine receptor *RYR1* result in altered intracellular calcium release through various pathophysiological mechanisms, including gain of function, loss of function, and reduction in protein amount.^1,2,3,4^ The human *RYR1* gene, spanning 150 kbp on chromosome 19, is transcribed into a 15 kbp mRNA corresponding to sequences of 106 exons.^5^ All exons except the alternative exons 70 and 83 are indispensable. The functional RyR1 calcium channel is a homotetramer, with each monomer containing 5037 amino acids in humans. Mutations in *RYR1* are distributed throughout the gene sequence,^6^ with over 1500 variations identified to date, accounting for dominant or recessive phenotypes (among which 700 are pathogenic or probably pathogenic, according to the ACMG classification^7^). *RYR1* mutations are responsible for various myopathies, including Central Core Disease (CCD), Multimini Core Disease (MmD), Dusty Core Disease (DuCD), Centronuclear Myopathy (CNM), collectively referred to as RyR1-related myopathies (RyR1-RM). These mutations are the most commonly found in patients with congenital myopathy.^2,3,8^ Patients typically present with muscle weakness of varying severity, from severe neonatal presentation with respiratory insufficiency to mild/isolated or generalized muscle weakness.^2^ Mutations in the *RYR1* gene can also result in Malignant Hyperthermia (MH), a hyperthermic and hypercontracture response triggered by anesthesia.^9^

Currently, there is no treatment for RyR1-RM, and various therapeutic strategies are under exploration.^4^ Gene therapy, although promising, faces challenges due to the *RYR1* coding sequence’s size, unsuitable for a single viral vector packaging. Additionally, truncating mutations with a protein which remains functional do not exist, ^10^ which makes so far gene transfer, potentially with a shorter version, not a possible option. Intervening at the mRNA level using antisense oligonucleotides or siRNA has shown promising results both *in cellulo*^11^ and *in vivo*. ^12^ However, these studies were limited to a single patient or family, as they targeted patient-specific mutations, making them overly personalized for widespread therapy. Gene editing using CRISPR/Cas9 is an active research field, with the potential to replace (thanks to Homology Directed Repair-HDR after DNA cleavage, or more recently thanks to base/prime editing) or delete (based on Non-Homologous End Joining-NHEJ) a mutant DNA.^13^ However, HDR-based correction is not applicable to non-dividing muscle fibers, limiting its use for myopathies. Successful deletion of a DNA segment or point correction with base/prime editing has been achieved in skeletal muscle.^14,15^ In this study, we show that CRISPR/Cas9-mediated DNA deletion in the *RYR1* gene is a potential therapeutic strategy for dominant RyR1-RM. Instead of targeting the patient’s mutation directly, we directed the nuclease cleavage to single nucleotide polymorphisms (SNPs) present on the same allele, inducing the exclusive loss-of-function of the mutant allele. We show that this strategy induces the extinction of the mutant allele in patient cells and a functional improvement in the calcium release. This mutation-independent allele deletion provides *in cellulo* proof of concept for the efficacy of CRISPR/Cas9 allele deletion in patients with dominant RyR1-RM.

## Results

### Patient description

The c.14387A>G variation resulting in the change of the tyrosine 4796 for a cysteine (p.Y4796C) in exon 100 of the *RYR1* gene has been previously identified in all the individuals of the family affected by a dominant form of RyR1-RM (Figure 1A and 1B).^16^ This neo-mutation affects individual II-8, who has two affected sons and two affected grand-sons (IV-1 and IV-2) and presents mild symptoms of Central Core Disease (walking ability maintained), associated with MH susceptibility, with typical images of cores at histological analysis of the muscle biopsy (Figure 1C). Her grand-son, (patient IV-1), presented as many classic dominant RYR1 mutated patients with congenital hip dislocation that was treated by surgery. He acquired walking ability, developed a spinal deformity that did not require surgery. At 17 years of age, at his last visit, he remained ambulant, showed no contractures and had normal pulmonary function tests. He had a non-progressive scoliosis (Cobb angle of 40°) with overall good trunk balance. His brother (patient IV-2) presented with a more severe phenotype, had congenital hip dislocation at birth which required early surgery and was never able to walk. He developed respiratory insufficiency and required bracing during childhood and spinal fusion at the end of the growth period. At his last clinical visit at 15 years of age, he had a moderately reduced vital capacity, without the need of nocturnal ventilator support (65% of normal values), showed hip and knee flexion contractures.

**Figure 1.**
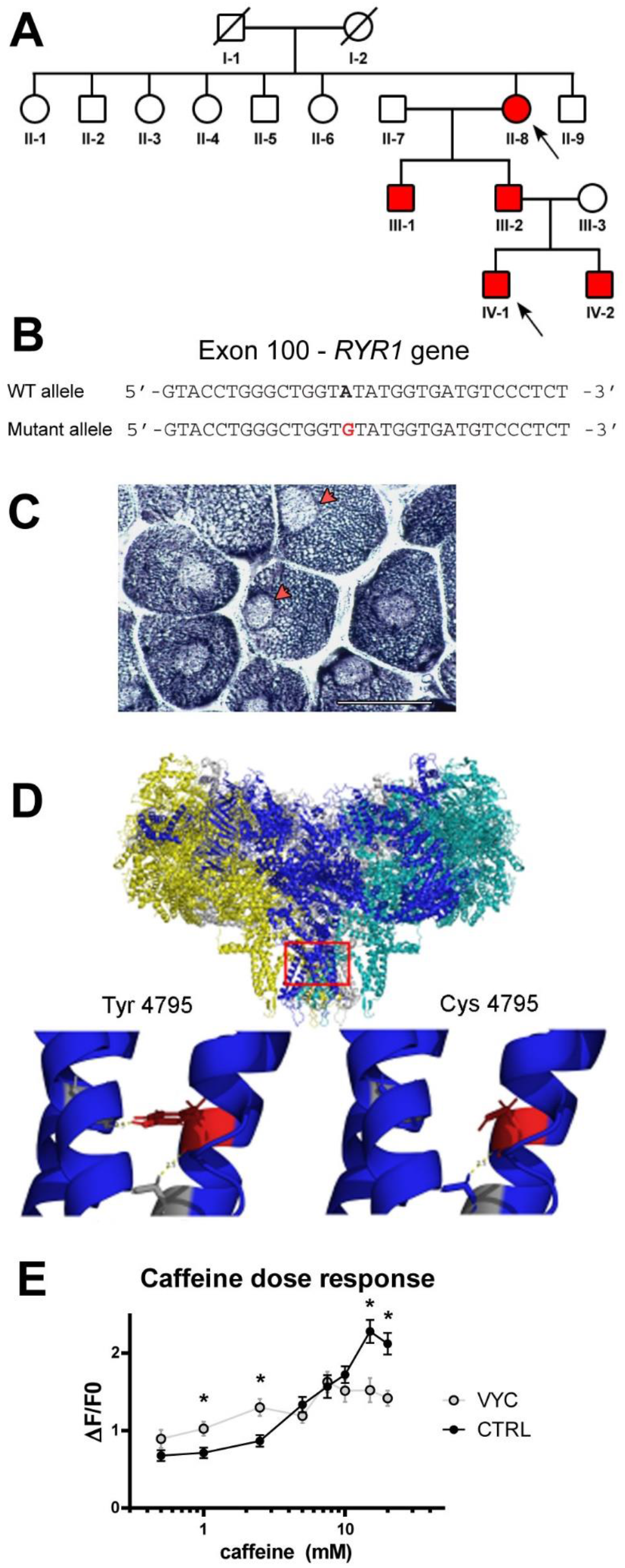
Description of the family and of the *RYR1* mutation. A) A dominant *RYR1* neo-mutation has been identified in patient II-8 (Monnier et al., 2000), resulting in the change of tyrosine 4796 for a cysteine. This mutation has been transmitted to the two sons, and to the two grandchildren, all being affected by Central Core Disease (affected patients are in red). B) At the DNA level, the mutation c.14387A>G is localized in exon 100 of *RYR1* gene. C) Histological analysis (NADH staining) shows the presence of cores (red arrows) in each muscle fiber of patient II-8, typical of Central Core Disease. Bar 50µm D) The structure of rabbit RyR1 and the position of the mutation in the pore of the channel (on the rabbit sequence, mutation is Y4795C). In presence of a cysteine in position 4795, the number of hydrogen bonds with the nearby α-helix is reduced compared to tyrosine. The reduction in the interaction between these two helixes in the pore probably results in a reduction in the strength of the closing of the channel. E) Mean peak of calcium release induced by caffeine stimulation on patient immortalized myotubes (VYC) or on control myotubes (CTRL), using Fluo-4 calcium imaging. For each concentration, values are presented as mean ± SEM of fluorescence variation measured on 85 to 170 myotubes. At low caffeine concentration (1mM - 2.5mM), a significantly higher calcium release, characteristic of MH sensitivity, is observed in the VYC patient cells compared to CTRL cells, whereas at high concentration (>15mM), a significantly lower calcium release is observed. Statistical analysis Student t-test between CRTL and VYC at each concentration: 0.5mM p=0.1077, 1mM p=0.0071, 2.5mM p=0.0019, 5mM p=0.2852, 7.5mM p=0.7634, 10mM p=0.2515, 15mM p=0.0005, 20mM p<0.0001; * p<0.05.

The c.14387A>G;p.Y4796C mutation is localized within the pore of the channel (Figure 1D), and the presence of the cysteine results in a reduction of the hydrogen bonds between two α-helixes in the pore. Ectopic expression of a RyR1 channel with this mutation in HEK cells has shown that the mutated RyR1 channel is hypersensitive to caffeine stimulation and presents a gain-of-function with a calcium leak resulting in depletion of the calcium store,^16^ leading to the associated diagnosis of Malignant Hyperthermia susceptibility.^17^ Primary myoblasts have been obtained from a muscle biopsy of patient II-8, and immortalized as described previously^18^ by a double retroviral transduction (hTERT and Cdk4) followed by clonal selection. These myoblasts, so called V-Y4796C or VYC cells, have been further used for the development of the proof of concept of the functional benefits of allele deletion. The hypersensitivity to caffeine stimulation of the mutant RyR1 has been confirmed on these myoblasts (Figure 1E), in straight line with the previously reported malignant hyperthermia susceptibility associated with this mutation.^16^

### Identification of the SNPs and gRNAs selection

It has been previously observed that one functional *RYR1* allele is sufficient for a normal muscle function in human. Indeed, in families with recessive form of RyR1-RM, individuals who harbor one loss-of-function allele and one normal *RYR1* allele have no clinical signs.^19^ This has been further confirmed in mice heterozygous for a *RYR1*-KO allele that show no phenotype.^20^ In addition, allele specific gene silencing using siRNA has demonstrated a functional benefit in two mouse models with dominant *RYR1* mutations resulting in CCD and MH.^12^ The goal of the project was therefore to show that switching-off the mutant *RYR1* allele while preserving the normal one was beneficial in muscle cells of a patient with a dominant form of RyR1-RM. The allele knock-down was induced using the S*p*Cas9 nuclease, and a double cleavage that would result in the deletion of an essential part of the *RYR1* gene associated to frameshift was designed. In order to develop a strategy that could further be applied to other mutations, the targets of nuclease were SNPs present on the same allele as the mutation, and not the mutation itself. We chose to target SNPs with a frequency >1% in order to develop a versatile tool that could be applied to other patients. The first step of the project was to identify the SNPs present on each allele, and to determine which ones could be used for the design of gRNA.

Whole genome sequencing (WGS) was performed on DNA from patients II-8 and IV-1, allowing the identification of all the SNPs present in the two patients. Only the SNPs present in the *RYR1* gene and shared by the two patients were further considered (Figure 2A). The SNPs with a frequency above 1%, present in the *RYR1* gene at an heterozygous state were further used for the screening. From the 79 SNPs identified, we selected the 14 SNPs in which the SNP resulted in the formation of a *Sp*Cas9 protospacer-adjacent motif (PAM) NGG (Figure 2A). These 14 SNPs were therefore present only on the mutant *RYR1* allele, and absent from the wild type (WT) allele. *Sp*Cas9 targeted cleavage would theoretically result in a cleavage 3 bases upstream the PAM sequence, and only on the mutant allele. Guides RNA (gRNAs) were further designed on these SNPs/PAM (Figure 2B), and 5 gRNAs were selected. As our goal was to knock-out specifically the mutant *RYR1* allele, gRNA were chosen to obtain two cleavage sites leading to deletion of at least one exon and a disruption of the reading frame in *RYR1*. Pairs of gRNAs were further formed and cloned in a lentiviral vector, to produce the so-called “Lenti-guides” as described before,^21^ expressing the two gRNAs independently. Five pairs were selected to produce Lenti-G1 to Lenti-G5, resulting in deletion of 1 to 70 exons of the *RYR1* gene (Figure 2C), from 1.740 bp to more than 90.000 bp.

**Figure 2.**
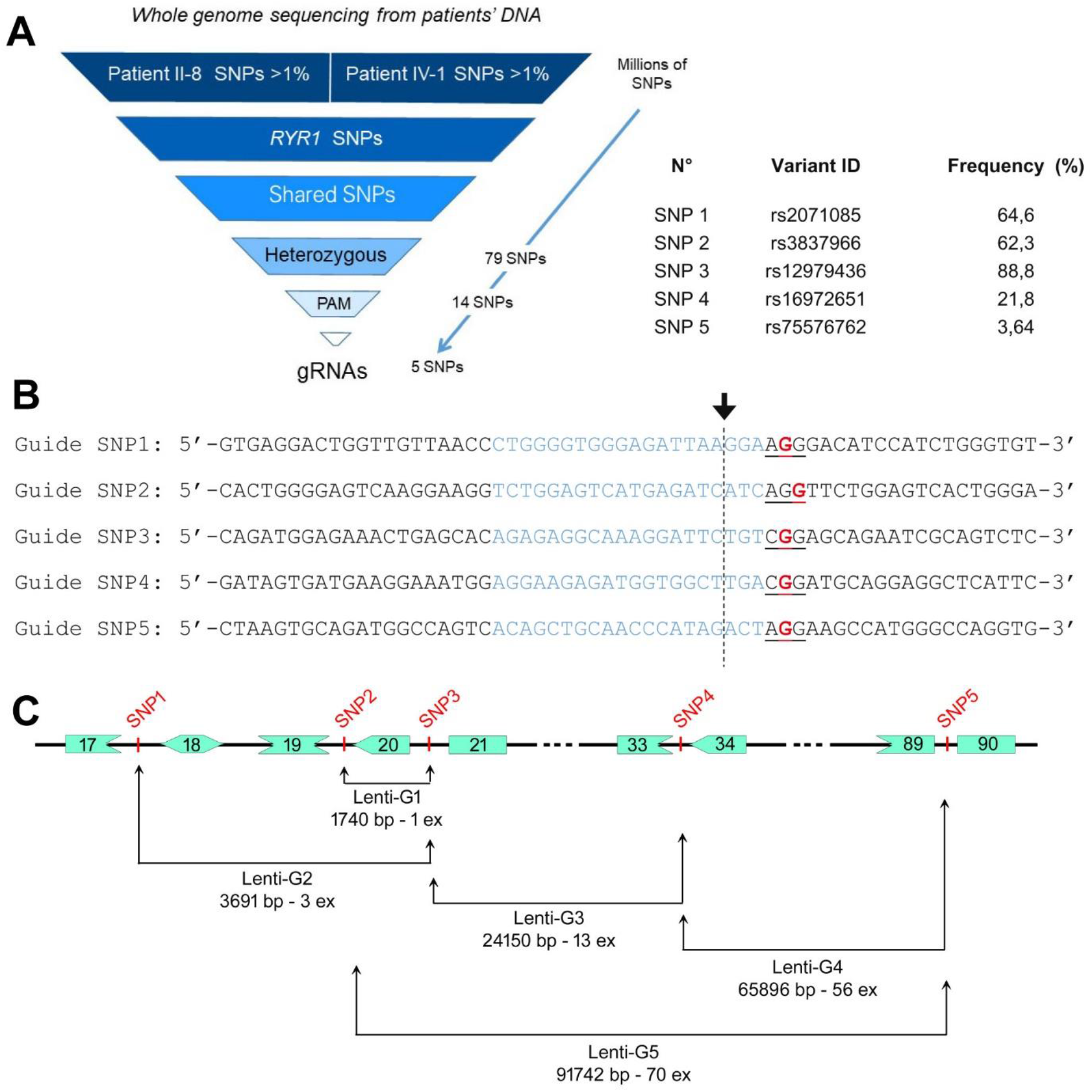
Pipeline for selection of the SNP and identification of gRNA. A) Whole genome sequencing has been performed on patient II-8 and patient IV-1 DNA, and the frequent SNPs in each patient’s genome collected. The SNPs were further filtered to select only the ones present in the *RYR1* gene, and common to both patients, at heterozygous state. At this step, the 79 SNPs present in the *RYR1* gene only on the mutant allele were identified. After an additional selection, only the 14 SNPs creating an *Sp*Cas9 PAM (NGG) were used to select the best possible gRNAs using Crispor tool (http://crispor.tefor.net/) based on specificity, efficiency and predicted off-target. Only 5 SNPs further resulted in the identification of gRNA of top quality (based on MIT score and off-target prediction). In parallel, a dedicated script has been developed to perform automated SNP screening and selection (available at https://github.com/clbenoit/CutOneStrand), leading to selection of the same gRNA. The frequency of the 5 selected SNP, determined using gnomAD v3.1.1 on January 11^th^, 2023, and the variant ID is presented on the right. B) The five best gRNAs corresponding to 5 heterozygous SNPs of the mutant *RYR1* allele were further used. Each SNP is represented in bold red, the PAM is underlined and the corresponding gRNA is in blue. The *Sp*Cas9 cleavage site is represented by the arrow and the dashed line. C) Pairs of gRNAs were further formed to produce lentiviruses called “Lenti-G” that lead to deletion of at least on exon and disruption of the reading frame. The localization of the selected SNP/gRNA is represented with reading frame of the surrounding exons, allowing visualizing the frameshift, and the size of expected deletion with each pair of guide.

### Validation of the selected gRNA and clones production

Immortalized muscle cells from the patient (VYC cells) were transduced with two lentivirus encoding respectively the *Sp*Cas9 (Lenti-Cas9) and one of the gRNAs pair (Lenti-G1 to 5), as described previously,^21^ and 5 days later, the cells were transduced with a so-called Lenti-Killer^21^ encoding a gRNA targeting the *SpCas9* gene, in order to stop the nuclease production.^22^ After a seven days amplification, the cells were collected and the deletion in the *RYR1* gene checked by PCR analysis of the targeted regions (Figure 3A and B, Table 1). Only Lenti-G2 resulted in a deletion in the *RYR1* gene, visualized by PCR amplification of the corresponding DNA region in the whole cell population (Figure 3C). The experiments were thus further performed only with cells treated with Lenti-G2, corresponding to a deletion of the 3 exons 18-20 (deletion of 3691bp, Figure 2C). A pool of cells treated with the 3 lentiviruses (Lenti-Cas9, Lenti-G2 and Lenti-Killer) in which the deletion of the chosen region has been evidenced by PCR was submitted to single cell cloning by limited dilution. After amplification, each individual clone was checked for *RYR1* gene editing using the same PCR amplification, followed by Sanger sequencing of the amplified region (Figure 3D).Two populations were clearly identified in the selected clones: edited clones (so called EC), presenting two bands after PCR amplification corresponding respectively to the WT allele and the deleted allele (such as EC-A, EC-B and EC-C, Figure 3D), and non-edited clones (NEC) presenting only one band after PCR amplification resulting of amplification of the two unmodified alleles (such as NEC-A, NEC-B, and NEC-C, Figure 3D). For the edited clones, the Sanger sequencing confirmed a cleavage 3bp upstream of the PAM and the deletion of the sequence in between (Figure 3E), corresponding to deletion of intron 17 to intron 20. After PCR amplification of the regions encompassing each guide and Sanger sequencing, no deletion was observed in non-edited clones. For the subsequent molecular and functional characterization, 3 edited clones (EC-A, EC-B and EC-C) and 3 non-edited clones (NEC-A, NEC-B, NEC-C) were selected.

**Figure 3.**
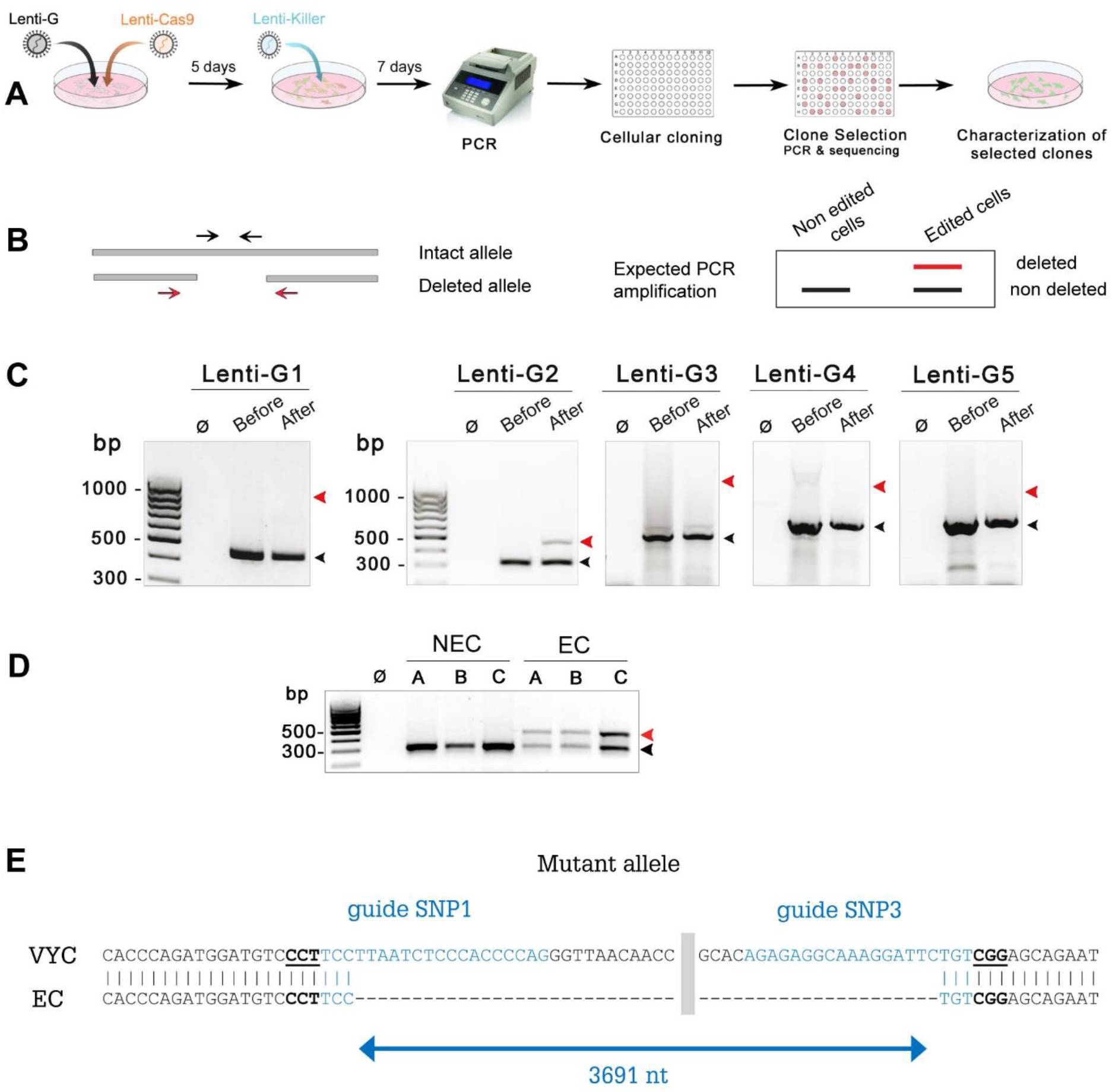
Validation of the gRNA and selection of clones. A) Workflow of the whole treatment of the cells to obtain edited cell lines. The patients cells were treated with two lentiviruses (Lenti-G1 to 5 and Lenti-Cas9), and 5 days later the nuclease was inhibited by Lenti-Killer. After 7 days, the edition efficiency was analyzed for each gRNA pair by PCR. When edition was observed the cells were cloned, amplified, and the editing controlled in each clone by PCR and sequencing. The functional consequence of *RYR1* edition was further analyzed by calcium imaging. B) For each Lenti-G (gRNA pair), primers were designed to amplify exclusively the intact allele (primer within the two cleavage sites, black arrows) or the deleted allele (primer on both side of the cleavage sites, red arrows) (Table 1). The red primers could theoretically amplify both the intact and the deleted alleles, but the fragment on the intact allele was too large to be amplify with the same PCR settings. The scheme on the right depicts the expected results of PCR amplification on edited and non-edited cells with the two sets of primers used together. Note that the red primers for the amplification of the deleted allele have been designed to produce a larger PCR product than the one corresponding to the intact allele produced with the black primers. C) PCR amplification of the edited (red primers) and non-edited allele (black primers) for each Lenti-G. PCR amplification has been performed without DNA (ø), or with DNA extracted from patient’s cells before and after treatment with Lenti-G and Lenti-Cas9. The DNA segment corresponding to the red primers cannot be amplified in the intact allele because it is too large. The size of the amplicons are respectively for lenti-G1: 440bp (intact allele, black arrow) and 985bp (deleted allele, red arrow); for lenti-G2 314bp (intact allele, black arrow) and 507bp (deleted allele, red arrow); for lenti-G3 470bp (intact allele, black arrow) and 1266bp (deleted allele, red arrow); for lenti-G4 659bp (intact allele, black arrow) and 1145bp (deleted allele, red arrow); for lenti-G5 619bp (intact allele, black arrow) and 1074bp (deleted allele, red arrow). D) PCR amplification of the edited and non-edited allele (red and black primers described above) after DNA purification of each individual clone. The edited clones (so called EC) present two bands after PCR amplification corresponding to the WT allele (lower band) and the deleted allele (upper band) (EC-A, EC-B, EC-C…), and the non-edited clones (NEC) present only one band after PCR amplification resulting of amplification of the two alleles (NEC-A, NEC-B, …). E) Sanger sequencing of the deleted PCR band produced in edited clones with the red primers, aligned on the sequence of the patient cells VYC. The sequence of each guide is represented in blue, and the PAM is underlined. The gray zone correspond to the sequence deleted not represented in VYC sequence. The Cas9 cleavage occurs 3bp upstream the PAM.

**Table 1:**
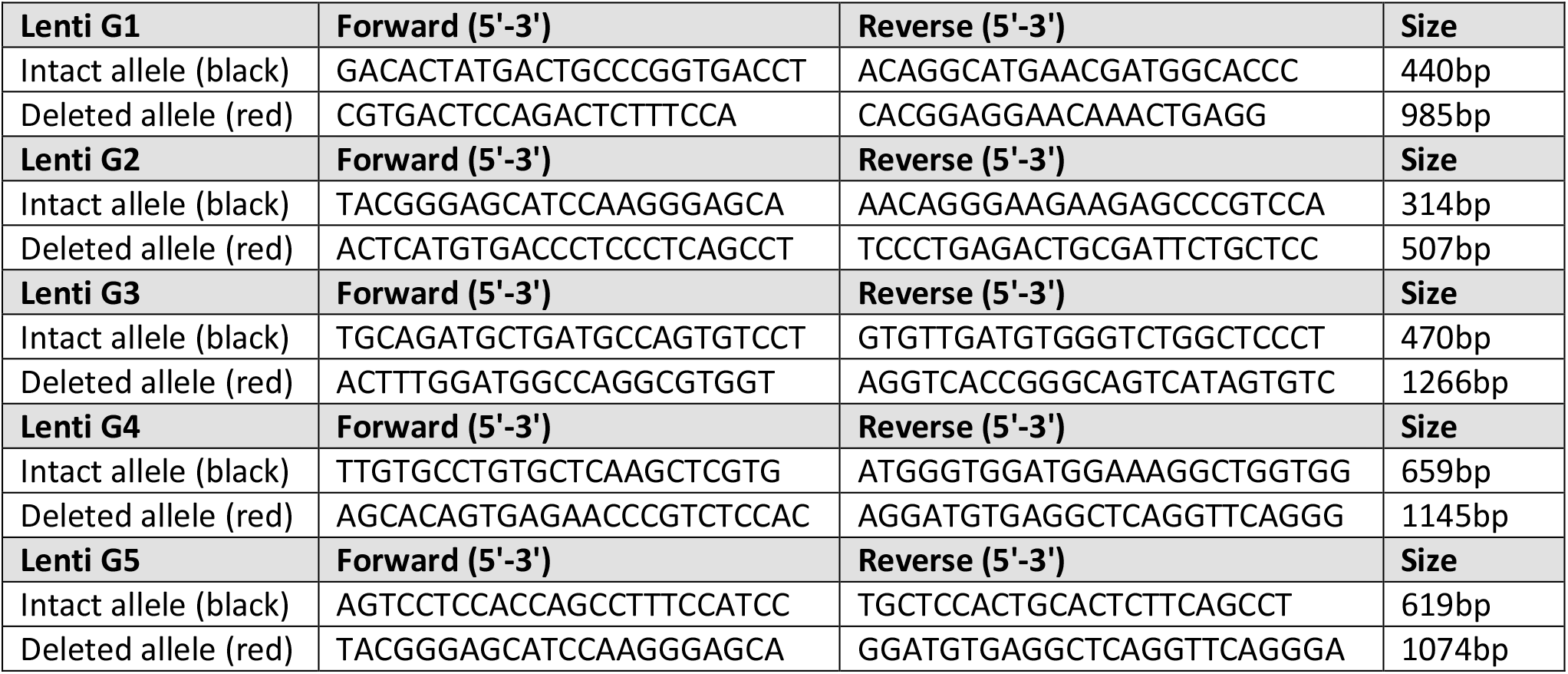
Primers for PCR amplification after cleavage with lenti-G1 to 5.

### Molecular characterization of the clones

In order to confirm the consequence of DNA deletion at the molecular level, and to confirm the deletion of the mutant allele, analysis of *RYR1* mRNA in the 3 edited and the 3 non edited clones was performed and compared to control cells (CTRL, immortalized myoblasts from a non affected individual) and VYC cells. RyR1 being expressed only in differentiated muscle cells, mRNA was extracted from myotubes produced for each clone after 7 days in differentiation. PCR amplification and NGS sequencing of the region encompassing the mutation demonstrated that in the CTRL, only a normal transcript is observed whereas in VYC cells, the WT and the mutant transcript are present in equivalent amounts. In the edited clones EC the relative amount of mutant transcript was reduced and the WT was increased whereas in the non-edited clones NEC as in the initial patient’s cells VYC, both the normal and the mutant transcripts were detected in equivalent amount (Figure 4A). In the edited clones, the deletion of the DNA region between the two guides lead to non-sense mediated decay (NMD) of the mutant allele (reduced from 50% to 33%), as estimated by quantification of the NGS reads corresponding to WT or mutant transcript (histogram 4A, right). The clones were further characterized at the protein level, using quantitative western blot. In order to be able to compare their function, their differentiation ability was estimated by the quantification of the amount of Myosin heavy chain (MYHC, expressed in differentiated myotubes) compared to GAPDH in each clone. No significant difference was observed between the EC, the NEC clones and the initial patient cells (Figure 4C, left histogram). The relative amount of RyR1 compared to MYHC was also estimated, and no difference between the EC and NEC clones was observed (Figure 4C, right histogram). The expression level of Cas9 was checked, in order to evaluate if after the whole procedure, and the addition of the Lenti-Killer, the Cas9 protein expression has been abolished as expected (Figure 4D). The Cas9 protein was not detected in any clones, EC or NEC.

**Figure 4:**
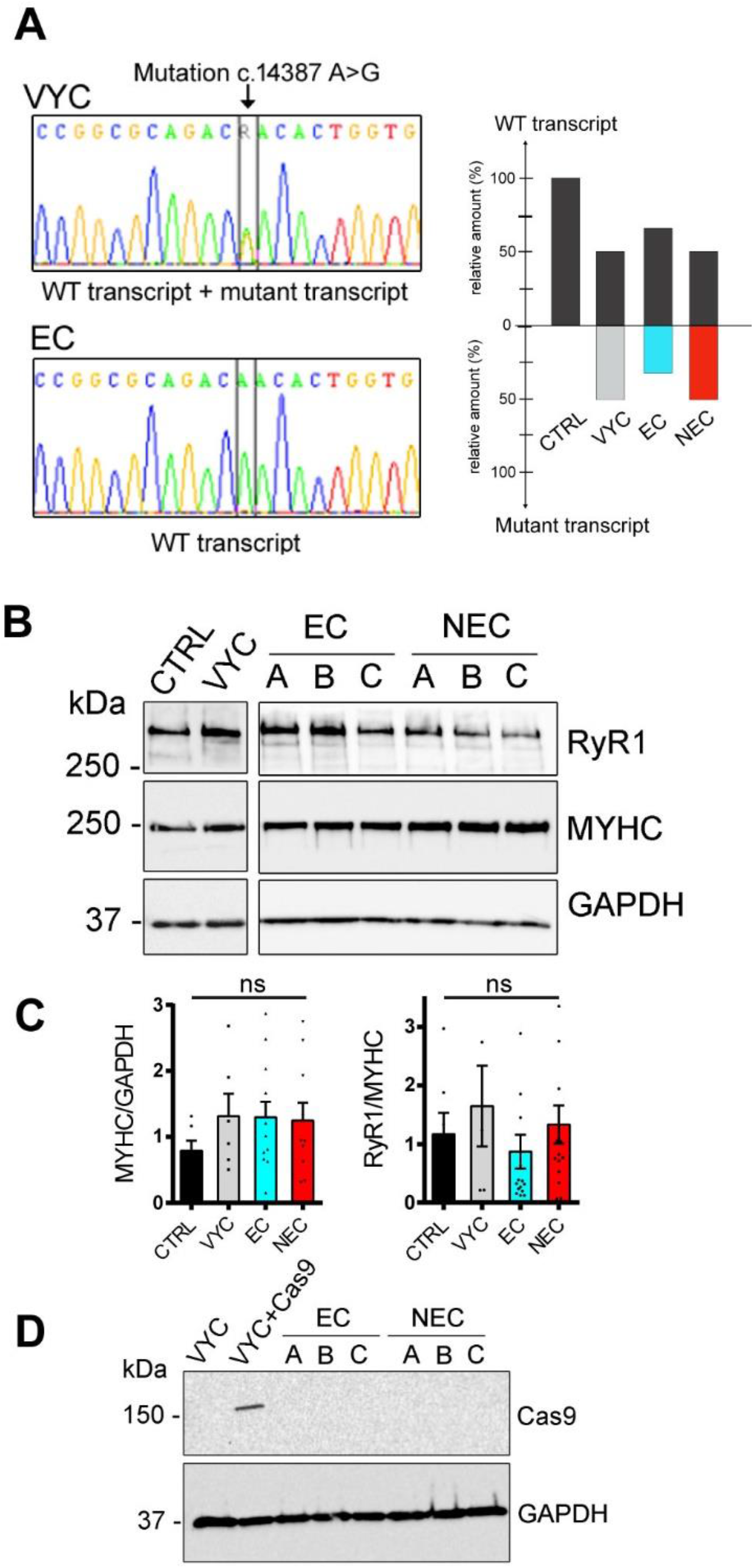
Molecular characterization of the cells. A) After mRNA extraction from differentiated myotubes and PCR amplification, the *RYR1* sequence encompassing the mutation c.14387A>G has been sequenced by NGS for the different clones. The electropherograms on the left show the presence of two transcripts in VYC cells, with an “A” and a “G” at the same position, whereas in EC only one transcript with an A is present. The bar graph on the right is a schematic representation based on the quantification of each transcript from NGS data. The relative amount of WT transcript is represented by the upper bar, and the relative amount of mutant transcript is the lower. In all the edited clones, the normal transcript (WT-upper black bar) is expressed in higher amount (66%) than the mutant transcript (lower blue bar, 33%), whereas in the non-edited clones, as in VYC cells, the two transcripts are observed in equivalent amount (WT, upper black bar 49% and mutant, lower red bar 51%). B) The analysis of different proteins has been performed using quantitative western blot in CTRL cells, patient’s cells VYC, 3 edited clones (EC-A to C) and 3 non edited clones (NEC-a to C). Representative image of 3 different western blots. C) Relative amount of Myosin/GAPDH confirmed the presence of equivalent amounts of Myosin heavy chain (MYHC) and indicates similar differentiation in the different clone. Similar relative amount of RyR1 protein are also observed in the different clones (RyR1/MYHC amount). Data are presented as the mean ± SEM of the 3 blots (CTRL and VYC) and for the 3 edited and the 3 non edited clones (EC and NEC). Statistical analysis: one-way ANOVA with Tukey’s multiple comparison test, ns: non significant difference. D) Western blot analysis of Cas9 protein in VYC cells, VYC transduced with lenti-Cas9 (VYC+Cas9) as a positive control, EC-A to -C and NEC-A to -C. This analysis confirms the absence of detectable amount of Cas9 protein in the different clones at the end of the treatment.

### Control of selectivity of the cleavage sites

The selected gRNAs have been chosen based on their individual predicted maximal efficiency and their minimal off-target cleavages. Therefore, none of the selected gRNAs presented any predicted off target cleavage with 0 or 1 mismatch. To further screen for potential off-target Cas9 cleavages, at least 10 DNA regions encompassing possible off-target cleavage sites with 2 or more mismatches, predicted by at least 2 different software, were amplified by PCR and sequenced. Three different software were used to identify the putative off-target sites for each gRNA (gRNA1 and gRNA2 for *RYR1*, and g-Killer for *Cas9*): Benchling (https://benchling.com/), Crispor (http://crispor.tefor.net/), and Cas-OFFinder (http://www.rgenome.net/cas-offinder/). For each gRNA the most probable first 10 to 13 sites predicted by at least two software were further selected for in depth analysis (Figure 5A). A total of 36 predicted off-target sites were therefore analyzed by PCR amplification (localization and primers presented in Table 2) followed by sequencing for each of the 3 EC and the 3 NEC. Among those, two sites corresponding to intergenic regions (OT11 and OT14) have not been successfully amplified and sequenced even after changing the primers. No off-target cleavage were identified in any of the predicted regions, in any of the clones (Figure 5B). In addition, using another cell line which has the two heterozygous SNPs on one allele (the AB1190 cell line, produced from another non-affected individual), we confirmed that the cleavage is observed when the two SNPs are present (AB1190 cells, Fig 5C), but no cleavage is observed when the two SNPs are not present (Figure 5C), such as in the CTRL cell line.

**Figure 5:**
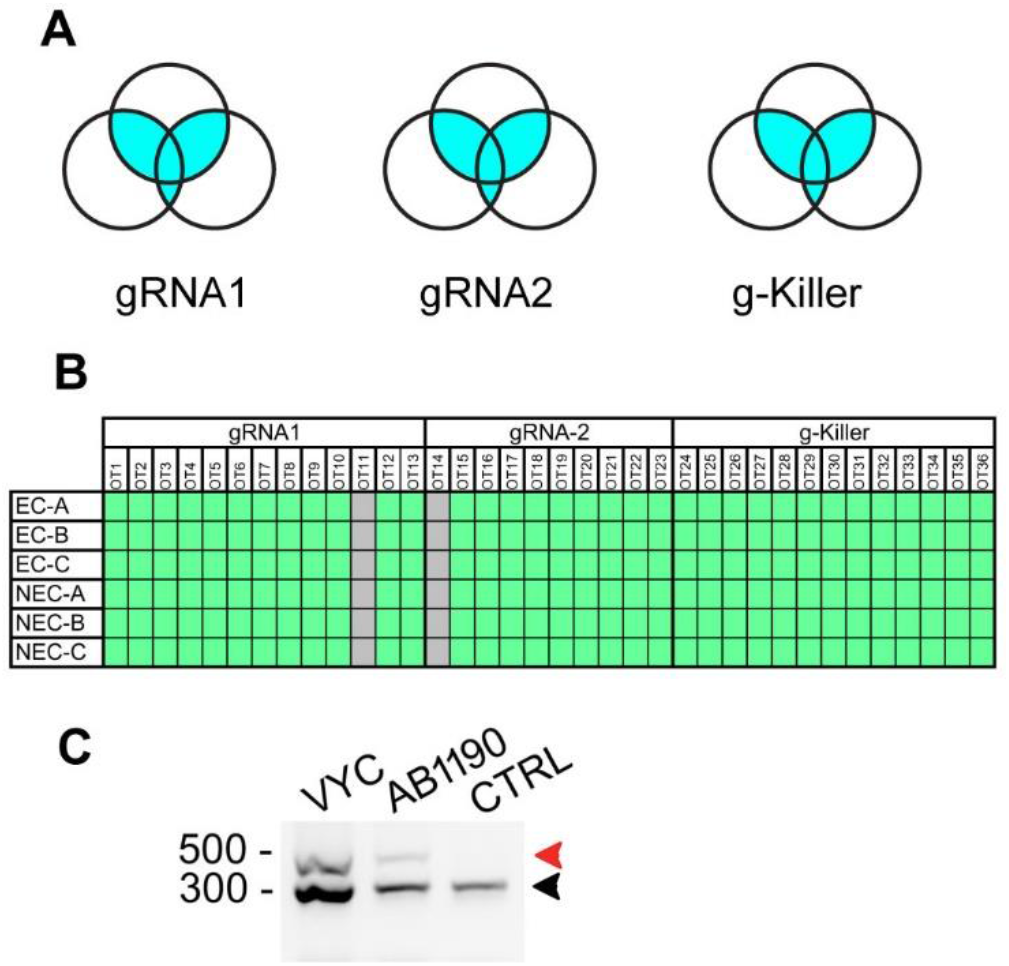
Specificity of the cleavages. A) Three different software were used to identify the putative off target sites for each gRNA (gRNA1 and gRNA2 for *RYR1*, and g-Killer for *Cas9*): Benchling, Crispor, and Cas-OFFinder. For each gRNA, the most probable first 10 to 13 sites predicted by at least two software were further selected for PCR amplification and sequencing. B) For each gRNA, 10 to 13 primer pairs were designed (so called OT1 to OT13 for gRNA1, OT14 to OT23 for gRNA2 and OT24 to OT36 for g-Killer, presented in table 2), and used for PCR amplification of the DNA extracted from the 3 EC cells (EC-A to -C) and the 3 NEC cells (NEC-A to -C). Two regions are represented in gray, which correspond to defective PCR amplification (OT11 and OT14, PCR amplification with additional primers also unsuccessful). The PCR fragments were further sequenced and the sequence of each region screened for modification. A green box corresponds to absence of modification, and a red box corresponds to insertion/deletion in the sequence. None of the predicted off-target sites presented any modification. C) The same procedure (transduction with Lenti-Cas9 and Lenti-G2) was applied to different human immortalized muscle cell lines, which contain the either two selected SNPs (VYC and AB1190) or only one SNP (CTRL), followed by DNA extraction and PCR amplification of the edited zone as in figure 3C. When the 2 SNPs are present (VYC and AB1190) the expected deletion can be observed, but when only one SNP is present (CTRL), no deletion is observed.

**Table 2:**
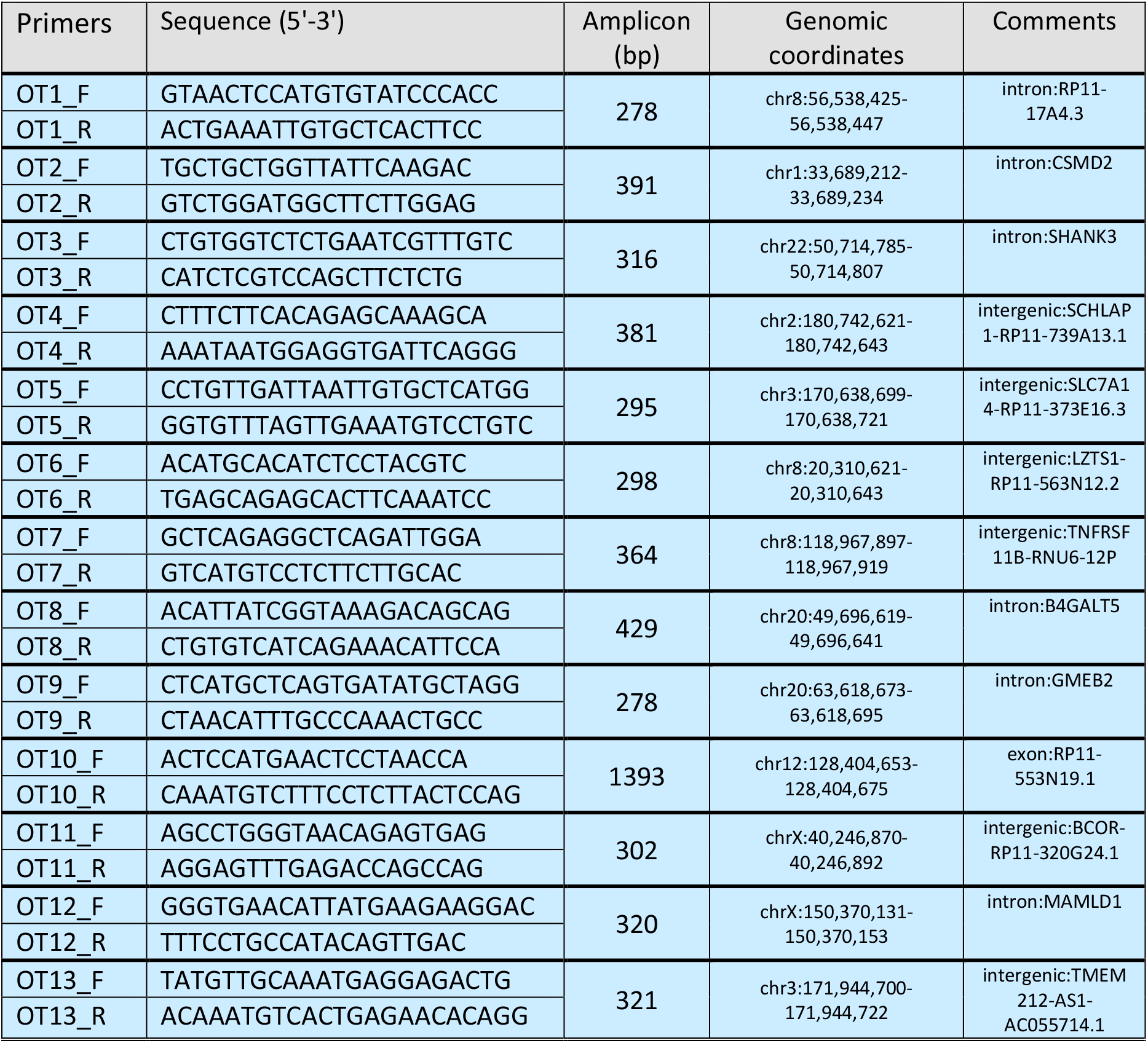
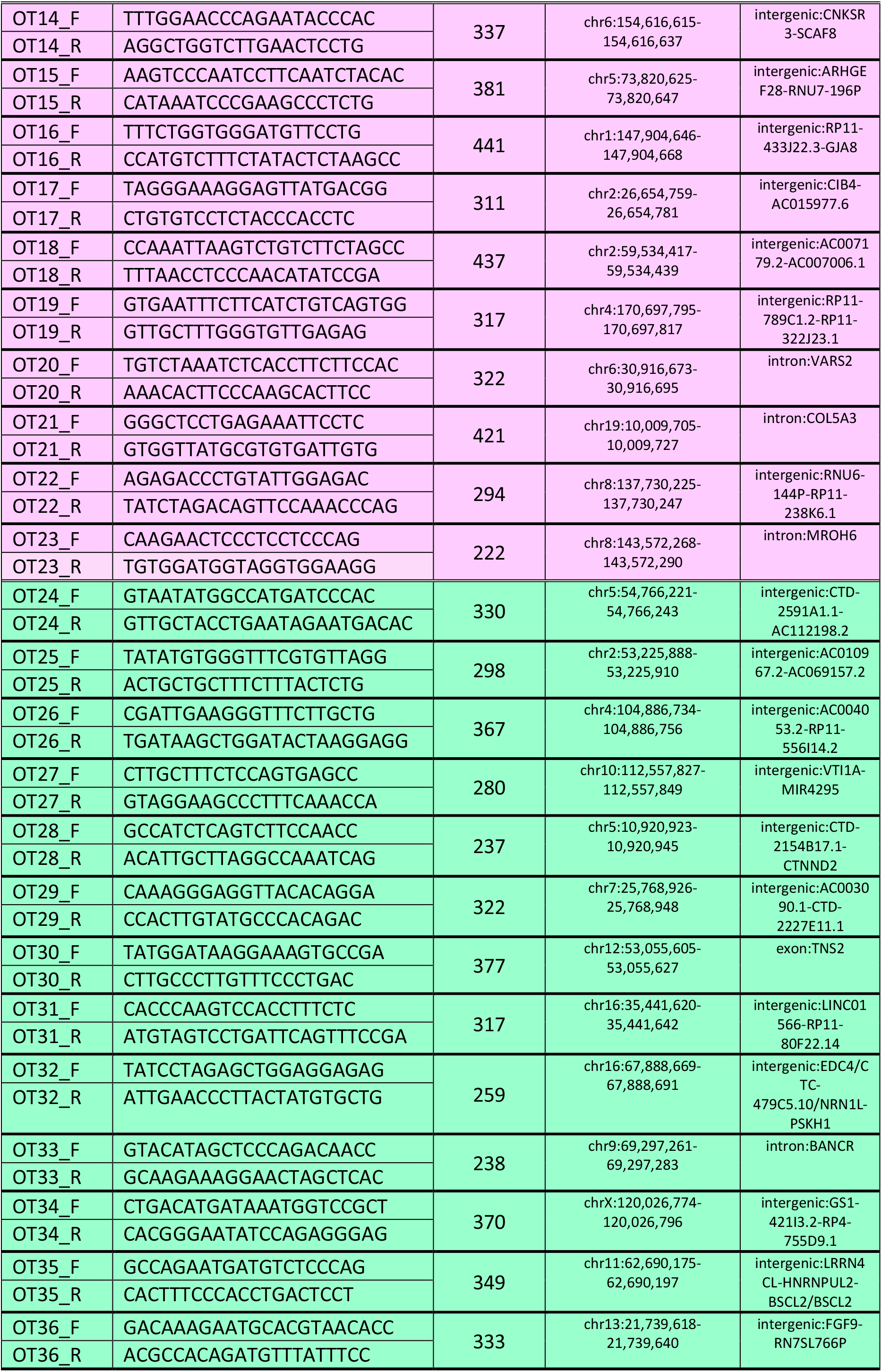
Primers for off-target PCR amplifications.

### Functional characterization of the selected clones

In the normal function of myotubes, membrane depolarization activates the voltage-activated calcium channel dihydropyridine receptor, which in turn activates the RyR1 by a direct molecular interaction between the two proteins,^23^ leading to a massive intracellular calcium release performed via RyR1. Therefore, to study the function of RyR1 after editing, calcium release has been studied using calcium imaging with the calcium sensitive dye Fluo-4 in the 3 edited clones and the 3 non-edited clones, compared to the initial patient cells VYC and CTRL cells. The mutation Y4796C re-expressed in HEK cells has previously been shown to result in a leaky RyR1, with hypersensitivity to caffeine and reduction in the amount of calcium stored into the sarcoplasmic reticulum.^16^ The effect of the mutation has now been directly studied in patient’s cells. Membrane depolarization induced by addition of KCl 140mM confirmed the reduction in the amount of calcium (reduction in peak amplitude and reduction in the area under the curve) in VYC cells compared to CTRL cells (Figure 6). The deletion of the mutant RyR1-allele in the edited clones EC induced a significant increase in the amplitude of the calcium release (Figure 6B), which became non significantly different from CTRL. The amount of calcium released, represented by the area under the curve, also increased but remained significantly different from CTRL (Figure 6C). Each individual EC had the same behavior (Supp Figure 1). In contrast, the non-edited clones NEC remained non significantly different from the patient’s cell VYC (Figure 6B and C), and all the 3 individual NEC clones behave identically (Supp Figure 1).

**Figure 6:**
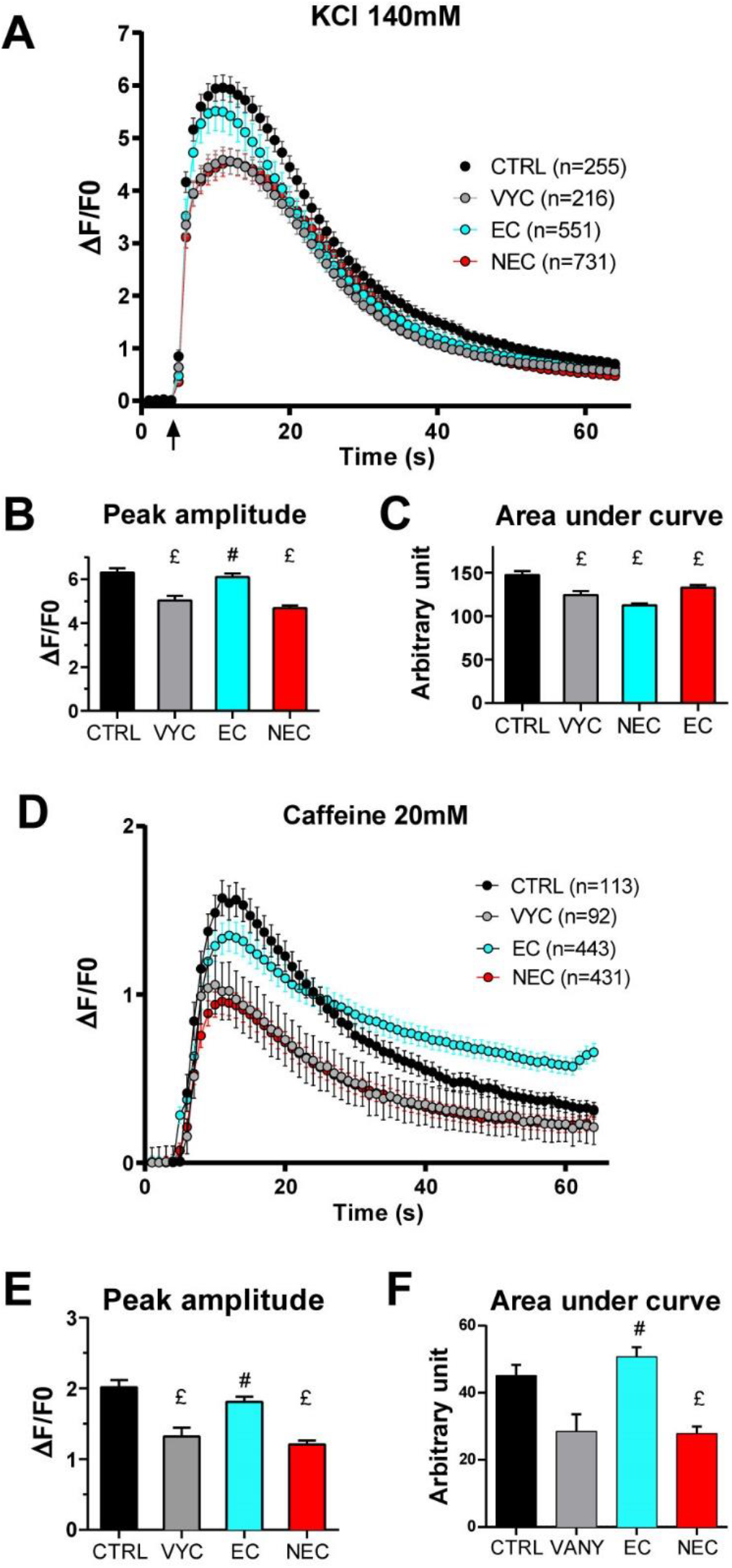
Functional characterization of the cells. Fluo-4 calcium imaging performed on myotubes produced from CTRL, VYC, EC or NEC cell lines. The curves represent the fluorescence variation in CTRL myotubes (black curve), VYC myotubes (gray curve), EC myotubes (blue curve) and NEC myotubes (red curve). All values are presented as mean ± standard error of mean (SEM) of n myotubes. In each condition, n = 180 to 255 myotubes have been analyzed, from at least three different experiments (exact number indicated for each curve). The values from the 3 EC and the 3 NEC lines have been pooled to simplify the presentation. The curves for each clone are presented individually in supplemental figure 1 A) Kinetics of calcium release upon stimulation by KCl 140mM. The stimulation is performed at 5s (arrow), and the fluorescence variation (ΔF/F0) recorded for 1min. The curves represent the mean ± SEM of fluorescence variation of the n different myotubes in each condition. B) Peak amplitude of calcium release after KCl stimulation presented as mean ± SEM of the maximum for each myotube. Statistical analysis: ANOVA with Tukey’s multiple comparison test. p value compared to CTRL: VYC p=0.0004, EC p=0.8588, NEC p<0.0001. p value compared to VYC: EC p=0.0008, NEC p=0.5398. £: significantly different from CRTL; #: significantly different from VYC. C) Mean area under the curve presented as mean ± SEM of the AUC for each myotube. Statistical analysis: ANOVA with Tukey’s multiple comparison test. p value compared to CTRL: VYC p=0.0003, EC p=0.0366, NEC p=0.0025. p value compared to VYC: EC p=0.1168, NEC p=0.4021. £: significantly different from CRTL. D) Kinetics of calcium release upon stimulation by caffeine 20mM E) Peak amplitude of calcium release after caffeine stimulation presented as mean ± SEM of the maximum for each myotube. Statistical analysis: ANOVA with Tukey’s multiple comparison test. p value compared to CTRL: VYC p=0.0011, EC p=0.4390, NEC p<0.0001. p value compared to VYC: EC p=0.0077, NEC p=0.8829. £: significantly different from CRTL; # significantly different from VYC. F) Mean area under the curve presented as mean ± SEM of the AUC for each myotube. Statistical analysis: ANOVA with Tukey’s multiple comparison test. p value compared to CTRL: VYC p=0.0925, EC p=0.7287, NEC p=0.0074. p value compared to VYC: EC p=0.0009, NEC p=0.9995. £: significantly different from CRTL; # significantly different from VYC.

Direct RyR1 stimulation by its agonist caffeine confirmed the hypersensitivity of patient cells VYC compared to CTRL cells and the normalization of this hypersensitivity by RyR1-editing (figure 6D, E, F). The 3 edited clones had the same behavior, and the 3 non-edited clones behave also identically (Supp Figure 1).

All these functional experiments therefore demonstrate that deletion of the mutant RyR1 allele in the edited clones reverses, at least partially, the abnormal calcium handling of the patient’s cells.

## Discussion

As of 2022, more than 700 variants in the *RYR1* gene have been classified as pathogenic or likely pathogenic, making it the most common gene associated with congenital myopathy and malignant hyperthermia.^10^ Over the period 2012-2022, during the genetic diagnosis performed in the Grenoble University Hospital, 161 pathogenic or likely pathogenic variations have been identified in the *RYR1* gene, with 120 variations of dominant transmission (RyR1-RM and MH). Unfortunately, there is currently no treatment available for any *RYR1* mutation. Among these variants, about 20% are associated to a dominant form of RyR1-RM and 54% to MH. The therapeutic approach presented here holds potential for application to any dominant pathology in which one functional allele is sufficient (MH and RyR1-RM, which reach about 75% of the variants). Searching the OMIM database, we identified a list of 571 genes associated to dominant mutations (Supp Figure 2), for which the strategy presented here could potentially apply if one allele is sufficient.

Whereas the complete knockout (KO) of the *RYR1* gene in mice has been reported to be lethal at birth,^24^ experimental evidences have demonstrated that a single wild-type (WT) allele is sufficient for normal muscle function. Specifically, heterozygous animals from an RyR1 KO mouse line showed that protein production with a single allele reaches 85% of the normal amount produced in WT animals.^20^ Moreover, mice with an RyR1 amount of 70% exhibited no muscle strength deficit, as observed in an inducible RyR1-KO mouse model.^25^ In humans, the presence of a null allele along with a normal *RYR1* allele has not been associated with muscle weakness, as seen in parents of patients with a recessive RyR1-RM.^19^ Therefore, the deletion of a mutant allele, leaving a single functional allele alone, will likely represent an effective therapeutic solution.

The therapeutic strategy developed here is based on the use of two guides targeting allele-specific and frequent single nucleotide polymorphisms (SNPs). In a similar approach, it has already been demonstrated *in vitro* a high mutant allele specificity for mutations in *TGFBI* gene linked to corneal dystrophy, with one gRNA targeting an heterozygous SNP, the second one targeting an intronic region present on both alleles.^26^ Thus, with two allele-specific gRNAs targeting the mutant *RYR1* allele, an even higher specificity with preservation of the WT allele was expected. Further experiments in different unrelated immortalized cell lines have confirmed that when one of the two targeted SNPs is absent, no deletion in the *RYR1* gene is observed, and the deletion occurs only when the two SNPs are present. This result strengthens the notion that when the two SNPs are absent, no deletion in the *RYR1* gene is observed, and conversely, when the SNPs are present, the deletion is observed, linking therapeutic efficiency to the presence of the SNPs rather than to the specific mutation. Consequently, patients with the same SNPs associated with a mutant allele of *RYR1* could potentially benefit from a similar therapeutic approach, regardless of the specific mutation on the gene, as long as the second functional allele lacks the targeted SNPs. It can also be proposed that a limited number of gRNA pairs targeting few SNPs have the potential to treat all patients with a dominant form of RyR1-RM. With the rapid evolution of whole genome sequencing, the SNPs present in each patient could be easily identified, and a limited number of pair of gRNA based on the geographic origin of the patients could be developed and tested for each gene to allow deletion of a single allele.

In parallel to the manual selection of the SNP/gRNA described here, a dedicated script, so called “CutOneStrand” has been developed for automated SNP screening and gRNA selection on one allele (https://github.com/clbenoit/CutOneStrand and Supplemental methods). Similar results have been obtained with the manual and the automated selection, the automated selection reducing considerably the time for SNPs selection. This script could be used for any of the 571 genes presented in Supp Figure 2, in order to induce the specific deletion of one allele, if one allele has been confirmed as sufficient.

The concept of deleting the mutant allele in the case of a dominant pathology has been previously observed using a different methodological approach with siRNA, not only for *RYR1*^12^ but also for *DNM2* mutations.^27^ In these cases, siRNA was designed to specifically target the mutation.^12,27^ The strategy has been further refined and proven efficiency for *DNM2* with siRNA targeting SNPs.^28^ Thus, this therapeutic strategy has the potential to be applicable for 75% of RyR1 patients. In the further extension to *in vivo* therapy, the use of viral vectors for the introduction of the CRISPR/Cas9 tools will allow the efficient targeting of all the muscles, even the most difficult muscles for injection, such as respiratory muscles (diaphragm and intercostal muscles). The constant improvement of viral vectors for systemic delivery in therapeutic use, increasing the desire tissue targeting (specifically muscle) and reducing the unexpected delivery to other organs such as liver, will increase the efficiency of a viral based gene therapy,^29^ allowing the combination of multiple vectors with preserve efficiency, because each muscle cell will receive all the viral vectors. The progressive deletion of the Cas9 induced by the gRNA against the Cas9 (the “Killer”) that can be combined to the other gRNAs under a weaker promoter^22^ and injected at the same time can also be used *in vivo* to avoid the long term expression of Cas9. Alternative methods such as nanoparticles could also be used to introduce the Cas9 as a protein, and not as a DNA, thus limiting its efficient to the natural protein turn over and avoiding the use of the Killer.

The positive impact of deleting the *RYR1* mutant allele has been directly observed in the intensity of calcium release upon stimulation. Now, the next step will be to translate this observation into muscle strength improvement, which will necessitate *in vivo* testing. The correlation between the amplitude of calcium release in cultured cells and the *in vivo* muscle strength has now to be established. As the patients with a same RyR1 dominant mutation have a congenital myopathy of various severity (mild in individual II-8 and more severe in individual IV-2), the efficiency of such a treatment in the context of various severity will have to be considered.

## Materials and methods

### SNPs identification and gRNA selection

DNA was extracted from blood samples of patients II-8 and IV-1 using the Nucleospin extraction kit (ref 740952-Macherey-Nagel). Subsequently, whole-genome sequencing was conducted on the DNA of each patient using Illumina HiSeq Analysis Software v2.1. Alignment was obtained by the Isaac aligner (Isaac-04.17.06.15) and variant calling was realized using Isaac variant caller workflow, the reference genome being GRCh38. SNPs from the two genomes were then filtered to retain those found in both patients with a frequency above 1% in the *RYR1* gene and heterozygous. Among the 79 remaining SNPs, only those creating one of the two “G” in an “NGG” PAM were selected, resulting in a list of 14 SNPs. Guides-RNA were then chosen from this list using CRISPOR software^30^ (http://crispor.tefor.net). Only the best gRNA matching the following criteria were further utilized: a MIT score of at least 50 out of 100, no predicted off-targets for 0 or 1 mismatch, and not classified as inefficient (according to Graf and coll.^31^). In the end, five SNPs were selected along with their corresponding gRNA (SNP 1 to 5).

### Bioinformatic analysis

A computational pipeline, named CutOneStrand, has been implemented in Bash and Python, using various bioinformatics tools. It integrates a series of scripts to select and analyze SNPs with sequence modifications at PAM sites for CRISPR-Cas9 genome editing. After uploading the sequence of Chr19 containing the *RYR1* gene (GRCh38), the script is design to identify all the heterozygous SNPs with a frequency >1% and creating or deleting a spCas9 PAM (NGG). The output is a file with the best guides corresponding to these SNPs. The overview of the pipeline is presented in Supplemental Methods.

### 3D molecular modeling

The molecular modeling has been developed based on the Protein Data Bank Entry 7M6L,^32^ depicting the open conformation of the RyR1 protein, as the structural template.^33^ The mutation modeling was enacted through the Prime software within the Schrodinger Suite, employing the “side chain prediction and residue minimization” protocol.^34^ The mutation process adhered to rigorous parameters, including the implementation of the OPLS 2005 forcefield.^35^ A 5.0 Angstrom minimization cutoff was applied, coupled with the automatic utilization of conjugate gradient and truncated Newton methods. The minimization protocol, executed with a maximum of 2 iterations, sought convergence based on a desired final RMS gradient of 0.10, an energy cutoff of 0.100 kcal/mol, and a maximum of 65 truncated Newton iterations per step.

Following the mutation procedure, the structural ramifications were quantified through key metrics. Delta energy and Delta SASA were computed, elucidating shifts in energetic stability and solvent-accessible surface area. The mutation process culminated in the generation of a distinct structural configuration, encapsulating cumulative mutational effects. To enhance result interpretation, PyMol, an integral visualization tool within the Schrodinger Suite, was employed (Schrödinger and DeLano 2020; http://www.pymol.org/pymol). This facilitated a comparative analysis between the original and mutated structures, with particular focus on the spatial relationship between Alanine 4566 and Tyrosine-Cystein 4795. Distance measurements within PyMol provided quantitative insights into the conformational disparities induced by the mutations, thus enriching the structural characterization.

### Plasmids and lentiviruses transduction

Lentiviruses were produced by a triple transfection of HEK 293T cells. Lenti-guide encoding gRNA against the chosen SNPs (Lenti-G) or the Cas9 gene (Lenti-Killer, as described by Merienne and coll. ^22^) were produced using Addgene plasmid #87919. Lenti-Cas9 encoding Cas9 was produced from Addgene plasmid #87904. The immortalized myoblasts were transduced in proliferation medium with the two lentiviruses Lenti-G and Lenti-Cas9 at a multiplicity of infection (MOI) 10, and 7 days later the Cas9 action was stopped by transduction with Lenti-Killer (MOI 20) as described previously.^21^

### Cell culture and differentiation

HEK293T cells were maintained in medium composed of Dulbecco’s Modified Eagle Medium (DMEM) high glucose pyruvate, supplemented with 10 % fetal bovine serum (Life technologies) and 2 % penicillin/streptomycin (Life technologies).

Immortalized human satellite cells (myoblasts) have been produced as described previously^18^ from a 25 years control individual (CTRL^36^) or from a muscle biopsy of patient II-8 (so-called here V-Y4796C or VYC cells). The myoblasts were amplified in proliferation medium composed of Ham’s F-10 (Life technologies) supplemented with 20% Fetal Bovine Serum (FBS) (Life technologies), 2% Ultroser G (Pall-Sartorius) and 2% Penicillin-Streptomycin (Life technologies). Differentiation into myotubes was induced by a shift to differentiation medium: DMEM low glucose (Life technologies) supplemented with 2% heat inactivated Horse Serum (Life technologies) and 1% Penicillin-Streptomycin. Myoblasts were amplified in proliferation medium composed of F-10 nutrient medium (Ham’s F10) supplemented with 20 % fetal bovine serum, 2 % ultroser G serum, 2 % penicillin/streptomycin. Myoblasts are kept at a confluence 50-60% max.

### Cellular cloning

After trypsination, the myoblasts have been diluted in proliferation medium at 10 cells/ml and seeded at 1 cell/well in 96-well plates containing 100 μl/well of proliferation medium. During 2 to 6 weeks, the clones have been progressively amplified to larger plates until reaching at least a 35 mm plate, while maintaining the confluency of the myoblasts below 50%.

### DNA and RNA extraction on cultured cells

The cells have been collected with TrypLE (ref 12605-010, Life Technologies) and DNA extracted with the Nucleospin tissue kit following the manufacturer recommendations (ref 740952, Macherey-Nagel) for large amount of cells (300.000 cells). For smaller amount of cells (*i.e.* during clone selection), the following classical protocol has been used for DNA purification. The cells have been lysed in 500µl lysis buffer supplemented with proteinase K 100µg/ml (Tris/HCl 10mM pH=7.5, EDTA 10mM, NaCl10mM, N-lauryl sarcosine 0.5% + 100µg/ml proteinase K). After 2h at 60°C, DNA was precipitated by addition of 1.5ml of precipitation buffet (NaCl 150mM-EtOH 100%), collected by centrifugation and washed with EtOH 70%.

RNA has been extracted from myotubes differentiated for 7 days using Trizol (Life Technologies) followed by Chloroform/Isoamyl alcool (49:1 v/v). RNA has been retro-transcribed to cDNA using oligo-dT and Transcriptor reverse transcriptase (Roche) according to manufacturer recommendations.

### DNA and RNA analysis

PCR **a**mplification of the DNA has been performed with Phusion high fidelity master mix with GC buffer (ref F532L, ThermoFisher). The primers for PCR amplification have been selected with PerlPrimer.^37^ The different primers pairs are presented in Table 1.

If a sequencing was further needed, after separation on a 1% agarose gel, the bands of interest were excised and purified with the Gel and PCR clean up (ref 740609, Macherey-Nagel) with a final elution volume of 20 µl, and submitted to Sanger sequencing (Eurofins genomics).

Total *RYR1* transcripts were amplified as 7 overlapping PCR products.^38^ They were sequenced after fragmentation and library preparation using NEBNEXT NGS workflow (New England Biolabs) according to manufacturer recommendation. The library was then sequenced on a S5 IonTorrent platform (ThermoFisher). FastQ files were analysed using nf-core pipeline (nf-core/rnaseq v3.0), alignments and splice junctions were produced using RNAStar.

### Protein analysis

Myoblasts (200.000) were seeded on a laminin-coated 35mm culture dish and differentiated for 7 days as described previously.^21^ The proteins were collected by cell lysis in RIPA (25mM Tris-HCl pH 7.6, 150mM NaCl, 1% NP-40, 1% sodium deoxycholate, 0.1% SDS) with protease inhibitors (200 mM phenylmethylsulfonyl fluoride, and 1 mM diisopropyl fluorophosphate). Protein quantification was performed using BCA kit (Pierce), and 20µg was loaded in each well of a 5-15% acrylamide gel for western blot analysis. The amount of RyR1 compared to Myosin Heavy chain (MHC) as a marker of differentiation was estimated using quantitative western blot, as described before,^21^ using a rabbit polyclonal antibody against RyR1^23^ and a monoclonal anti-MHC antibody (MF20, DSHB, Iowa City). Antibody against GAPDH was used as a loading control (anti GAPDH (14C10) rabbit mAb, Cell Signaling Technology). The presence of the Cas9 protein was tested using antibody against the V5 tag (ref R960-25, Invitrogen). Secondary antibodies used for Western blot were labelled with HRP (Jackson Immuno Research). Signal quantification was performed using a ChemiDoc Touch apparatus (Bio-Rad) and the Image Lab software (Bio-Rad). The amount of the chosen protein in each sample was corrected for differences in loading using the amount of GAPDH.

### Calcium imaging

Human myotubes (CTRL cells, V-Y4796C cells, 3 edited clones EC-A, EC-B and EC-C and 3 non-edited clones NEC-A, NEC-B, NEC-C) were cultured for seven to eight days before intracellular calcium measurements. Changes in intracellular calcium were measured on the cultured myotubes, as described previously, ^39^ using the calcium-dependent fluorescent dye Fluo 4-Direct (Molecular Probes) diluted in differentiation medium. Calcium imaging was performed in Krebs buffer (136 mM NaCl, 5 mM KCl, 2 mM CaCl_2_, 1 mM MgCl_2_, 10 mM HEPES, pH7.4). KCl stimulation (140 mM final concentration) was performed by application of Krebs in which NaCl was replaced by KCl (140 mM KCl, 2mM CaCl_2_, 1mM MgCl_2_, 10mM HEPES, pH7.4). Caffeine was diluted at 20mM final concentration in Krebs. The curves represent the mean ± S.E.M of fluorescence variation after stimulation during 65 sec of the n myotubes in each condition, from at least 3 different experiments. The peak amplitude is the mean ± S.E.M of the maximal amplitude of each myotube (n myotubes in the same condition, from 3 different experiments), and the area under the curve (AUC) is the mean ± S.E.M of the area under the fluorescence curve for each myotube.

### Statistics

The statistical analysis has been done with GraphPad Prism 6.0 software. The number of samples and the name of the parametric test applied are indicated in each figure legend. Results are considered as significant when p < 0.05, the exact value for p being indicated in the text or figure legends, and significant results are labeled on the graphs whatever the exact p value. All data are shown as mean ± SEM.

## Supporting information

Supplemental figure 2, list of genes

## Data availability

The data are available upon reasonable request to the corresponding author.

## Acknowledgments

We thank the Myoline Plateform (Myology Institute, Paris) for the immortalized human satellite cells, the French rare diseases Healthcare Network for neuromuscular diseases - FILNEMUS and the European Reference Network Euro-NMD. The work was supported by grants from Association Française contre les Myopathies (AFM-Téléthon), Inserm and from Auvergne-Rhône-Alpes Region.

## Author contributions

MB, JF, JR and IM conceived and designed study. MB, MM and JB conducted the experiments and analyzed the results. CB and ND conducted bioinformatics and structural analyses. KM produced the immortalized myoblasts. NBR, AFDL, SQR recruited the patient, collected blood and muscle samples, and analyzed the patients’ phenotype. IM wrote the manuscript. JF provided expert feedback. All authors contributed to manuscript review.

## Abbreviations

ACMG: American College of Medical Genetics
CCD: Central Core Disease
CNM: Centronuclear Myopathy
CRC: calcium release complex
CRISPR/Cas9: clustered regularly interspaced short palindromic repeats/*CRISPR*-associated protein 9
CTRL: Control
DHPR: dihydropyridine receptor
DNM2: Dynamin 2
DuCD: Dusty Core Disease
EC: Edited clones
GAPDH: Glyceraldehyde-3-phosphate dehydrogenase
gRNA: guide RNA
HDR: homology directed repair
KO: Knock-out
MH: Malignant hyperthermia
MmD: Multimini core disease
MYHC: Myosin heavy chain
NEC: Non edited clones
NHEJ: non homologous end joining
PAM: protospacer adjacent motif
OMIM: Online Mendelian Inheritance in Man
OT: Off-target
RyR1: ryanodine receptor type 1
RyR1-RM: RyR1-related myopathy
siRNA: small interfering RNA
SNP: single nucleotide polymorphism
WGS: Whole Genome sequencing
WT: wild type.

## Supplemental Material

**Supplemental Figure 1:**
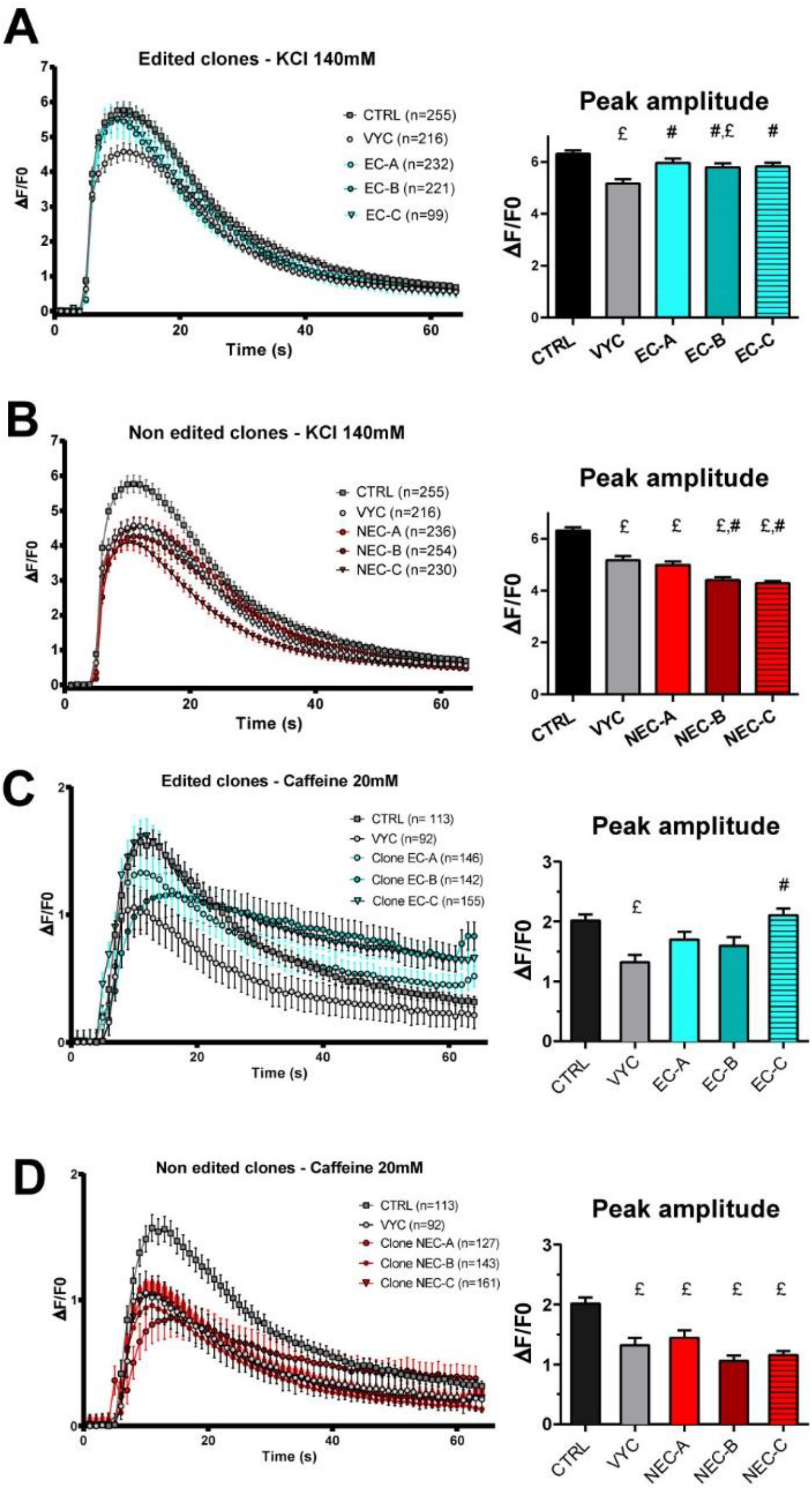
Calcium imaging on each individual clone.

**Legend Supp Fig 1**

Fluo-4 calcium imaging performed on myotubes produced from CTRL, VYC, EC-A to -C or NEC-A to -C cell lines. All values are presented as mean ± standard error of mean (SEM) of n myotubes. In each condition, n = 99 to 254 myotubes have been analyzed, from at least three different experiments (exact number indicated for each curve). Statistical analysis: One way ANOVA with Dunnett’s multiple comparison test; £: significantly different from CTRL; #: significantly different from VYC. A) Kinetics of calcium release upon stimulation by KCl 140mM in the 3 edited cell lines (EC-A, EC-B and EC-C) represented in blue, compared to CTRL myotubes (dark gray curve) and VYC myotubes (light gray curve).) Kinetics of calcium release upon stimulation by KCl 140mM in the 3 non edited cell lines (NEC-A, NEC-B and NEC-C) represented in red, compared to CTRL myotubes (dark gray curve) and VYC myotubes (light gray curve). C) Kinetics of calcium release upon stimulation by Caffeine 20mM in the 3 edited cell lines (EC-A, EC-B and EC-C) represented in blue, compared to CTRL myotubes (dark gray curve) and VYC myotubes (light gray curve). D) Kinetics of calcium release upon stimulation by Caffeine 20mM the 3 non-edited cell lines (NEC-A, NEC-B and NEC-C) represented in red, compared to CTRL myotubes (dark gray curve) and VYC myotubes (light gray curve).

**Supplemental Figure 2: List of all the genes potentially eligible to a similar strategy**

Excel file: Genes with dominant mutations.xlsx

**Legend Supp Fig 2**

All the genes present in the human genome and affected by dominant mutations have been screened in the OMIM database, and if a single allele could be enough, these genes are eligible to a similar strategy. The table presents the gene symbol (alphabetic order), the Entrez Gene ID, the Ensemble Gene ID and the phenotypes associated to mutation in the gene.

## Supplemental Methods

### The CutOneStrand script

The script is available at https://github.com/clbenoit/CutOneStrand. This computational pipeline integrates a series of scripts to select and analyze SNPs with sequence modifications at PAM sites for CRISPR-Cas9 genome editing. The pipeline is implemented in Bash and Python, utilizing various bioinformatics tools.

The bioinformatics tools required and their use is described below:

*Python 3: Primary programming language for scripting; Pandas: Data manipulation and analysis; Regular Expressions (re): String matching and manipulation; Argparse: Command-line argument parsing; VCF Module: Processing Variant Call Format (VCF) files; OS Module: Operating system interactions; Conda: Environment and package management ; Samtools, Bedtools, Picard, JVARKIT: Bioinformatics tools for genomic data processing; FlashFry: CRISPR target identification tool*.

The input Files and Parameters are as follow:

*FASTA File: Contains SNP alleles and flanking sequences; VCF File: SNP candidates in VCF format; Mutation List: Mutations of interest in relation to PAM; Candidates FASTA File: Genomic context for target sites; CRISPR Cas Variants: Details of Cas9 variants for PAM identification*.

The algorithm and procedures performed are the following:

*Setup and Parameter Parsing: Bash script parses input parameters including gene, Cas9 variant, variant frequency, and output file; Software Environment Setup: Installation and activation of necessary bioinformatics tools and Python environments; Data Retrieval and Preprocessing: Downloading and preparing chromosomal reference sequences, gene annotations, and SNP data from GnomAD; SNP Filtering: Filtering SNPs based on frequency and gene annotations; Genomic Context Retrieval: Extracting genomic context for candidate SNPs using JVARKIT; SNP Selection: Python script selectSNPs.py selects SNPs modifying CRISPR cutting sites; CRISPR Target Identification: Using FlashFry (McKenna and Shendure, 2018) for discovering off-targets and computing CRISPR target scores; Data Merging and Formatting: Python script flashFryResultsSNPinfosMerging.py merges FlashFry results with SNP data; Output Generation: Producing final results in the specified output file*.

The output Files are presented as follow:

*Selected SNP Data: Information on selected SNPs, including genomic context and CRISPR target data; FlashFry Results: Detailed CRISPR target and off-target information*.

