## Supplemental figure 2, list of genes for "Functional benefit of CRISPR/Cas9-induced allele deletion for *RYR1* dominant mutation"

| Approved Gene Symbol | Entrez Gene ID | Ensembl Gene ID | Phenotypes |
|---|---|---|---|
| AARS1 | 16 | ENSG00000090861 | Developmental and epileptic encephalopathy 29, 616339 (3), Autosomal recessive; Charcot-Marie-Tooth disease, axonal, type 2N, 613287 (3), Autosomal dominant; ?Leukoencephalopathy, hereditary diffuse, with spheroids 2, 619661 (3), Autosomal dominant; Trichothiodystrophy 8, nonphotosensitive, 619691 (3), Autosomal recessive |
| ABCA1 | 19 | ENSG00000165029 | Tangier disease, 205400 (3), Autosomal recessive; HDL deficiency, familial, 1, 604091 (3), Autosomal dominant |
| ABCA4 | 24 | ENSG00000198691 | recessive; {Macular degeneration, age-related, 2}, 153800 (3), Autosomal dominant; Cone-rod dystrophy 3, 604116 (3), Autosomal recessive; Fundus flavimaculatus, 248200 (3), Autosomal recessive; Stargardt disease 1, 248200 (3), Autosomal recessive |
| ABCB4 | 5244 | ENSG00000005471 | 614972 (3), Autosomal dominant, Autosomal recessive; Cholestasis, progressive familial intrahepatic 3, 602347 (3), Autosomal recessive |
| ABCC6 | 368 | ENSG00000091262 | Pseudoxanthoma elasticum, 264800 (3), Autosomal recessive; Arterial calcification, generalized, of infancy, 2, 614473 (3), Autosomal recessive; Pseudoxanthoma elasticum, forme fruste, 177850 (3), Autosomal dominant |
| ABCC8 | 6833 | ENSG00000006071 | recessive; Diabetes mellitus, transient neonatal 2, 610374 (3); Diabetes mellitus, noninsulin-dependent, 125853 (3), Autosomal dominant; Hypoglycemia of infancy, leucine-sensitive, 240800 (3), Autosomal dominant; Hyperinsulinemic hypoglycemia, familial, 1, 256450 (3), Autosomal dominant, Autosomal recessive |
| ABCC9 | 10060 | ENSG00000069431 | 239850 (3), Autosomal dominant; ?Atrial fibrillation, familial, 12, 614050 (3), Autosomal dominant; Intellectual disability and myopathy syndrome, 619719 (3), Autosomal recessive |
| ACAN | 176 | ENSG00000157766 | ?Spondyloepiphyseal dysplasia, Kimberley type, 608361 (3), Autosomal dominant; Short stature and advanced bone age, with or without early-onset osteoarthritis and/or osteochondritis dissecans, 165800 (3), Autosomal dominant; Spondyloepimetaphyseal dysplasia, aggrecan type, 612813 (3), Autosomal recessive |
| ACD | 65057 | ENSG00000102977 | ?Dyskeratosis congenita, autosomal recessive 7, 616553 (3), Autosomal dominant, Autosomal recessive; ?Dyskeratosis congenita, autosomal dominant 6, 616553 (3), Autosomal dominant, Autosomal recessive |
| ACKR1 | 2532 | ENSG00000213088 | [Blood group, Duffy system], 110700 (3), Autosomal dominant, Autosomal recessive; [White blood cell count QTL], 611862 (3), Autosomal recessive; {Malaria, vivax, protection against}, 611162 (3) |
| ACO2 | 50 | ENSG00000100412 | Autosomal recessive |
| ACOX1 | 51 | ENSG00000161533 | recessive |
| ACTA1 | 58 | ENSG00000143632 | Congenital myopathy 2B, severe infantile, autosomal recessive, 620265 (3), Autosomal recessive; ?Myopathy, scapulohumeroperoneal, 616852 (3), Autosomal dominant; Congenital myopathy 2C, severe infantile, autosomal dominant, 620278 (3), Autosomal dominant; Congenital myopathy 2A, typical, autosomal dominant, 161800 (3), Autosomal dominant |
| ACTL6B | 51412 | ENSG00000077080 | Developmental and epileptic encephalopathy 76, 618468 (3), Autosomal recessive; Intellectual developmental disorder with severe speech and ambulation defects, 618470 (3), Autosomal dominant |
| ADAR | 103 | ENSG00000160710 | Autosomal recessive |
| ADCY5 | 111 | ENSG00000173175 | with hyperkinetic movements and dyskinesia, 619651 (3), Autosomal recessive; Dyskinesia with orofacial involvement, autosomal recessive, 619647 (3), Autosomal recessive |
| ADGRV1 | 84059 | ENSG00000164199 | Usher syndrome, type 2C, 605472 (3), Digenic dominant, Autosomal recessive; Usher syndrome, type 2C, GPR98/PDZD7 digenic, 605472 (3), Digenic dominant, Autosomal recessive; ?Febrile seizures, familial, 4, 604352 (3), Autosomal dominant |
| ADRB3 | 155 | ENSG00000188778 | {Obesity, susceptibility to}, 601665 (3), Multifactorial, Autosomal dominant, Autosomal recessive |
| AFG3L2 | 10939 | ENSG00000141385 | Spastic ataxia 5, autosomal recessive, 614487 (3), Autosomal recessive; Optic atrophy 12, 618977 (3), Autosomal dominant; Spinocerebellar ataxia 28, 610246 (3), Autosomal dominant |
| AFP | 174 | ENSG00000081051 | Autosomal recessive |

| | | | |
|---|---|---|---|
| AGRP | 181 | ENSG00000159723 | {Leanness, inherited}, 601665 (3), Multifactorial, Autosomal dominant, Autosomal recessive; {Obesity, late-onset}, 601665 (3), Multifactorial, Autosomal dominant, Autosomal recessive |
| AIPL1 | 23746 | ENSG00000129221 | Leber congenital amaurosis 4, 604393 (3), Autosomal dominant, Autosomal recessive; Retinitis pigmentosa, juvenile, 604393 (3), Autosomal dominant, Autosomal recessive; Cone-rod dystrophy, 604393 (3), Autosomal dominant, Autosomal recessive |
| AIRE | 326 | ENSG00000160224 | Autoimmune polyendocrinopathy syndrome , type I, with or without reversible metaphyseal dysplasia, 240300 (3), Autosomal dominant, Autosomal recessive |
| ALB | 213 | ENSG00000163631 | ?[Dysalbuminemic hypertriiodothyroninemia], 615999 (3), Autosomal dominant, Autosomal recessive; Analbuminemia, 616000 (3), Autosomal recessive; [Dysalbuminemic hyperthyroxinemia], 615999 (3), Autosomal dominant, Autosomal recessive |
| ALDH18A1 | 5832 | ENSG00000059573 | 219150 (3), Autosomal recessive; Spastic paraplegia 9B, autosomal recessive, 616586 (3), Autosomal recessive; Cutis laxa, autosomal dominant 3, 616603 (3), Autosomal dominant |
| ALG8 | 79053 | ENSG00000159063 | Congenital disorder of glycosylation, type Ih, 608104 (3), Autosomal recessive; Polycystic liver disease 3 with or without kidney cysts, 617874 (3), Autosomal dominant |
| ALPL | 249 | ENSG00000162551 | Autosomal recessive; Hypophosphatasia, childhood, 241510 (3), Autosomal recessive; Hypophosphatasia, adult, 146300 (3), Autosomal dominant, Autosomal recessive |
| ALX4 | 60529 | ENSG00000052850 | Parietal foramina 2, 609597 (3), Autosomal dominant; {Craniosynostosis 5, susceptibility to}, 615529 (3), Autosomal dominant; Frontonasal dysplasia 2, 613451 (3), Autosomal recessive |
| ANK1 | 286 | ENSG00000029534 | Spherocytosis, type 1, 182900 (3), Autosomal dominant, Autosomal recessive |
| ANO5 | 203859 | ENSG00000171714 | Muscular dystrophy, limb-girdle, autosomal recessive 12, 611307 (3), Autosomal recessive; Miyoshi muscular dystrophy 3, 613319 (3), Autosomal recessive; Gnathodiaphyseal dysplasia, 166260 (3), Autosomal dominant |
| ANTXR1 | 84168 | ENSG00000169604 | Autosomal dominant |
| AP1G1 | 164 | ENSG00000166747 | Usmani-Riazuddin syndrome, autosomal recessive, 619548 (3), Autosomal recessive; Usmani-Riazuddin syndrome, autosomal dominant, 619467 (3), Autosomal dominant |
| AP4E1 | 23431 | ENSG00000081014 | Autosomal recessive |
| APOA1 | 335 | ENSG00000118137 | Hypoalphalipoproteinemia, primary, 2, 618463 (3), Autosomal recessive; Amyloidosis, 3 or more types, 105200 (3), Autosomal dominant; Hypoalphalipoproteinemia, primary, 2, intermediate, 619836 (3), Autosomal dominant |
| APOA2 | 336 | ENSG00000158874 | recessive |
| APOB | 338 | ENSG00000084674 | recessive |
| APOE | 348 | ENSG00000130203 | {?Alzheimer disease, protection against, due to APOE3-Christchurch}, 607822 (3), Autosomal dominant; {Coronary artery disease, severe, susceptibility to}, 617347 (3); Lipoprotein glomerulopathy, 611771 (3); {?Macular degeneration, age-related}, 603075 (3), Autosomal dominant; Hyperlipoproteinemia, type III, 617347 (3) |
| AQP2 | 359 | ENSG00000167580 | Diabetes insipidus, nephrogenic, 2, 125800 (3), Autosomal dominant, Autosomal recessive |
| ARL3 | 403 | ENSG00000138175 | Retinitis pigmentosa 83, 618173 (3), Autosomal dominant; Joubert syndrome 35, 618161 (3), Autosomal recessive |
| ATAD3A | 55210 | ENSG00000197785 | Harel-Yoon syndrome, 617183 (3), Autosomal dominant, Autosomal recessive; Pontocerebellar hypoplasia, hypotonia, and respiratory insufficiency syndrome, neonatal lethal, 618810 (3), Autosomal recessive |
| ATM | 472 | ENSG00000149311 | susceptibility to}, 114480 (3), Somatic mutation, Autosomal dominant; T-cell prolymphocytic leukemia, somatic (3); Lymphoma, mantle cell, somatic (3) |
| ATP1A2 | 477 | ENSG00000018625 | microcephaly, polymicrogyria, and dysmorphic facies, 619602 (3), Autosomal recessive; Alternating hemiplegia of childhood 1, 104290 (3), Autosomal dominant; Migraine, familial basilar, 602481 (3), Autosomal dominant; Migraine, familial hemiplegic, 2, 602481 (3), Autosomal dominant |
| ATP2B2 | 491 | ENSG00000157087 | (3), Autosomal recessive |

| | | | |
|---|---|---|---|
| ATP5F1A | 498 | ENSG00000152234 | Mitochondrial complex V (ATP synthase) deficiency, nuclear type 4A, 620358 (3), Autosomal dominant; ?Combined oxidative phosphorylation deficiency 22, 616045 (3), Autosomal recessive; ?Mitochondrial complex V (ATP synthase) deficiency, nuclear type 4B, encephalopathic type, 615228 (3), Autosomal recessive |
| ATP6V0A1 | 535 | ENSG00000033627 | Neurodevelopmental disorder with epilepsy and brain atrophy, 619971 (3), Autosomal recessive; Developmental and epileptic encephalopathy 104, 619970 (3), Autosomal dominant |
| ATP6V1A | 523 | ENSG00000114573 | Cutis laxa, autosomal recessive, type IID, 617403 (3), Autosomal recessive; Developmental and epileptic encephalopathy 93, 618012 (3), Autosomal dominant |
| ATP8B1 | 5205 | ENSG00000081923 | Cholestasis, progressive familial intrahepatic 1, 211600 (3), Autosomal recessive; Cholestasis, intrahepatic, of pregnancy, 1, 147480 (3), Autosomal dominant; Cholestasis, benign recurrent intrahepatic, 243300 (3), Autosomal recessive |
| ATR | 545 | ENSG00000175054 | [illegible] Autosomal dominant |
| B2M | 567 | ENSG00000166710 | ?Amyloidosis, familial visceral, 105200 (3), Autosomal dominant; Immunodeficiency 43, 241600 (3), Autosomal recessive |
| BEST1 | 7439 | ENSG00000167995 | Macular dystrophy, vitelliform, 2, 153700 (3), Autosomal dominant; ?Microcornea, rod-cone dystrophy, cataract, and posterior staphyloma 2, 193220 (3), Autosomal dominant; Retinitis pigmentosa-50, 613194 (3); Retinitis pigmentosa, concentric, 613194 (3); Vitreoretinochoroidopathy, 193220 (3), Autosomal dominant; Bestrophinopathy, autosomal recessive, 611809 (3) |
| BFSP1 | 631 | ENSG00000125864 | Cataract 33, multiple types, 611391 (3), Autosomal dominant, Autosomal recessive |
| BLVRA | 644 | ENSG00000106605 | Hyperbiliverdinemia, 614156 (3), Autosomal dominant, Autosomal recessive |
| BMP2 | 650 | ENSG00000125845 | dominant; Brachydactyly, type A2, 112600 (3), Autosomal dominant; {HFE hemochromatosis, modifier of}, 235200 (3), Autosomal recessive |
| BMPR1B | 658 | ENSG00000138696 | Acromesomelic dysplasia 3, 609441 (3), Autosomal recessive; Brachydactyly, type A2, 112600 (3), Autosomal dominant; Brachydactyly, type A1, D, 616849 (3), Autosomal dominant |
| BRCA1 | 672 | ENSG00000012048 | Fanconi anemia, complementation group S, 617883 (3), Autosomal recessive; {Breast-ovarian cancer, familial, 1}, 604370 (3), Multifactorial, Autosomal dominant; {Pancreatic cancer, susceptibility to, 4}, 614320 (3) |
| BRCA2 | 675 | ENSG00000139618 | recessive; {Medulloblastoma}, 155255 (3), Somatic mutation, Autosomal dominant, Autosomal recessive; {Prostate cancer}, 176807 (3), X-linked, Somatic mutation, Autosomal dominant; {Breast-ovarian cancer, familial, 2}, 612555 (3), Autosomal dominant; {Breast cancer, male, susceptibility to}, 114480 (3), Somatic mutation, Autosomal dominant; {Pancreatic cancer 2}, 613347 (3); Wilms tumor, 194070 (3), Somatic mutation, Autosomal dominant |
| BSCL2 | 26580 | ENSG00000168000 | Lipodystrophy, congenital generalized, type 2, 269700 (3), Autosomal recessive; Neuronopathy, distal hereditary motor, autosomal dominant 13, 619112 (3), Autosomal dominant; Silver spastic paraplegia syndrome, 270685 (3), Autosomal dominant; Encephalopathy, progressive, with or without lipodystrophy, 615924 (3), Autosomal recessive |
| BUB1B | 701 | ENSG00000156970 | Colorectal cancer, somatic, 114500 (3); [Premature chromatid separation trait], 176430 (3), Autosomal dominant; Mosaic variegated aneuploidy syndrome 1, 257300 (3), Autosomal recessive |
| C19orf12 | 83636 | ENSG00000131943 | Neurodegeneration with brain iron accumulation 4, 614298 (3), Autosomal dominant, Autosomal recessive; ?Spastic paraplegia 43, autosomal recessive, 615043 (3), Autosomal recessive |
| C3 | 718 | ENSG00000125730 | C3 deficiency, 613779 (3), Autosomal recessive; {Hemolytic uremic syndrome, atypical, susceptibility to, 5}, 612925 (3), Autosomal dominant; {Macular degeneration, age-related, 9}, 611378 (3) |
| C5 | 727 | ENSG00000106804 | C5 deficiency, 609536 (3), Autosomal recessive; [Eculizumab, poor response to], 615749 (3), Autosomal dominant |
| CACNA1D | 776 | ENSG00000157388 | Primary aldosteronism, seizures, and neurologic abnormalities, 615474 (3), Autosomal dominant; Sinoatrial node dysfunction and deafness, 614896 (3), Autosomal recessive |
| CACNA1S | 779 | ENSG00000081248 | {Thyrotoxic periodic paralysis, susceptibility to, 1}, 188580 (3), Autosomal dominant; Congenital myopathy 18 due to dihydropyridine receptor defect, 620246 (3), Autosomal dominant, Autosomal recessive; Hypokalemic periodic paralysis, type 1, 170400 (3), Autosomal dominant; {Malignant hyperthermia susceptibility 5}, 601887 (3), Autosomal dominant |

| | | | |
|---|---|---|---|
| CAMK2A | 815 | ENSG00000070808 | Intellectual developmental disorder, autosomal dominant 53, 617798 (3), Autosomal dominant; ?Intellectual developmental disorder, autosomal recessive 63, 618095 (3), Autosomal recessive |
| CAPN3 | 825 | ENSG00000092529 | Muscular dystrophy, limb-girdle, autosomal recessive 1, 253600 (3), Autosomal recessive; Muscular dystrophy, limb-girdle, autosomal dominant 4, 618129 (3), Autosomal dominant |
| CARD11 | 84433 | ENSG00000198286 | B-cell expansion with NFKB and T-cell anergy, 616452 (3), Autosomal dominant; Immunodeficiency 11B with atopic dermatitis, 617638 (3), Autosomal dominant; Immunodeficiency 11A, 615206 (3), Autosomal recessive |
| CARTPT | 9607 | ENSG00000164326 | {?Obesity, susceptibility to}, 601665 (3), Multifactorial, Autosomal dominant, Autosomal recessive |
| CASP8 | 841 | ENSG00000064012 | syndrome, 607271 (3), Autosomal recessive; Hepatocellular carcinoma, somatic, 114550 (3); {Lung cancer, protection against}, 211980 (3), Somatic mutation, Autosomal dominant |
| CASR | 846 | ENSG00000036828 | 239200 (3), Autosomal dominant, Autosomal recessive; Hypocalcemia, autosomal dominant, 601198 (3), Autosomal dominant; Hypocalciuric hypercalcemia, type I, 145980 (3), Autosomal dominant; {?Epilepsy idiopathic generalized, susceptibility to, 8}, 612899 (3), Autosomal dominant |
| CAV1 | 857 | ENSG00000105974 | Lipodystrophy, congenital generalized, type 3, 612526 (3), Autosomal recessive; Pulmonary hypertension, primary, 3, 615343 (3), Autosomal dominant; Lipodystrophy, familial partial, type 7, 606721 (3), Autosomal dominant |
| CCDC88C | 440193 | ENSG00000015133 | ?Spinocerebellar ataxia 40, 616053 (3), Autosomal dominant; Hydrocephalus, congenital, 1, 236600 (3), Autosomal recessive |
| CD46 | 4179 | ENSG00000117335 | {Hemolytic uremic syndrome, atypical, susceptibility to, 2}, 612922 (3), Autosomal dominant, Autosomal recessive |
| CDH11 | 1009 | ENSG00000140937 | recessive |
| CDH2 | 1000 | ENSG00000170558 | 619957 (3), Autosomal recessive; Agenesis of corpus callosum, cardiac, ocular, and genital syndrome, 618929 (3), Autosomal dominant |
| CDH23 | 64072 | ENSG00000107736 | Usher syndrome, type 1D, 601067 (3), Digenic recessive, Autosomal recessive; {Pituitary adenoma 5, multiple types}, 617540 (3), Autosomal dominant; Usher syndrome, type 1D/F digenic, 601067 (3), Digenic recessive, Autosomal recessive; Deafness, autosomal recessive 12, 601386 (3), Autosomal recessive |
| CDSN | 1041 | ENSG00000204539 | Hypotrichosis 2, 146520 (3), Autosomal dominant; Peeling skin syndrome 1, 270300 (3), Autosomal recessive |
| CEACAM16 | 388551 | ENSG00000213892 | Autosomal recessive |
| CEBPE | 1053 | ENSG00000092067 | ?Immunodeficiency 108 with autoinflammation, 260570 (3), Autosomal recessive; Specific granule deficiency, 245480 (3), Autosomal dominant, Autosomal recessive |
| CFAP43 | 80217 | ENSG00000197748 | recessive |
| CFB | 629 | ENSG00000243649 | ?Complement factor B deficiency, 615561 (3), Autosomal recessive; {Hemolytic uremic syndrome, atypical, susceptibility to, 4}, 612924 (3), Autosomal dominant; {Macular degeneration, age-related, 14, reduced risk of}, 615489 (3), Digenic dominant |
| CFH | 3075 | ENSG00000000971 | {Macular degeneration, age-related, 4}, 610698 (3), Autosomal dominant; Basal laminar drusen, 126700 (3), Autosomal dominant; Complement factor H deficiency, 609814 (3), Autosomal dominant, Autosomal recessive; {Hemolytic uremic syndrome, atypical, susceptibility to, 1}, 235400 (3), Autosomal dominant, Autosomal recessive |
| CFHR1 | 3078 | ENSG00000244414 | {Macular degeneration, age-related, reduced risk of}, 603075 (3), Autosomal dominant; {Hemolytic uremic syndrome, atypical, susceptibility to}, 235400 (3), Autosomal dominant, Autosomal recessive |
| CFHR3 | 10878 | ENSG00000116785 | {Macular degeneration, age-related, reduced risk of}, 603075 (3), Autosomal dominant; {Hemolytic uremic syndrome, atypical, susceptibility to}, 235400 (3), Autosomal dominant, Autosomal recessive |
| CFI | 3426 | ENSG00000205403 | related, 13, susceptibility to}, 615439 (3), Autosomal dominant; Complement factor I deficiency, 610984 (3), Autosomal recessive |
| CFTR | 1080 | ENSG00000001626 | Cystic fibrosis, 219700 (3), Autosomal recessive; Sweat chloride elevation without CF (3); Congenital bilateral absence of vas deferens, 277180 (3), Autosomal recessive; {Pancreatitis, hereditary}, 167800 (3), Autosomal dominant; {Bronchiectasis with or without elevated sweat chloride 1, modifier of}, 211400 (3), Autosomal dominant; {Hypertrypsinemia, neonatal} (3) |

| | | | |
|---|---|---|---|
| CHRNA1 | 1134 | ENSG00000138435 | syndrome, congenital, 1A, slow-channel, 601462 (3), Autosomal dominant; Multiple pterygium syndrome, lethal type, 253290 (3), Autosomal recessive |
| CHRNB1 | 1140 | ENSG00000170175 | ?Myasthenic syndrome, congenital, 2C, associated with acetylcholine receptor deficiency, 616314 (3), Autosomal recessive; Myasthenic syndrome, congenital, 2A, slow-channel, 616313 (3), Autosomal dominant |
| CHRND | 1144 | ENSG00000135902 | Multiple pterygium syndrome, lethal type, 253290 (3), Autosomal recessive; Myasthenic syndrome, congenital, 3B, fast-channel, 616322 (3), Autosomal recessive; ?Myasthenic syndrome, congenital, 3A, slow-channel, 616321 (3), Autosomal dominant |
| CHRNE | 1145 | ENSG00000108556 | Myasthenic syndrome, congenital, 4A, slow-channel, 605809 (3), Autosomal dominant, Autosomal recessive; Myasthenic syndrome, congenital, 4C, associated with acetylcholine receptor deficiency, 608931 (3), Autosomal recessive; Myasthenic syndrome, congenital, 4B, fast-channel, 616324 (3), Autosomal recessive |
| CILK1 | 22858 | ENSG00000112144 | 612651 (3), Autosomal recessive |
| CLCN1 | 1180 | ENSG00000188037 | Myotonia congenita, recessive, 255700 (3), Autosomal recessive; Myotonia congenita, dominant, 160800 (3), Autosomal dominant; Myotonia levior, 160800 (3), Autosomal dominant |
| CLCN2 | 1181 | ENSG00000114859 | Autosomal dominant; {Epilepsy, juvenile myoclonic, susceptibility to, 8}, 607628 (3), Autosomal dominant; {Epilepsy, juvenile absence, susceptibility to, 2}, 607628 (3), Autosomal dominant; {Epilepsy, idiopathic generalized, susceptibility to, 11}, 607628 (3), Autosomal dominant |
| CLCN3 | 1182 | ENSG00000109572 | Neurodevelopmental disorder with seizures and brain abnormalities, 619517 (3), Autosomal recessive; Neurodevelopmental disorder with hypotonia and brain abnormalities, 619512 (3), Autosomal dominant |
| CLCN7 | 1186 | ENSG00000103249 | Osteopetrosis, autosomal recessive 4, 611490 (3), Autosomal recessive; Osteopetrosis, autosomal dominant 2, 166600 (3), Autosomal dominant |
| CLPB | 81570 | ENSG00000162129 | VIIB, autosomal recessive, 616271 (3), Autosomal recessive; 3-methylglutaconic aciduria, type VIIA, autosomal dominant, 619835 (3), Autosomal dominant |
| CNNM2 | 54805 | ENSG00000148842 | Hypomagnesemia 6, renal, 613882 (3), Autosomal dominant; Hypomagnesemia, seizures, and impaired intellectual development 1, 616418 (3), Autosomal dominant, Autosomal recessive |
| COCH | 1690 | ENSG00000100473 | Autosomal recessive |
| COG4 | 25839 | ENSG00000103051 | dominant |
| COL11A1 | 1301 | ENSG00000060718 | Marshall syndrome, 154780 (3), Autosomal dominant; Deafness, autosomal dominant 37, 618533 (3), Autosomal dominant; {Lumbar disc herniation, susceptibility to}, 603932 (3) |
| COL11A2 | 1302 | ENSG00000204248 | recessive, 215150 (3), Autosomal recessive; Fibrochondrogenesis 2, 614524 (3), Autosomal dominant, Autosomal recessive; Deafness, autosomal recessive 53, 609706 (3), Autosomal recessive; Otospondylomegaepiphyseal dysplasia, autosomal dominant, 184840 (3), Autosomal dominant |
| COL12A1 | 1303 | ENSG00000111799 | recessive |
| COL17A1 | 1308 | ENSG00000065618 | 619787 (3), Autosomal recessive |
| COL18A1 | 80781 | ENSG00000182871 | dominant |
| COL1A2 | 1278 | ENSG00000164692 | dominant; Ehlers-Danlos syndrome, arthrochalasia type, 2, 617821 (3), Autosomal dominant; Combined osteogenesis imperfecta and Ehlers-Danlos syndrome 2, 619120 (3), Autosomal dominant; Ehlers-Danlos syndrome, cardiac valvular type, 225320 (3), Autosomal recessive; Osteogenesis imperfecta, type IV, 166220 (3), Autosomal dominant; Osteogenesis imperfecta, type II, 166210 (3), Autosomal dominant |
| COL3A1 | 1281 | ENSG00000168542 | Ehlers-Danlos syndrome, vascular type, 130050 (3), Autosomal dominant; Polymicrogyria with or without vascular-type EDS, 618343 (3), Autosomal recessive |

| | | | |
|---|---|---|---|
| COL4A3 | 1285 | ENSG00000169031 | Alport syndrome 3A, autosomal dominant, 104200 (3), Autosomal dominant; Hematuria, benign familial, 2, 620320 (3); Alport syndrome 3B, autosomal recessive, 620536 (3) |
| COL4A4 | 1286 | ENSG00000081052 | Autosomal recessive |
| COL6A1 | 1291 | ENSG00000142156 | Bethlem myopathy 1, 158810 (3), Autosomal dominant, Autosomal recessive; Ullrich congenital muscular dystrophy 1, 254090 (3), Autosomal dominant, Autosomal recessive |
| COL6A2 | 1292 | ENSG00000142173 | Bethlem myopathy 1, 158810 (3), Autosomal dominant, Autosomal recessive; ?Myosclerosis, congenital, 255600 (3), Autosomal recessive; Ullrich congenital muscular dystrophy 1, 254090 (3), Autosomal dominant, Autosomal recessive |
| COL6A3 | 1293 | ENSG00000163359 | Ullrich congenital muscular dystrophy 1, 254090 (3), Autosomal dominant, Autosomal recessive; Dystonia 27, 616411 (3), Autosomal recessive; Bethlem myopathy 1, 158810 (3), Autosomal dominant, Autosomal recessive |
| COL7A1 | 1294 | ENSG00000114270 | 132000 (3), Autosomal dominant; Epidermolysis bullosa dystrophica inversa, 226600 (3), Autosomal recessive; Epidermolysis bullosa dystrophica, autosomal recessive, 226600 (3), Autosomal recessive; Epidermolysis bullosa, pretibial, 131850 (3), Autosomal dominant, Autosomal recessive; Epidermolysis bullosa dystrophica, autosomal dominant, 131750 (3), Autosomal dominant; Transient bullous of the newborn, 131705 (3), Autosomal dominant, Autosomal recessive; Epidermolysis bullosa pruriginosa, 604129 (3), Autosomal dominant, Autosomal recessive; Epidermolysis bullosa dystrophica, localisata variant, 226600 (3), Autosomal recessive |
| COL9A1 | 1297 | ENSG00000112280 | dominant |
| COL9A2 | 1298 | ENSG00000049089 | recessive |
| COL9A3 | 1299 | ENSG00000092758 | {Intervertebral disc disease, susceptibility to}, 603932 (3); Epiphyseal dysplasia, multiple, 3, with or without myopathy, 600969 (3), Autosomal dominant; Stickler syndrome, type VI, 620022 (3), Autosomal recessive |
| COPB2 | 9276 | ENSG00000184432 | Osteoporosis, childhood- or juvenile-onset, with developmental delay, 619884 (3), Autosomal dominant; ?Microcephaly 19, primary, autosomal recessive, 617800 (3), Autosomal recessive |
| COQ2 | 27235 | ENSG00000173085 | {Multiple system atrophy, susceptibility to}, 146500 (3), Autosomal dominant, Autosomal recessive; Coenzyme Q10 deficiency, primary, 1, 607426 (3), Autosomal recessive |
| CPA6 | 57094 | ENSG00000165078 | dominant, Autosomal recessive |
| CPOX | 1371 | ENSG00000080819 | Coproporphyria, 121300 (3), Autosomal dominant, Autosomal recessive; Harderoporphyria, 618892 (3), Autosomal recessive |
| CPT2 | 1376 | ENSG00000157184 | {Encephalopathy, acute, infection-induced, 4, susceptibility to}, 614212 (3), Autosomal dominant, Autosomal recessive; CPT II deficiency, infantile, 600649 (3), Autosomal recessive; CPT II deficiency, lethal neonatal, 608836 (3), Autosomal recessive; CPT II deficiency, myopathic, stress-induced, 255110 (3), Autosomal dominant, Autosomal recessive |
| CRB1 | 23418 | ENSG00000134376 | Leber congenital amaurosis 8, 613835 (3), Autosomal recessive; Retinitis pigmentosa-12, 600105 (3), Autosomal recessive; Pigmented paravenous chorioretinal atrophy, 172870 (3), Autosomal dominant |
| CRYAA | 1409 | ENSG00000160202 | Cataract 9, multiple types, 604219 (3), Autosomal dominant, Autosomal recessive |
| CRYAB | 1410 | ENSG00000109846 | Myopathy, myofibrillar, fatal infantile hypertonic, alpha-B crystallin-related, 613869 (3), Autosomal recessive; Myopathy, myofibrillar, 2, 608810 (3), Autosomal dominant; Cataract 16, multiple types, 613763 (3), Autosomal dominant, Autosomal recessive; Cardiomyopathy, dilated, 1II, 615184 (3), Autosomal dominant |
| CRYBB1 | 1414 | ENSG00000100122 | Cataract 17, multiple types, 611544 (3), Autosomal dominant, Autosomal recessive |
| CRYBB3 | 1417 | ENSG00000100053 | Cataract 22, 609741 (3), Autosomal dominant, Autosomal recessive |
| CSF1R | 1436 | ENSG00000182578 | Brain abnormalities, neurodegeneration, and dysosteosclerosis, 618476 (3), Autosomal recessive; Leukoencephalopathy, diffuse hereditary, with spheroids 1, 221820 (3), Autosomal dominant |
| CSF3R | 1441 | ENSG00000119535 | (3), Autosomal dominant |

| | | | |
|---|---|---|---|
| CYP11B1 | 1584 | ENSG00000160882 | Aldosteronism, glucocorticoid-remediable, 103900 (3), Autosomal dominant; Adrenal hyperplasia, congenital, due to 11-beta-hydroxylase deficiency, 202010 (3), Autosomal recessive |
| DCC | 1630 | ENSG00000187323 | somatic, 133239 (3); Colorectal cancer, somatic, 114500 (3); Gaze palsy, familial horizontal, with progressive scoliosis, 2, 617542 (3), Autosomal recessive |
| DCHS1 | 8642 | ENSG00000166341 | Mitral valve prolapse 2, 607829 (3), Autosomal dominant; Van Maldergem syndrome 1, 601390 (3), Autosomal recessive |
| DCTN1 | 1639 | ENSG00000204843 | dominant, Autosomal recessive; Neuronopathy, distal hereditary motor, autosomal dominant 14, 607641 (3), Autosomal dominant |
| DDR2 | 4921 | ENSG00000162733 | 271665 (3), Autosomal recessive |
| DEAF1 | 10522 | ENSG00000177030 | Vulto-van Silfout-de Vries syndrome, 615828 (3), Autosomal dominant; Neurodevelopmental disorder with hypotonia, impaired expressive language, and with or without seizures, 617171 (3), Autosomal recessive |
| DEPDC5 | 9681 | ENSG00000100150 | Epilepsy, familial focal, with variable foci 1, 604364 (3), Autosomal dominant; Developmental and epileptic encephalopathy 111, 620504 (3), Autosomal recessive |
| DES | 1674 | ENSG00000175084 | Scapuloperoneal syndrome, neurogenic, Kaeser type, 181400 (3), Autosomal dominant; Cardiomyopathy, dilated, 1I, 604765 (3), Autosomal dominant; Myopathy, myofibrillar, 1, 601419 (3), Autosomal dominant, Autosomal recessive |
| DHDDS | 79947 | ENSG00000117682 | disorder of glycosylation, type 1bb, 613861 (3), Autosomal recessive; Retinitis pigmentosa 59, 613861 (3), Autosomal recessive |
| DHTKD1 | 55526 | ENSG00000181192 | ?Charcot-Marie-Tooth disease, axonal, type 2Q, 615025 (3), Autosomal dominant; Alpha-aminoadipic and alpha-ketoadipic aciduria, 204750 (3), Autosomal recessive |
| DHX37 | 57647 | ENSG00000150990 | Neurodevelopmental disorder with brain anomalies and with or without vertebral or cardiac anomalies, 618731 (3), Autosomal recessive; 46XY sex reversal 11, 273250 (3), Autosomal dominant |
| DIAPH1 | 1729 | ENSG00000131504 | Deafness, autosomal dominant 1, with or without thrombocytopenia, 124900 (3), Autosomal dominant; Seizures, cortical blindness, microcephaly syndrome, 616632 (3), Autosomal recessive |
| DLX5 | 1749 | ENSG00000105880 | Split-hand/foot malformation 1, 183600 (3), Autosomal dominant; ?Split-hand/foot malformation 1 with sensorineural hearing loss, 220600 (3), Autosomal recessive |
| DMXL2 | 23312 | ENSG00000104093 | Developmental and epileptic encephalopathy 81, 618663 (3), Autosomal recessive; ?Deafness, autosomal dominant 71, 617605 (3), Autosomal dominant; ?Polyendocrine-polyneuropathy syndrome, 616113 (3), Autosomal recessive |
| DNA2 | 1763 | ENSG00000138346 | ?Seckel syndrome 8, 615807 (3), Autosomal recessive; Progressive external ophthalmoplegia with mitochondrial DNA deletions, autosomal dominant 6, 615156 (3), Autosomal dominant |
| DNAAF4 | 161582 | ENSG00000256061 | recessive |
| DNM1 | 1759 | ENSG00000106976 | Developmental and epileptic encephalopathy 31B, autosomal recessive, 620352 (3), Autosomal recessive; Developmental and epileptic encephalopathy 31A, autosomal dominant, 616346 (3), Autosomal dominant |
| DNM1L | 10059 | ENSG00000087470 | Optic atrophy 5, 610708 (3), Autosomal dominant; Encephalopathy, lethal, due to defective mitochondrial peroxisomal fission 1, 614388 (3), Autosomal dominant, Autosomal recessive |
| DNM2 | 1785 | ENSG00000079805 | Centronuclear myopathy 1, 160150 (3), Autosomal dominant; Charcot-Marie-Tooth disease, axonal type 2M, 606482 (3), Autosomal dominant; Charcot-Marie-Tooth disease, dominant intermediate B, 606482 (3), Autosomal dominant; Lethal congenital contracture syndrome 5, 615368 (3), Autosomal recessive |
| DSC2 | 1824 | ENSG00000134755 | dominant, Autosomal recessive; Arrhythmogenic right ventricular dysplasia 11, 610476 (3), Autosomal dominant, Autosomal recessive |
| DSG1 | 1828 | ENSG00000134760 | Keratosis palmoplantaris striata I, AD, 148700 (3), Autosomal dominant; Erythroderma, congenital, with palmoplantar keratoderma, hypotrichosis, and hyper IgE, 615508 (3), Autosomal recessive |
| DSG2 | 1829 | ENSG00000046604 | Autosomal dominant |

| | | | |
|---|---|---|---|
| DSP | 1832 | ENSG00000096696 | 609638 (3), Autosomal recessive; Keratosis palmoplantaris striata II, 612908 (3), Autosomal dominant; Dilated cardiomyopathy with woolly hair, keratoderma, and tooth agenesis, 615821 (3), Autosomal dominant; Cardiomyopathy, dilated, with woolly hair and keratoderma, 605676 (3), Autosomal recessive |
| DSTYK | 25778 | ENSG00000133059 | Spastic paraplegia 23, autosomal recessive, 270750 (3), Autosomal recessive; Congenital anomalies of kidney and urinary tract 1, 610805 (3), Autosomal dominant |
| DZIP1 | 22873 | ENSG00000134874 | Spermatogenic failure 47, 619102 (3), Autosomal recessive; ?Mitral valve prolapse 3, 610840 (3), Autosomal dominant |
| EDAR | 10913 | ENSG00000135960 | 129490 (3), Autosomal dominant; Ectodermal dysplasia 10B, hypohidrotic/hair/tooth type, autosomal recessive, 224900 (3), Autosomal recessive |
| EDARADD | 128178 | ENSG00000186197 | Ectodermal dysplasia 11B, hypohidrotic/hair/tooth type, autosomal recessive, 614941 (3), Autosomal recessive; Ectodermal dysplasia 11A, hypohidrotic/hair/tooth type, autosomal dominant, 614940 (3), Autosomal dominant |
| EDN1 | 1906 | ENSG00000078401 | recessive |
| EDN3 | 1908 | ENSG00000124205 | Waardenburg syndrome, type 4B, 613265 (3), Autosomal dominant, Autosomal recessive; {Hirschsprung disease, susceptibility to, 4}, 613712 (3), Autosomal dominant |
| EDNRB | 1910 | ENSG00000136160 | {Hirschsprung disease, susceptibility to, 2}, 600155 (3), Autosomal dominant; ?ABCD syndrome, 600501 (3), Autosomal recessive; Waardenburg syndrome, type 4A, 277580 (3), Autosomal dominant, Autosomal recessive |
| EGFR | 1956 | ENSG00000146648 | tyrosine kinase inhibitor in, 211980 (3), Somatic mutation, Autosomal dominant; Adenocarcinoma of lung, response to tyrosine kinase inhibitor in, 211980 (3), Somatic mutation, Autosomal dominant; {Nonsmall cell lung cancer, susceptibility to}, 211980 (3), Somatic mutation, Autosomal dominant |
| EGR2 | 1959 | ENSG00000122877 | 607678 (3), Autosomal dominant; Hypomyelinating neuropathy, congenital, 1, 605253 (3), Autosomal dominant, Autosomal recessive |
| EIF2AK2 | 5610 | ENSG00000055332 | Leukoencephalopathy, developmental delay, and episodic neurologic regression syndrome, 618877 (3), Autosomal dominant; Dystonia 33, 619687 (3), Autosomal dominant, Autosomal recessive |
| EIF4A2 | 1974 | ENSG00000156976 | Autosomal recessive |
| ELMOD3 | 84173 | ENSG00000115459 | Autosomal dominant |
| ELOVL1 | 64834 | ENSG00000066322 | recessive |
| ELOVL4 | 6785 | ENSG00000118402 | Spinocerebellar ataxia 34, 133190 (3), Autosomal dominant; Stargardt disease 3, 600110 (3), Autosomal dominant; Ichthyosis, spastic quadriplegia, and impaired intellectual development, 614457 (3), Autosomal recessive |
| ELP1 | 8518 | ENSG00000070061 | (3), Autosomal recessive |
| ENAM | 10117 | ENSG00000132464 | Autosomal dominant |
| ENPP1 | 5167 | ENSG00000197594 | autosomal recessive, 2, 613312 (3), Autosomal recessive; {Diabetes mellitus, non-insulin-dependent, susceptibility to}, 125853 (3), Autosomal dominant; Arterial calcification, generalized, of infancy, 1, 208000 (3), Autosomal recessive; Cole disease, 615522 (3), Autosomal dominant |
| EPB41 | 2035 | ENSG00000159023 | Elliptocytosis-1, 611804 (3), Autosomal dominant, Autosomal recessive |
| EPCAM | 4072 | ENSG00000119888 | dominant |
| EPHX2 | 2053 | ENSG00000120915 | {Hypercholesterolemia, familial, due to LDLR defect, modifier of}, 143890 (3), Autosomal dominant, Autosomal recessive |
| EPO | 2056 | ENSG00000130427 | {Microvascular complications of diabetes 2}, 612623 (3); Erythrocytosis, familial, 5, 617907 (3), Autosomal dominant; ?Diamond-Blackfan anemia-like, 617911 (3), Autosomal recessive |
| ERBB3 | 2065 | ENSG00000065361 | ?Lethal congenital contractural syndrome 2, 607598 (3), Autosomal recessive; {?Erythroleukemia, familial, susceptibility to}, 133180 (3), Autosomal dominant; Visceral neuropathy, familial, 1, autosomal recessive, 243180 (3), Autosomal recessive |

| | | | |
|---|---|---|---|
| ERCC6 | 2074 | ENSG00000225830 | UV-sensitive syndrome 1, 600630 (3), Autosomal recessive; Cerebrooculofacioskeletal syndrome 1, 214150 (3), Autosomal recessive; ?De Sanctis-Cacchione syndrome, 278800 (3), Autosomal recessive; Cockayne syndrome, type B, 133540 (3), Autosomal recessive; {Macular degeneration, age-related, susceptibility to, 5}, 613761 (3); Premature ovarian failure 11, 616946 (3), Autosomal dominant; {Lung cancer, susceptibility to}, 211980 (3), Somatic mutation, Autosomal dominant |
| ERLIN2 | 11160 | ENSG00000147475 | Spastic paraplegia 18A, autosomal dominant, 620512 (3), Autosomal dominant; Spastic paraplegia 18B, autosomal recessive, 611225 (3), Autosomal recessive |
| ESPN | 83715 | ENSG00000187017 | Deafness, neurosensory, without vestibular involvement, autosomal dominant, 609006 (3), Autosomal recessive; Deafness, autosomal recessive 36, 609006 (3), Autosomal recessive; ?Usher syndrome, type 1M, 618632 (3), Autosomal recessive |
| ESR1 | 2099 | ENSG00000091831 | Breast cancer, somatic, 114480 (3); {Migraine, susceptibility to}, 157300 (3), Autosomal dominant; Estrogen resistance, 615363 (3), Autosomal recessive; {Myocardial infarction, susceptibility to}, 608446 (3) |
| EVC | 2121 | ENSG00000072840 | dominant |
| EVC2 | 132884 | ENSG00000173040 | Ellis-van Creveld syndrome, 225500 (3), Autosomal recessive; Weyers acrofacial dysostosis, 193530 (3), Autosomal dominant |
| EXT2 | 2132 | ENSG00000151348 | Autosomal dominant |
| F11 | 2160 | ENSG00000088926 | Factor XI deficiency, autosomal dominant, 612416 (3); Factor XI deficiency, autosomal recessive, 612416 (3) |
| F12 | 2161 | ENSG00000131187 | Angioedema, hereditary, 3, 610618 (3), Autosomal dominant; Factor XII deficiency, 234000 (3), Autosomal recessive |
| F13A1 | 2162 | ENSG00000124491 | Factor XIIIA deficiency, 613225 (3), Autosomal recessive; {Myocardial infarction, protection against}, 608446 (3); {Venous thrombosis, protection against}, 188050 (3), Autosomal dominant |
| F2 | 2147 | ENSG00000180210 | Hypoprothrombinemia, 613679 (3), Autosomal recessive; {Pregnancy loss, recurrent, susceptibility to, 2}, 614390 (3), Autosomal dominant; Dysprothrombinemia, 613679 (3), Autosomal recessive; Thrombophilia 1 due to thrombin defect, 188050 (3), Autosomal dominant; {Stroke, ischemic, susceptibility to}, 601367 (3), Multifactorial |
| F5 | 2153 | ENSG00000198734 | susceptibility to, 1}, 614389 (3), Autosomal dominant; {Thrombophilia, susceptibility to, due to factor V Leiden}, 188055 (3), Autosomal dominant; {Budd-Chiari syndrome}, 600880 (3), Autosomal recessive; {Stroke, ischemic, susceptibility to}, 601367 (3), Multifactorial; Factor V deficiency, 227400 (3), Autosomal recessive |
| FAR1 | 84188 | ENSG00000197601 | Peroxisomal fatty acyl-CoA reductase 1 disorder, 616154 (3), Autosomal recessive; Cataracts, spastic paraparesis, and speech delay, 619338 (3), Autosomal dominant |
| FBLN5 | 10516 | ENSG00000140092 | 1H, 619764 (3), Autosomal dominant; Macular degeneration, age-related, 3, 608895 (3), Autosomal dominant; Neuropathy, hereditary, with or without age-related macular degeneration, 608895 (3), Autosomal dominant; ?Cutis laxa, autosomal dominant 2, 614434 (3), Autosomal dominant |
| FCGR2A | 2212 | ENSG00000143226 | {Malaria, severe, susceptibility to}, 611162 (3); {Pseudomonas aeruginosa, susceptibility to chronic infection by, in cystic fibrosis}, 219700 (3), Autosomal recessive; {Lupus nephritis, susceptibility to}, 152700 (3), Autosomal dominant |
| FGA | 2243 | ENSG00000171560 | Hypodysfibrinogenemia, congenital, 616004 (3); Dysfibrinogenemia, congenital, 616004 (3); Amyloidosis, familial visceral, 105200 (3), Autosomal dominant; Afibrinogenemia, congenital, 202400 (3), Autosomal recessive |
| FGF23 | 8074 | ENSG00000118972 | Tumoral calcinosis, hyperphosphatemic, familial, 2, 617993 (3), Autosomal recessive; Hypophosphatemic rickets, autosomal dominant, 193100 (3), Autosomal dominant |
| FGFR3 | 2261 | ENSG00000068078 | (3), Autosomal dominant; Thanatophoric dysplasia, type II, 187601 (3), Autosomal dominant; Nevus, epidermal, somatic, 162900 (3); CATSHL syndrome, 610474 (3), Autosomal dominant, Autosomal recessive; Thanatophoric dysplasia, type I, 187600 (3), Autosomal dominant; Spermatocytic seminoma, somatic, 273300 (3); Bladder cancer, somatic, 109800 (3); LADD syndrome 2, 620192 (3), Autosomal dominant; Achondroplasia, 100800 (3), Autosomal dominant; Cervical cancer, somatic, 603956 (3); Colorectal cancer, somatic, 114500 (3); Crouzon syndrome with acanthosis nigricans, 612247 (3), Autosomal |
| FH | 2271 | ENSG00000091483 | recessive |

| | | | |
|---|---|---|---|
| FIG4 | 9896 | ENSG00000112367 | recessive; Amyotrophic lateral sclerosis 11, 612577 (3), Autosomal dominant; Charcot-Marie-Tooth disease, type 4J, 611228 (3), Autosomal recessive |
| FLG | 2312 | ENSG00000143631 | Ichthyosis vulgaris, 146700 (3), Autosomal dominant, Autosomal recessive; {Dermatitis, atopic, susceptibility to, 2}, 605803 (3) |
| FLI1 | 2313 | ENSG00000151702 | Bleeding disorder, platelet-type, 21, 617443 (3), Autosomal dominant, Autosomal recessive |
| FLNB | 2317 | ENSG00000136068 | Atelosteogenesis, type III, 108721 (3), Autosomal dominant; Spondylocarpotarsal synostosis syndrome, 272460 (3), Autosomal recessive; Boomerang dysplasia, 112310 (3), Autosomal dominant |
| FOXE1 | 2304 | ENSG00000178919 | dominant |
| FOXE3 | 2301 | ENSG00000186790 | Anterior segment dysgenesis 2, multiple subtypes, 610256 (3), Autosomal recessive; {Aortic aneurysm, familial thoracic 11, susceptibility to}, 617349 (3), Autosomal dominant; Cataract 34, multiple types, 612968 (3) |
| FOXI3 | 344167 | ENSG00000214336 | Craniofacial microsomia 2, 620444 (3), Autosomal dominant, Autosomal recessive |
| FOXL2 | 668 | ENSG00000183770 | Blepharophimosis, epicanthus inversus, and ptosis, type 1, 110100 (3), Autosomal dominant, Autosomal recessive; Premature ovarian failure 3, 608996 (3), Autosomal dominant |
| FOXN1 | 8456 | ENSG00000109101 | T-cell lymphopenia, infantile, with or without nail dystrophy, autosomal dominant, 618806 (3), Autosomal dominant; T-cell immunodeficiency, congenital alopecia, and nail dystrophy, 601705 (3), Autosomal recessive |
| FREM1 | 158326 | ENSG00000164946 | Manitoba oculotrichoanal syndrome, 248450 (3), Autosomal recessive; Bifid nose with or without anorectal and renal anomalies, 608980 (3), Autosomal recessive; Trigonocephaly 2, 614485 (3), Autosomal dominant |
| FSHR | 2492 | ENSG00000170820 | Ovarian response to FSH stimulation, 276400 (3), Autosomal recessive; Ovarian hyperstimulation syndrome, 608115 (3), Autosomal dominant; Ovarian dysgenesis 1, 233300 (3), Autosomal recessive |
| FTL | 2512 | ENSG00000087086 | (3), Autosomal dominant, Autosomal recessive; Neurodegeneration with brain iron accumulation 3, 606159 (3), Autosomal dominant |
| GATA5 | 140628 | ENSG00000130700 | Congenital heart defects, multiple types, 5, 617912 (3), Autosomal dominant, Autosomal recessive |
| GATM | 2628 | ENSG00000171766 | Autosomal dominant |
| GBA1 | 2629 | ENSG00000177628 | {Lewy body dementia, susceptibility to}, 127750 (3), Autosomal dominant; Gaucher disease, type II, 230900 (3), Autosomal recessive; Gaucher disease, type IIIC, 231005 (3), Autosomal recessive; Gaucher disease, type III, 231000 (3), Autosomal recessive; Gaucher disease, type I, 230800 (3), Autosomal recessive; Gaucher disease, perinatal lethal, 608013 (3), Autosomal recessive; {Parkinson disease, late-onset, susceptibility to}, 168600 (3), Multifactorial, Autosomal dominant |
| GCH1 | 2643 | ENSG00000131979 | Dystonia, DOPA-responsive, 128230 (3), Autosomal dominant, Autosomal recessive; Hyperphenylalaninemia, BH4-deficient, B, 233910 (3), Autosomal recessive |
| GCK | 2645 | ENSG00000106633 | Hyperinsulinemic hypoglycemia, familial, 3, 602485 (3), Autosomal dominant; Diabetes mellitus, noninsulin-dependent, late onset, 125853 (3), Autosomal dominant |
| GCM2 | 9247 | ENSG00000124827 | 617343 (3), Autosomal dominant |
| GCNT2 | 2651 | ENSG00000111846 | [Blood group, Ii], 110800 (3), Autosomal dominant; Adult i phenotype without cataract, 110800 (3), Autosomal dominant; Cataract 13 with adult i phenotype, 116700 (3), Autosomal recessive |
| GDAP1 | 54332 | ENSG00000104381 | Charcot-Marie-Tooth disease, axonal, with vocal cord paresis, 607706 (3), Autosomal recessive; Charcot-Marie-Tooth disease, recessive intermediate, A, 608340 (3), Autosomal recessive; Charcot-Marie-Tooth disease, axonal, type 2K, 607831 (3), Autosomal dominant, Autosomal recessive; Charcot-Marie-Tooth disease, type 4A, 214400 (3), Autosomal recessive |
| GDF1 | 2657 | ENSG00000130283 | Autosomal recessive |
| GDF5 | 8200 | ENSG00000125965 | recessive; Multiple synostoses syndrome 2, 610017 (3), Autosomal dominant; Symphalangism, proximal, 1B, 615298 (3), Autosomal dominant; Brachydactyly, type A2, 112600 (3), Autosomal dominant; ?Acromesomelic dysplasia 2C, Hunter-Thompson type, 201250 (3), Autosomal recessive; Brachydactyly, type C, 113100 (3), Autosomal dominant; {Osteoarthritis-5}, 612400 (3); Brachydactyly, type A1, C, 615072 (3), Autosomal dominant, Autosomal recessive |

| | | | |
|---|---|---|---|
| GDF6 | 392255 | ENSG00000156466 | Microphthalmia with coloboma 6, digenic, 613703 (3), Autosomal dominant; Microphthalmia, isolated 4, 613094 (3); Leber congenital amaurosis 17, 615360 (3), Autosomal recessive; Multiple synostoses syndrome 4, 617898 (3), Autosomal dominant; Klippel-Feil syndrome 1, autosomal dominant, 118100 (3), Autosomal dominant |
| GFI1B | 8328 | ENSG00000165702 | Bleeding disorder, platelet-type, 17, 187900 (3), Autosomal dominant, Autosomal recessive |
| GH1 | 2688 | ENSG00000259384 | dominant; Growth hormone deficiency, isolated, type IB, 612781 (3); Growth hormone deficiency, isolated, type IA, 262400 (3), Autosomal recessive |
| GHR | 2690 | ENSG00000112964 | dominant; Growth hormone insensitivity, partial, 604271 (3), Autosomal dominant; {Hypercholesterolemia, familial, modifier of}, 143890 (3), Autosomal dominant, Autosomal recessive |
| GHRL | 51738 | ENSG00000157017 | {Obesity, susceptibility to}, 601665 (3), Multifactorial, Autosomal dominant, Autosomal recessive |
| GHSR | 2693 | ENSG00000121853 | Growth hormone deficiency, isolated partial, 615925 (3), Autosomal dominant, Autosomal recessive |
| GJA1 | 2697 | ENSG00000152661 | Erythrokeratodermia variabilis et progressiva 3, 617525 (3), Autosomal dominant; Craniometaphyseal dysplasia, autosomal recessive, 218400 (3), Autosomal recessive; Oculodentodigital dysplasia, 164200 (3), Autosomal dominant; Palmoplantar keratoderma with congenital alopecia, 104100 (3), Autosomal dominant; Syndactyly, type III, 186100 (3), Autosomal dominant; Oculodentodigital dysplasia, autosomal recessive, 257850 (3), Autosomal recessive |
| GJB2 | 2706 | ENSG00000165474 | Digenic dominant, Autosomal recessive; Deafness, autosomal dominant 3A, 601544 (3), Autosomal dominant; Hystrix-like ichthyosis with deafness, 602540 (3), Autosomal dominant; Bart-Pumphrey syndrome, 149200 (3), Autosomal dominant; Keratitis-ichthyosis-deafness syndrome, 148210 (3), Autosomal dominant; Vohwinkel syndrome, 124500 (3), Autosomal dominant |
| GJB3 | 2707 | ENSG00000188910 | Deafness, digenic, GJB2/GJB3, 220290 (3), Digenic dominant, Autosomal recessive; Deafness, autosomal recessive (3); Deafness, autosomal dominant 2B, 612644 (3), Autosomal dominant; Erythrokeratodermia variabilis et progressiva 1, 133200 (3), Autosomal dominant, Autosomal recessive; Deafness, autosomal dominant, with peripheral neuropathy (3) |
| GJB6 | 10804 | ENSG00000121742 | Autosomal dominant; Deafness, autosomal recessive 1B, 612645 (3), Autosomal recessive; Deafness, digenic GJB2/GJB6, 220290 (3), Digenic dominant, Autosomal recessive |
| GJC2 | 57165 | ENSG00000198835 | Lymphatic malformation 3, 613480 (3), Autosomal dominant; ?Spastic paraplegia 44, autosomal recessive, 613206 (3), Autosomal recessive; Leukodystrophy, hypomyelinating, 2, 608804 (3), Autosomal recessive |
| GLRA1 | 2741 | ENSG00000145888 | Hyperekplexia 1, 149400 (3), Autosomal dominant, Autosomal recessive |
| GLS | 2744 | ENSG00000115419 | Global developmental delay, progressive ataxia, and elevated glutamine, 618412 (3), Autosomal recessive; ?Infantile cataract, skin abnormalities, glutamate excess, and impaired intellectual development, 618339 (3), Autosomal dominant; Developmental and epileptic encephalopathy 71, 618328 (3), Autosomal recessive |
| GNAT1 | 2779 | ENSG00000114349 | Night blindness, congenital stationary, autosomal dominant 3, 610444 (3), Autosomal dominant; Night blindness, congenital stationary, type 1G, 616389 (3), Autosomal recessive |
| GNE | 10020 | ENSG00000159921 | Sialuria, 269921 (3), Autosomal dominant; Nonaka myopathy, 605820 (3), Autosomal recessive |
| GP1BA | 2811 | ENSG00000185245 | Bernard-Soulier syndrome, type A1 (recessive), 231200 (3), Autosomal recessive; Bernard-Soulier syndrome, type A2 (dominant), 153670 (3), Autosomal dominant; von Willebrand disease, platelet-type, 177820 (3), Autosomal dominant; {Nonarteritic anterior ischemic optic neuropathy, susceptibility to}, 258660 (3), Autosomal recessive |
| GPR161 | 23432 | ENSG00000143147 | {Medulloblastoma predisposition syndrome}, 155255 (3), Somatic mutation, Autosomal dominant, Autosomal recessive |
| GRHL2 | 79977 | ENSG00000083307 | Deafness, autosomal dominant 28, 608641 (3), Autosomal dominant; Ectodermal dysplasia/short stature syndrome, 616029 (3), Autosomal recessive; Corneal dystrophy, posterior polymorphous, 4, 618031 (3), Autosomal dominant |
| GRIA1 | 2890 | ENSG00000155511 | ?Intellectual developmental disorder, autosomal recessive 76, 619931 (3), Autosomal recessive; Intellectual developmental disorder, autosomal dominant 67, 619927 (3), Autosomal dominant |

| | | | |
|---|---|---|---|
| GRIK2 | 2898 | ENSG00000164418 | Neurodevelopmental disorder with impaired language and ataxia and with or without seizures, 619580 (3), Autosomal dominant; Intellectual developmental disorder, autosomal recessive 6, 611092 (3), Autosomal recessive |
| GRIN1 | 2902 | ENSG00000176884 | Autosomal recessive; Developmental and epileptic encephalopathy 101, 619814 (3), Autosomal recessive; Neurodevelopmental disorder with or without hyperkinetic movements and seizures, autosomal dominant, 614254 (3), Autosomal dominant |
| GRM1 | 2911 | ENSG00000152822 | Autosomal dominant |
| GRN | 2896 | ENSG00000030582 | Aphasia, primary progressive, 607485 (3), Autosomal dominant; Frontotemporal lobar degeneration with ubiquitin-positive inclusions, 607485 (3), Autosomal dominant; Ceroid lipofuscinosis, neuronal, 11, 614706 (3), Autosomal recessive |
| GUCY2C | 2984 | ENSG00000070019 | Diarrhea 6, 614616 (3), Autosomal dominant; Meconium ileus, 614665 (3), Autosomal recessive |
| GUCY2D | 3000 | ENSG00000132518 | (3), Autosomal dominant; Leber congenital amaurosis 1, 204000 (3), Autosomal recessive; Night blindness, congenital stationary, type 1I, 618555 (3), Autosomal recessive |
| HAL | 3034 | ENSG00000084110 | [Histidinemia], 235800 (3), Autosomal dominant, Autosomal recessive |
| HARS1 | 3035 | ENSG00000170445 | Autosomal recessive |
| HBB | 3043 | ENSG00000244734 | Autosomal dominant; Sickle cell disease, 603903 (3), Autosomal recessive; Thalassemia, beta, 613985 (3); Delta-beta thalassemia, 141749 (3), Autosomal dominant; {Malaria, resistance to}, 611162 (3); Hereditary persistence of fetal hemoglobin, 141749 (3), Autosomal dominant; Erythrocytosis, familial, 6, 617980 (3), Autosomal dominant; Heinz body anemia, 140700 (3), Autosomal dominant |
| HEPACAM | 220296 | ENSG00000165478 | leukoencephalopathy with subcortical cysts 2B, remitting, with or without impaired intellectual development, 613926 (3), Autosomal dominant |
| HESX1 | 8820 | ENSG00000163666 | 182230 (3), Autosomal dominant, Autosomal recessive; Growth hormone deficiency with pituitary anomalies, 182230 (3), Autosomal dominant, Autosomal recessive |
| HK1 | 3098 | ENSG00000156515 | Retinitis pigmentosa 79, 617460 (3), Autosomal dominant; Neuropathy, hereditary motor and sensory, Russe type, 605285 (3), Autosomal recessive; Neurodevelopmental disorder with visual defects and brain anomalies, 618547 (3), Autosomal dominant; Hemolytic anemia due to hexokinase deficiency, 235700 (3), Autosomal recessive |
| HLA-DQB1 | 3119 | ENSG00000179344 | {Celiac disease, susceptibility to}, 212750 (3), Multifactorial, Autosomal recessive; {Multiple sclerosis, susceptibility to, 1}, 126200 (3), Multifactorial; {Creutzfeldt-Jakob disease, variant, resistance to}, 123400 (3), Autosomal dominant |
| HNF1A | 6927 | ENSG00000135100 | Hepatic adenoma, somatic, 142330 (3); Diabetes mellitus, insulin-dependent, 20, 612520 (3); {Diabetes mellitus, noninsulin-dependent, 2}, 125853 (3), Autosomal dominant; MODY, type III, 600496 (3), Autosomal dominant; {Diabetes mellitus, insulin-dependent}, 222100 (3), Autosomal recessive; Renal cell carcinoma, 144700 (3) |
| HNMT | 3176 | ENSG00000150540 | 600807 (3), Autosomal dominant |
| HOXA2 | 3199 | ENSG00000105996 | Microtia with or without hearing impairment (AD), 612290 (3), Autosomal dominant, Autosomal recessive; ?Microtia, hearing impairment, and cleft palate (AR), 612290 (3), Autosomal dominant, Autosomal recessive |
| HPD | 3242 | ENSG00000158104 | Hawkinsinuria, 140350 (3), Autosomal dominant; Tyrosinemia, type III, 276710 (3), Autosomal recessive |
| HSPA9 | 3313 | ENSG00000113013 | Even-plus syndrome, 616854 (3), Autosomal recessive; Anemia, sideroblastic, 4, 182170 (3), Autosomal dominant |
| HSPD1 | 3329 | ENSG00000144381 | (3), Autosomal recessive |
| HTRA1 | 5654 | ENSG00000166033 | {Macular degeneration, age-related, neovascular type}, 610149 (3); {Macular degeneration, age-related, 7}, 610149 (3); CARASIL syndrome, 600142 (3), Autosomal recessive; Cerebral arteriopathy, autosomal dominant, with subcortical infarcts and leukoencephalopathy, type 2, 616779 (3), Autosomal dominant |
| HTT | 3064 | ENSG00000197386 | Lopes-Maciel-Rodan syndrome, 617435 (3), Autosomal recessive; Huntington disease, 143100 (3), Autosomal dominant |

| | | | |
|---|---|---|---|
| IFIH1 | 64135 | ENSG00000115267 | Immunodeficiency 95, 619773 (3), Autosomal recessive; Aicardi-Goutieres syndrome 7, 615846 (3), Autosomal dominant; Singleton-Merten syndrome 1, 182250 (3), Autosomal dominant |
| IFNG | 3458 | ENSG00000111537 | {Hepatitis C virus, response to therapy of}, 609532 (3); {TSC2 angiomyolipomas, renal, modifier of}, 613254 (3), Autosomal dominant; {Aplastic anemia}, 609135 (3); ?Immunodeficiency 69, mycobacteriosis, 618963 (3), Autosomal recessive; {Tuberculosis, protection against}, 607948 (3); {AIDS, rapid progression to}, 609423 (3) |
| IFNGR1 | 3459 | ENSG00000027697 | {H. pylori infection, susceptibility to}, 600263 (3); Immunodeficiency 27A, mycobacteriosis, AR, 209950 (3), Autosomal recessive; Immunodeficiency 27B, mycobacteriosis, AD, 615978 (3), Autosomal dominant; {Tuberculosis infection, protection against}, 607948 (3); {Tuberculosis, susceptibility to}, 607948 (3); {Hepatitis B virus infection, susceptibility to}, 610424 (3) |
| IGF1R | 3480 | ENSG00000140443 | Insulin-like growth factor I, resistance to, 270450 (3), Autosomal dominant, Autosomal recessive |
| IHH | 3549 | ENSG00000163501 | Acrocapitofemoral dysplasia, 607778 (3), Autosomal recessive; Brachydactyly, type A1, 112500 (3), Autosomal dominant |
| IKBKB | 3551 | ENSG00000104365 | Immunodeficiency 15B, 615592 (3), Autosomal recessive; Immunodeficiency 15A, 618204 (3), Autosomal dominant |
| IL17RD | 54756 | ENSG00000144730 | recessive |
| IL6 | 3569 | ENSG00000136244 | hemorrhage in brain cerebrovascular malformations, susceptibility to}, 108010 (3), Somatic mutation; {Type 1 diabetes mellitus}, 222100 (3), Autosomal recessive; {Crohn disease-associated growth failure}, 266600 (3), Multifactorial; {Kaposi sarcoma, susceptibility to}, 148000 (3), Autosomal dominant |
| IL6ST | 3572 | ENSG00000134352 | syndrome 2, 619751 (3), Autosomal recessive; Hyper-IgE syndrome 4B, autosomal recessive, with recurrent infections, 618523 (3), Autosomal recessive; ?Immunodeficiency 94 with autoinflammation and dysmorphic facies, 619750 (3), Autosomal dominant |
| IMPG1 | 3617 | ENSG00000112706 | Autosomal dominant |
| IMPG2 | 50939 | ENSG00000081148 | Retinitis pigmentosa 56, 613581 (3), Autosomal recessive; Macular dystrophy, vitelliform, 5, 616152 (3), Autosomal dominant |
| INS | 3630 | ENSG00000254647 | 613370 (3), Autosomal dominant; Hyperproinsulinemia, 616214 (3), Autosomal dominant; Diabetes mellitus, permanent neonatal 4, 618858 (3), Autosomal dominant, Autosomal recessive |
| INSR | 3643 | ENSG00000171105 | nigricans, 610549 (3); Donohue syndrome, 246200 (3), Autosomal recessive; Hyperinsulinemic hypoglycemia, familial, 5, 609968 (3), Autosomal dominant |
| IRF8 | 3394 | ENSG00000140968 | Immunodeficiency 32A, mycobacteriosis, autosomal dominant, 614893 (3), Autosomal dominant; Immunodeficiency 32B, monocyte and dendritic cell deficiency, autosomal recessive, 226990 (3), Autosomal recessive |
| ITGA2B | 3674 | ENSG00000005961 | Thrombocytopenia, neonatal alloimmune, BAK antigen related (3); Glanzmann thrombasthenia 1, 273800 (3), Autosomal recessive; Bleeding disorder, platelet-type, 16, autosomal dominant, 187800 (3), Autosomal dominant |
| ITGB3 | 3690 | ENSG00000259207 | posttransfusion (3); {Myocardial infarction, susceptibility to}, 608446 (3); Glanzmann thrombasthenia 2, 619267 (3), Autosomal recessive |
| ITPR1 | 3708 | ENSG00000150995 | Gillespie syndrome, 206700 (3), Autosomal dominant, Autosomal recessive; Spinocerebellar ataxia 29, congenital nonprogressive, 117360 (3), Autosomal dominant; Spinocerebellar ataxia 15, 606658 (3), Autosomal dominant |
| ITPR3 | 3710 | ENSG00000096433 | Charcot-Marie-Tooth disease, demyelinating, type 1J, 620111 (3), Autosomal dominant; {Diabetes, type 1, susceptibility to}, 222100 (2), Autosomal recessive |
| JPH1 | 56704 | ENSG00000104369 | ?Charcot-Marie-Tooth disease, axonal, autosomal dominant, type 2K, 607831 (3), Autosomal dominant, Autosomal recessive |
| JPH2 | 57158 | ENSG00000149596 | dominant |
| JUP | 3728 | ENSG00000173801 | dominant |
| KCNE1 | 3753 | ENSG00000180509 | dominant |

| | | | |
|---|---|---|---|
| KCNJ11 | 3767 | ENSG00000187486 | 2, susceptibility to}, 125853 (3), Autosomal dominant; Maturity-onset diabetes of the young, type 13, 616329 (3), Autosomal dominant; Diabetes mellitus, transient neonatal 3, 610582 (3), Autosomal dominant; Hyperinsulinemic hypoglycemia, familial, 2, 601820 (3), Autosomal dominant, Autosomal recessive |
| KCNJ13 | 3769 | ENSG00000115474 | [illegible] recessive |
| KCNMA1 | 3778 | ENSG00000156113 | {Epilepsy, idiopathic generalized, susceptibility to, 16}, 618596 (3), Autosomal dominant; Paroxysmal nonkinesigenic dyskinesia, 3, with or without generalized epilepsy, 609446 (3), Autosomal dominant; Cerebellar atrophy, developmental delay, and seizures, 617643 (3), Autosomal recessive; Liang-Wang syndrome, 618729 (3), Autosomal dominant |
| KCNQ1 | 3784 | ENSG00000053918 | Short QT syndrome 2, 609621 (3), Autosomal dominant; Atrial fibrillation, familial, 3, 607554 (3), Autosomal dominant; Long QT syndrome 1, 192500 (3), Autosomal dominant; {Long QT syndrome 1, acquired, susceptibility to}, 192500 (3), Autosomal dominant; Jervell and Lange-Nielsen syndrome, 220400 (3), Autosomal recessive |
| KIDINS220 | 57498 | ENSG00000134313 | Spastic paraplegia, intellectual disability, nystagmus, and obesity, 617296 (3), Autosomal dominant; Ventriculomegaly and arthrogryposis, 619501 (3), Autosomal recessive |
| KIF1A | 547 | ENSG00000130294 | NESCAV syndrome, 614255 (3), Autosomal dominant; Neuropathy, hereditary sensory, type IIC, 614213 (3), Autosomal recessive; Spastic paraplegia 30, autosomal dominant, 610357 (3), Autosomal dominant, Autosomal recessive; Spastic paraplegia 30, autosomal recessive, 610357 (3), Autosomal dominant, Autosomal recessive |
| KISS1R | 84634 | ENSG00000116014 | Hypogonadotropic hypogonadism 8 with or without anosmia, 614837 (3), Autosomal recessive; ?Precocious puberty, central, 1, 176400 (3), Autosomal dominant |
| KITLG | 4254 | ENSG00000049130 | 619947 (3), Autosomal recessive; Deafness, autosomal dominant 69, unilateral or asymmetric, 616697 (3), Autosomal dominant; [Skin/hair/eye pigmentation 7, blond/brown hair], 611664 (3) |
| KLHL24 | 54800 | ENSG00000114796 | Cardiomyopathy, familial hypertrophic, 29, with polyglucosan bodies, 620236 (3), Autosomal recessive; Epidermolysis bullosa simplex 6, generalized intermediate, with or without cardiomyopathy, 617294 (3), Autosomal dominant |
| KLHL3 | 26249 | ENSG00000146021 | Pseudohypoaldosteronism, type IID, 614495 (3), Autosomal dominant, Autosomal recessive |
| KLHL7 | 55975 | ENSG00000122550 | Retinitis pigmentosa 42, 612943 (3), Autosomal dominant; PERCHING syndrome, 617055 (3), Autosomal recessive |
| KNG1 | 3827 | ENSG00000113889 | [Kininogen deficiency], 228960 (3), Autosomal recessive; Angioedema, hereditary, 6, 619363 (3), Autosomal dominant; [High molecular weight kininogen deficiency], 228960 (3), Autosomal recessive |
| KRT10 | 3858 | ENSG00000186395 | dominant, Autosomal recessive; ?Ichthyosis histrix, Lambert type, 146600 (3), Autosomal dominant; Ichthyosis with confetti, 609165 (3), Autosomal dominant |
| KRT14 | 3861 | ENSG00000186847 | recessive; Epidermolysis bullosa simplex 1C, localized, 131800 (3), Autosomal dominant; Dermatopathia pigmentosa reticularis, 125595 (3), Autosomal dominant; Epidermolysis bullosa simplex 1A, generalized severe, 131760 (3), Autosomal dominant; Naegeli-Franceschetti-Jadassohn syndrome, 161000 (3), Autosomal dominant; Epidermolysis bullosa simplex 1B, generalized intermediate, 131900 (3), Autosomal dominant |
| KRT5 | 3852 | ENSG00000186081 | [illegible] (3), Autosomal dominant; Epidermolysis bullosa simplex 2F, with mottled pigmentation, 131960 (3), Autosomal dominant; Epidermolysis bullosa simplex 2D, generalized, intermediate or severe, autosomal recessive, 619599 (3), Autosomal recessive; Epidermolysis bullosa simplex 2B, generalized intermediate, 619588 (3), Autosomal dominant; Epidermolysis bullosa simplex 2C, localized, 619594 (3), Autosomal dominant; Epidermolysis bullosa simplex 2E, with migratory circinate erythema, 609352 (3), Autosomal dominant |
| KRT74 | 121391 | ENSG00000170484 | Woolly hair, autosomal dominant, 194300 (3), Autosomal dominant; ?Hypotrichosis 3, 613981 (3), Autosomal dominant; ?Ectodermal dysplasia 7, hair/nail type, 614929 (3), Autosomal recessive |
| KRT83 | 3889 | ENSG00000170523 | Monilethrix, 158000 (3), Autosomal dominant; Erythrokeratodermia variabilis et progressiva 5, 617756 (3), Autosomal recessive |
| LAMB3 | 3914 | ENSG00000196878 | Epidermolysis bullosa, junctional 1B, severe, 226700 (3), Autosomal recessive; Epidermolysis bullosa, junctional 1A, intermediate, 226650 (3), Autosomal recessive; Amelogenesis imperfecta, type IA, 104530 (3), Autosomal dominant |

| | | | |
|---|---|---|---|
| LBR | 3930 | ENSG00000143815 | skeletal dysplasia with or without Pelger-Huet anomaly, 618019 (3), Autosomal recessive; Greenberg skeletal dysplasia, 215140 (3), Autosomal recessive |
| LDLR | 3949 | ENSG00000130164 | LDL cholesterol level QTL2, 143890 (3), Autosomal dominant, Autosomal recessive; Hypercholesterolemia, familial, 1, 143890 (3), Autosomal dominant, Autosomal recessive |
| LEMD2 | 221496 | ENSG00000161904 | [illegible] recessive |
| LHCGR | 3973 | ENSG00000138039 | 238320 (3), Autosomal recessive; Leydig cell hypoplasia with hypergonadotropic hypogonadism, 238320 (3), Autosomal recessive; Luteinizing hormone resistance, female, 238320 (3), Autosomal recessive; Precocious puberty, male, 176410 (3), Autosomal dominant |
| LIM2 | 3982 | ENSG00000105370 | Cataract 19, multiple types, 615277 (3), Autosomal dominant, Autosomal recessive |
| LIPC | 3990 | ENSG00000166035 | {Diabetes mellitus, noninsulin-dependent}, 125853 (3), Autosomal dominant; Hepatic lipase deficiency, 614025 (3), Autosomal recessive; [High density lipoprotein cholesterol level QTL 12], 612797 (3) |
| LMAN2L | 81562 | ENSG00000114988 | ?Intellectual developmental disorder, autosomal dominant 69, 617863 (3); ?Intellectual developmental disorder, autosomal recessive 52, 616887 (3), Autosomal recessive |
| LMBR1 | 64327 | ENSG00000105983 | Triphalangeal thumb, type I, 174500 (3), Autosomal dominant; Syndactyly, type IV, 186200 (3), Autosomal dominant; Laurin-Sandrow syndrome, 135750 (3), Autosomal dominant; Hypoplastic or aplastic tibia with polydactyly, 188740 (3), Autosomal dominant, Autosomal recessive; Polydactyly, preaxial type II, 174500 (3), Autosomal dominant; Acheiropody, 200500 (3), Autosomal recessive; Triphalangeal thumb-polysyndactyly syndrome, 190605 (3), Autosomal dominant |
| LMNA | 4000 | ENSG00000160789 | dominant; Cardiomyopathy, dilated, 1A, 115200 (3), Autosomal dominant; Emery-Dreifuss muscular dystrophy 3, autosomal recessive, 616516 (3), Autosomal recessive; Restrictive dermopathy 2, 619793 (3); Charcot-Marie-Tooth disease, type 2B1, 605588 (3), Autosomal recessive; Emery-Dreifuss muscular dystrophy 2, autosomal dominant, 181350 (3), Autosomal dominant; Hutchinson-Gilford progeria, 176670 (3), Autosomal dominant; Lipodystrophy, familial partial, type 2, 151660 (3), Autosomal dominant; Muscular dystrophy, congenital, 613205 (3), Autosomal dominant; Malouf syndrome, 212112 (3), |
| LMNB2 | 84823 | ENSG00000176619 | Microcephaly 27, primary, autosomal dominant, 619180 (3), Autosomal dominant; ?Epilepsy, progressive myoclonic, 9, 616540 (3), Autosomal recessive; {Lipodystrophy, partial, acquired, susceptibility to}, 608709 (3), Autosomal dominant |
| LPL | 4023 | ENSG00000175445 | Lipoprotein lipase deficiency, 238600 (3), Autosomal recessive; [High density lipoprotein cholesterol level QTL 11], 238600 (3), Autosomal recessive; Combined hyperlipidemia, familial, 144250 (3), Autosomal dominant |
| LRP4 | 4038 | ENSG00000134569 | ?Myasthenic syndrome, congenital, 17, 616304 (3), Autosomal recessive; Sclerosteosis 2, 614305 (3), Autosomal dominant, Autosomal recessive; Cenani-Lenz syndactyly syndrome, 212780 (3), Autosomal recessive |
| LRP5 | 4041 | ENSG00000162337 | Autosomal dominant; Polycystic liver disease 4 with or without kidney cysts, 617875 (3), Autosomal dominant; Endosteal hyperostosis, 144750 (3), Autosomal dominant; Osteoporosis-pseudoglioma syndrome, 259770 (3), Autosomal recessive; Exudative vitreoretinopathy 4, 601813 (3), Autosomal dominant, Autosomal recessive |
| LRSAM1 | 90678 | ENSG00000148356 | Charcot-Marie-Tooth disease, axonal, type 2P, 614436 (3), Autosomal dominant, Autosomal recessive |
| LTBP3 | 4054 | ENSG00000168056 | [illegible] dominant |
| LZTR1 | 8216 | ENSG00000099949 | Noonan syndrome 2, 605275 (3), Autosomal recessive; Noonan syndrome 10, 616564 (3), Autosomal dominant; {Schwannomatosis-2, susceptibility to}, 615670 (3), Autosomal dominant |
| MAB21L2 | 10586 | ENSG00000181541 | Microphthalmia/coloboma and skeletal dysplasia syndrome, 615877 (3), Autosomal dominant, Autosomal recessive |
| MAPT | 4137 | ENSG00000186868 | Supranuclear palsy, progressive, 601104 (3), Autosomal dominant; Supranuclear palsy, progressive atypical, 260540 (3), Autosomal recessive; Dementia, frontotemporal, with or without parkinsonism, 600274 (3), Autosomal dominant; {Parkinson disease, susceptibility to}, 168600 (3), Multifactorial, Autosomal dominant; Pick disease, 172700 (3), Autosomal dominant |
| MARS1 | 4141 | ENSG00000166986 | Spastic paraplegia 70, autosomal recessive, 620323 (3), Autosomal recessive; Interstitial lung and liver disease, 615486 (3), Autosomal recessive; ?Trichothiodystrophy 9, nonphotosensitive, 619692 (3), Autosomal recessive; Charcot-Marie-Tooth disease, axonal, type 2U, 616280 (3), Autosomal dominant |

| | | | |
|---|---|---|---|
| MAT1A | 4143 | ENSG00000151224 | Autosomal dominant, Autosomal recessive; Methionine adenosyltransferase deficiency, autosomal recessive, 250850 (3), Autosomal dominant, Autosomal recessive |
| MATN3 | 4148 | ENSG00000132031 | {Osteoarthritis susceptibility 2}, 140600 (3), Autosomal dominant; Spondyloepimetaphyseal dysplasia, Borochowitz-Cormier-Daire type, 608728 (3), Autosomal recessive; Epiphyseal dysplasia, multiple, 5, 607078 (3), Autosomal dominant |
| MBD4 | 8930 | ENSG00000129071 | Autosomal recessive |
| MC4R | 4160 | ENSG00000166603 | Obesity (BMIQ20), 618406 (3), Autosomal dominant, Autosomal recessive; {Obesity, resistance to (BMIQ20)}, 618406 (3), Autosomal dominant, Autosomal recessive |
| MDM2 | 4193 | ENSG00000135679 | Autosomal recessive |
| MEFV | 4210 | ENSG00000103313 | Neutrophilic dermatosis, acute febrile, 608068 (3), Autosomal dominant; Familial Mediterranean fever, AR, 249100 (3), Autosomal recessive; Familial Mediterranean fever, AD, 134610 (3), Autosomal dominant |
| MET | 4233 | ENSG00000105976 | Renal cell carcinoma, papillary, 1, familial and somatic, 605074 (3); ?Arthrogryposis, distal, type 11, 620019 (3), Autosomal dominant; Hepatocellular carcinoma, childhood type, somatic, 114550 (3); {Osteofibrous dysplasia, susceptibility to}, 607278 (3), Autosomal dominant; ?Deafness, autosomal recessive 97, 616705 (3), Autosomal recessive |
| METTL13 | 51603 | ENSG00000010165 | {?Deafness, autosomal recessive 26, modifier of}, 605429 (3), Autosomal dominant |
| MFN2 | 9927 | ENSG00000116688 | Lipomatosis, multiple symmetric, with or without peripheral neuropathy, 151800 (3), Autosomal recessive; Charcot-Marie-Tooth disease, axonal, type 2A2A, 609260 (3), Autosomal dominant; Charcot-Marie-Tooth disease, axonal, type 2A2B, 617087 (3), Autosomal recessive; Hereditary motor and sensory neuropathy VIA, 601152 (3), Autosomal dominant |
| MINPP1 | 9562 | ENSG00000107789 | 619527 (3), Autosomal recessive |
| MIR2861 | 100422910 | ENSG00000284547 | [Bone mineral density QTL 15], 613418 (3), Autosomal dominant, Autosomal recessive |
| MITF | 4286 | ENSG00000187098 | 614456 (3); Tietz albinism-deafness syndrome, 103500 (3), Autosomal dominant; COMMAD syndrome, 617306 (3), Autosomal recessive |
| MLH1 | 4292 | ENSG00000076242 | Lynch syndrome 2, 609310 (3); Muir-Torre syndrome, 158320 (3), Autosomal dominant; Mismatch repair cancer syndrome 1, 276300 (3), Autosomal recessive |
| MME | 4311 | ENSG00000196549 | ?Spinocerebellar ataxia 43, 617018 (3), Autosomal dominant; Charcot-Marie-Tooth disease, axonal, type 2T, 617017 (3), Autosomal dominant, Autosomal recessive |
| MMP13 | 4322 | ENSG00000137745 | ?Spondyloepimetaphyseal dysplasia, Missouri type, 602111 (3), Autosomal dominant; Metaphyseal anadysplasia 1, 602111 (3), Autosomal dominant; Metaphyseal dysplasia, Spahr type, 250400 (3), Autosomal recessive |
| MPL | 4352 | ENSG00000117400 | Myelofibrosis with myeloid metaplasia, somatic, 254450 (3); Thrombocythemia 2, 601977 (3), Somatic mutation, Autosomal dominant; Thrombocytopenia, congenital amegakaryocytic, 604498 (3), Autosomal recessive |
| MPO | 4353 | ENSG00000005381 | {Alzheimer disease, susceptibility to}, 104300 (3), Autosomal dominant; Myeloperoxidase deficiency, 254600 (3), Autosomal recessive; {Lung cancer, protection against, in smokers} (3) |
| MPZ | 4359 | ENSG00000158887 | dominant, Autosomal recessive; Charcot-Marie-Tooth disease, type 1B, 118200 (3), Autosomal dominant; Roussy-Levy syndrome, 180800 (3), Autosomal dominant; Charcot-Marie-Tooth disease, dominant intermediate D, 607791 (3), Autosomal dominant; Hypomyelinating neuropathy, congenital, 2, 618184 (3), Autosomal dominant; Charcot-Marie-Tooth disease, type 2J, 607736 (3), Autosomal dominant |
| MSH2 | 4436 | ENSG00000095002 | Lynch syndrome 1, 120435 (3), Autosomal dominant; Muir-Torre syndrome, 158320 (3), Autosomal dominant; Mismatch repair cancer syndrome 2, 619096 (3), Autosomal recessive |
| MSH6 | 2956 | ENSG00000116062 | Lynch syndrome 5, 614350 (3), Autosomal dominant; Mismatch repair cancer syndrome 3, 619097 (3), Autosomal recessive; {Endometrial cancer, familial}, 608089 (3), Somatic mutation, Autosomal dominant |
| MSTO1 | 55154 | ENSG00000125459 | Myopathy, mitochondrial, and ataxia, 617675 (3), Autosomal dominant, Autosomal recessive |

| | | | |
|---|---|---|---|
| MTHFR | 4524 | ENSG00000177000 | {Vascular disease, susceptibility to} (3); Homocystinuria due to MTHFR deficiency, 236250 (3), Autosomal recessive; {Thromboembolism, susceptibility to}, 188050 (3), Autosomal dominant; {Schizophrenia, susceptibility to}, 181500 (3), Autosomal dominant; {Neural tube defects, susceptibility to}, 601634 (3), Autosomal recessive |
| MVK | 4598 | ENSG00000110921 | Hyper-IgD syndrome, 260920 (3), Autosomal recessive; Porokeratosis 3, multiple types, 175900 (3), Autosomal dominant; Mevalonic aciduria, 610377 (3), Autosomal recessive |
| MYBPC1 | 4604 | ENSG00000196091 | Congenital myopathy 16, 618524 (3), Autosomal dominant; Lethal congenital contracture syndrome 4, 614915 (3), Autosomal recessive; Arthrogryposis, distal, type 1B, 614335 (3), Autosomal dominant |
| MYBPC3 | 4607 | ENSG00000134571 | Cardiomyopathy, hypertrophic, 4, 115197 (3), Autosomal dominant, Autosomal recessive; Cardiomyopathy, dilated, 1MM, 615396 (3), Autosomal dominant; Left ventricular noncompaction 10, 615396 (3), Autosomal dominant |
| MYH11 | 4629 | ENSG00000133392 | Megacystis-microcolon-intestinal hypoperistalsis syndrome 2, 619351 (3), Autosomal recessive; Aortic aneurysm, familial thoracic 4, 132900 (3), Autosomal dominant; Visceral myopathy 2, 619350 (3), Autosomal dominant |
| MYH2 | 4620 | ENSG00000125414 | Congenital myopathy 6 with ophthalmoplegia, 605637 (3), Autosomal dominant, Autosomal recessive |
| MYH3 | 4621 | ENSG00000109063 | pterygia, and spondylocarpotarsal fusion syndrome 1B, 618469 (3), Autosomal recessive; Arthrogryposis, distal, type 2B3 (Sheldon-Hall), 618436 (3), Autosomal dominant; Arthrogryposis, distal, type 2A (Freeman-Sheldon), 193700 (3), Autosomal dominant |
| MYH7 | 4625 | ENSG00000092054 | Laing distal myopathy, 160500 (3), Autosomal dominant; Cardiomyopathy, hypertrophic, 1, 192600 (3), Digenic dominant, Autosomal dominant; Left ventricular noncompaction 5, 613426 (3), Autosomal dominant; Cardiomyopathy, dilated, 1S, 613426 (3), Autosomal dominant; Congenital myopathy 7B, myosin storage, autosomal recessive, 255160 (3), Autosomal recessive; Congenital myopathy 7A, myosin storage, autosomal dominant, 608358 (3), Autosomal dominant |
| MYL11 | 29895 | ENSG00000180209 | Arthrogryposis, distal, type 1C, 619110 (3), Autosomal dominant, Autosomal recessive |
| MYL2 | 4633 | ENSG00000111245 | Cardiomyopathy, hypertrophic, 10, 608758 (3), Autosomal dominant; Myopathy, myofibrillar, 12, infantile-onset, with cardiomyopathy, 619424 (3), Autosomal recessive |
| MYL3 | 4634 | ENSG00000160808 | Cardiomyopathy, hypertrophic, 8, 608751 (3), Autosomal dominant, Autosomal recessive |
| MYLK | 4638 | ENSG00000065534 | Megacystis-microcolon-intestinal hypoperistalsis syndrome 1, 249210 (3), Autosomal recessive; Aortic aneurysm, familial thoracic 7, 613780 (3), Autosomal dominant |
| MYO6 | 4646 | ENSG00000196586 | Deafness, autosomal dominant 22, with hypertrophic cardiomyopathy, 606346 (3), Autosomal dominant; Deafness, autosomal dominant 22, 606346 (3), Autosomal dominant; Deafness, autosomal recessive 37, 607821 (3), Autosomal recessive |
| MYO7A | 4647 | ENSG00000137474 | Deafness, autosomal recessive 2, 600060 (3), Autosomal recessive; Usher syndrome, type 1B, 276900 (3), Autosomal recessive; Deafness, autosomal dominant 11, 601317 (3), Autosomal dominant |
| MYPN | 84665 | ENSG00000138347 | recessive; Cardiomyopathy, familial restrictive, 4, 615248 (3), Autosomal dominant; Cardiomyopathy, dilated, 1KK, 615248 (3), Autosomal dominant |
| NAGLU | 4669 | ENSG00000108784 | ?Charcot-Marie-Tooth disease, axonal, type 2V, 616491 (3), Autosomal dominant; Mucopolysaccharidosis type IIIB (Sanfilippo B), 252920 (3), Autosomal recessive |
| NALCN | 259232 | ENSG00000102452 | Congenital contractures of the limbs and face, hypotonia, and developmental delay, 616266 (3), Autosomal dominant; Hypotonia, infantile, with psychomotor retardation and characteristic facies 1, 615419 (3), Autosomal recessive |
| NARS1 | 4677 | ENSG00000134440 | 619092 (3), Autosomal dominant; Neurodevelopmental disorder with microcephaly, impaired language, and gait abnormalities, autosomal recessive, 619091 (3), Autosomal recessive |
| NEFH | 4744 | ENSG00000100285 | Charcot-Marie-Tooth disease, axonal, type 2CC, 616924 (3), Autosomal dominant; {?Amyotrophic lateral sclerosis, susceptibility to}, 105400 (3), Autosomal dominant, Autosomal recessive |
| NEFL | 4747 | ENSG00000277586 | dominant intermediate G, 617882 (3), Autosomal dominant; Charcot-Marie-Tooth disease, type 2E, 607684 (3), Autosomal dominant |

| | | | |
|---|---|---|---|
| NEK1 | 4750 | ENSG00000137601 | Short-rib thoracic dysplasia 6 with or without polydactyly, 263520 (3), Digenic recessive, Autosomal recessive; {Amyotrophic lateral sclerosis, susceptibility to, 24}, 617892 (3), Autosomal dominant |
| NLRP1 | 22861 | ENSG00000091592 | {Vitiligo-associated multiple autoimmune disease susceptibility 1}, 606579 (3); ?Respiratory papillomatosis, juvenile recurrent, congenital, 618803 (3), Autosomal recessive; Autoinflammation with arthritis and dyskeratosis, 617388 (3), Autosomal dominant, Autosomal recessive; Palmoplantar carcinoma, multiple self-healing, 615225 (3), Autosomal dominant |
| NOP10 | 55505 | ENSG00000182117 | hearing impairment, nephrotic syndrome, and enterocolitis 2, 620425 (3), Autosomal recessive; ?Dyskeratosis congenita, autosomal recessive 1, 224230 (3), Autosomal recessive |
| NPPA | 4878 | ENSG00000175206 | Atrial standstill 2, 615745 (3), Autosomal recessive; Atrial fibrillation, familial, 6, 612201 (3), Autosomal dominant |
| NPR2 | 4882 | ENSG00000159899 | abnormalities, 616255 (3), Autosomal dominant; Acromesomelic dysplasia 1, Maroteaux type, 602875 (3), Autosomal recessive |
| NR0B2 | 8431 | ENSG00000131910 | Obesity, mild, early-onset, 601665 (3), Multifactorial, Autosomal dominant, Autosomal recessive |
| NR2E3 | 10002 | ENSG00000278570 | Autosomal recessive |
| NRL | 4901 | ENSG00000129535 | (3) |
| NUS1 | 116150 | ENSG00000153989 | Intellectual developmental disorder, autosomal dominant 55, with seizures, 617831 (3), Autosomal dominant; ?Congenital disorder of glycosylation, type 1aa, 617082 (3), Autosomal recessive |
| OPA1 | 4976 | ENSG00000198836 | Optic atrophy plus syndrome, 125250 (3), Autosomal dominant; {Glaucoma, normal tension, susceptibility to}, 606657 (3); Optic atrophy 1, 165500 (3), Autosomal dominant; Behr syndrome, 210000 (3), Autosomal recessive; ?Mitochondrial DNA depletion syndrome 14 (encephalocardiomyopathic type), 616896 (3), Autosomal recessive |
| OPA3 | 80207 | ENSG00000125741 | dominant |
| OPLAH | 26873 | ENSG00000178814 | 5-oxoprolinase deficiency, 260005 (3), Autosomal dominant, Autosomal recessive |
| OPTN | 10133 | ENSG00000123240 | Glaucoma 1, open angle, E, 137760 (3), Autosomal dominant; Amyotrophic lateral sclerosis 12 with or without frontotemporal dementia, 613435 (3), Autosomal dominant, Autosomal recessive; {Glaucoma, normal tension, susceptibility to}, 606657 (3) |
| ORAI1 | 84876 | ENSG00000276045 | Immunodeficiency 9, 612782 (3), Autosomal recessive; Myopathy, tubular aggregate, 2, 615883 (3), Autosomal dominant |
| OTULIN | 90268 | ENSG00000154124 | Autoinflammation, panniculitis, and dermatosis syndrome, 617099 (3), Autosomal recessive; {Immunodeficiency 107, susceptibility to invasive staphylococcus aureus infection}, 619986 (3), Autosomal dominant |
| PARN | 5073 | ENSG00000140694 | Dyskeratosis congenita, autosomal recessive 6, 616353 (3), Autosomal recessive; Pulmonary fibrosis and/or bone marrow failure syndrome, telomere-related, 4, 616371 (3), Autosomal dominant |
| PAX3 | 5077 | ENSG00000135903 | Autosomal dominant, Autosomal recessive; Waardenburg syndrome, type 1, 193500 (3), Autosomal dominant; Rhabdomyosarcoma 2, alveolar, 268220 (3), Somatic mutation |
| PAX4 | 5078 | ENSG00000106331 | {Diabetes mellitus, ketosis-prone, susceptibility to}, 612227 (3), Autosomal dominant, Autosomal recessive; Maturity-onset diabetes of the young, type IX, 612225 (3); Diabetes mellitus, type 2, 125853 (3), Autosomal dominant |
| PDE10A | 10846 | ENSG00000112541 | Striatal degeneration, autosomal dominant, 616922 (3), Autosomal dominant; Dyskinesia, limb and orofacial, infantile-onset, 616921 (3), Autosomal recessive |
| PDE6B | 5158 | ENSG00000133256 | 163500 (3), Autosomal dominant |
| PDE6H | 5149 | ENSG00000139053 | dominant, Autosomal recessive |
| PDX1 | 3651 | ENSG00000139515 | {Diabetes mellitus, type II, susceptibility to}, 125853 (3), Autosomal dominant; Pancreatic agenesis 1, 260370 (3), Autosomal recessive; MODY, type IV, 606392 (3) |
| PERP | 64065 | ENSG00000112378 | Autosomal dominant |
| PEX6 | 5190 | ENSG00000124587 | Peroxisome biogenesis disorder 4B, 614863 (3), Autosomal dominant, Autosomal recessive; Peroxisome biogenesis disorder 4A (Zellweger), 614862 (3), Autosomal recessive; Heimler syndrome 2, 616617 (3), Autosomal recessive |

| | | | |
|---|---|---|---|
| PIEZO1 | 9780 | ENSG00000103335 | Dehydrated hereditary stomatocytosis with or without pseudohyperkalemia and/or perinatal edema, 194380 (3), Autosomal dominant |
| PIEZO2 | 63895 | ENSG00000154864 | 617146 (3), Autosomal recessive; Arthrogryposis, distal, type 3, 114300 (3), Autosomal dominant; ?Marden-Walker syndrome, 248700 (3), Autosomal dominant |
| PIGT | 51604 | ENSG00000124155 | ?Paroxysmal nocturnal hemoglobinuria 2, 615399 (3), Somatic mutation, Autosomal dominant; Multiple congenital anomalies-hypotonia-seizures syndrome 3, 615398 (3), Autosomal recessive |
| PIK3CD | 5293 | ENSG00000171608 | Immunodeficiency 14A, autosomal dominant, 615513 (3), Autosomal dominant; Immunodeficiency 14B, autosomal recessive, 619281 (3), Autosomal recessive; ?Roifman-Chitayat syndrome, digenic, 613328 (3), Digenic recessive |
| PIK3R1 | 5295 | ENSG00000145675 | Immunodeficiency 36, 616005 (3), Autosomal dominant; ?Agammaglobulinemia 7, autosomal recessive, 615214 (3), Autosomal recessive; SHORT syndrome, 269880 (3), Autosomal dominant |
| PITX3 | 5309 | ENSG00000107859 | subtypes, 107250 (3), Autosomal dominant; Cataract 11, syndromic, autosomal recessive, 610623 (3), Autosomal dominant, Autosomal recessive |
| PKLR | 5313 | ENSG00000143627 | Autosomal recessive |
| PLCB4 | 5332 | ENSG00000101333 | dominant |
| PLCD1 | 5333 | ENSG00000187091 | Nail disorder, nonsyndromic congenital, 3, (leukonychia), 151600 (3), Autosomal dominant, Autosomal recessive |
| PLEC | 5339 | ENSG00000178209 | ?Epidermolysis bullosa simplex 5D, generalized intermediate, autosomal recessive, 616487 (3), Autosomal recessive; Epidermolysis bullosa simplex 5B, with muscular dystrophy, 226670 (3), Autosomal recessive; Epidermolysis bullosa simplex 5C, with pyloric atresia, 612138 (3), Autosomal recessive; Epidermolysis bullosa simplex 5A, Ogna type, 131950 (3), Autosomal dominant; Muscular dystrophy, limb-girdle, autosomal recessive 17, 613723 (3), Autosomal recessive |
| PLEKHM1 | 9842 | ENSG00000225190 | Autosomal dominant |
| PLG | 5340 | ENSG00000122194 | Dysplasminogenemia, 217090 (3), Autosomal recessive; Angioedema, hereditary, 4, 619360 (3), Autosomal dominant; Plasminogen deficiency, type I, 217090 (3), Autosomal recessive |
| PMP22 | 5376 | ENSG00000109099 | dominant; Charcot-Marie-Tooth disease, type 1E, 118300 (3), Autosomal dominant; ?Neuropathy, inflammatory demyelinating, 139393 (3), ?Autosomal dominant; Neuropathy, recurrent, with pressure palsies, 162500 (3), Autosomal dominant; Dejerine-Sottas disease, 145900 (3), Autosomal dominant, Autosomal recessive |
| PNPT1 | 87178 | ENSG00000138035 | neurodegeneration, 614934 (3), Autosomal recessive; Combined oxidative phosphorylation deficiency 13, 614932 (3), Autosomal recessive |
| POGLUT1 | 56983 | ENSG00000163389 | (3), Autosomal recessive |
| POLE | 5426 | ENSG00000177084 | {Colorectal cancer, susceptibility to, 12}, 615083 (3), Autosomal dominant; FILS syndrome, 615139 (3), Autosomal recessive; IMAGE-I syndrome, 618336 (3), Autosomal recessive |
| POLG | 5428 | ENSG00000140521 | Mitochondrial recessive ataxia syndrome (includes SANDO and SCAE), 607459 (3), Autosomal recessive; Mitochondrial DNA depletion syndrome 4B (MNGIE type), 613662 (3), Autosomal recessive; Mitochondrial DNA depletion syndrome 4A (Alpers type), 203700 (3), Autosomal recessive; Progressive external ophthalmoplegia, autosomal dominant 1, 157640 (3), Autosomal dominant; Progressive external ophthalmoplegia, autosomal recessive 1, 258450 (3), Autosomal recessive |
| POLG2 | 11232 | ENSG00000256525 | Progressive external ophthalmoplegia with mitochondrial DNA deletions, autosomal dominant 4, 610131 (3), Autosomal dominant; ?Mitochondrial DNA depletion syndrome 16 (hepatic type), 618528 (3), Autosomal recessive; ?Mitochondrial DNA depletion syndrome 16B (neuroophthalmic type), 619425 (3), Autosomal recessive |
| POLR1D | 51082 | ENSG00000186184 | Treacher Collins syndrome 2, 613717 (3), Autosomal dominant, Autosomal recessive |
| POLR3B | 55703 | ENSG00000013503 | Leukodystrophy, hypomyelinating, 8, with or without oligodontia and/or hypogonadotropic hypogonadism, 614381 (3), Autosomal recessive; Charcot-Marie-Tooth disease, demyelinating, type 1I, 619742 (3), Autosomal dominant |
| POLRMT | 5442 | ENSG00000099821 | Combined oxidative phosphorylation deficiency 55, 619743 (3), Autosomal dominant, Autosomal recessive |

| | | | |
|---|---|---|---|
| POMC | 5443 | ENSG00000115138 | {Obesity, early-onset, susceptibility to}, 601665 (3), Multifactorial, Autosomal dominant, Autosomal recessive; Obesity, adrenal insufficiency, and red hair due to POMC deficiency, 609734 (3), Autosomal recessive |
| POMP | 51371 | ENSG00000132963 | Proteasome-associated autoinflammatory syndrome 2, 618048 (3), Autosomal dominant; Keratosis linearis with ichthyosis congenita and sclerosing keratoderma, 601952 (3), Autosomal recessive |
| POT1 | 25913 | ENSG00000128513 | cysts 3, 620368 (3), Autosomal recessive; ?Pulmonary fibrosis and/or bone marrow failure syndrome, telomere-related, 8, 620367 (3), Autosomal dominant |
| POU1F1 | 5449 | ENSG00000064835 | Pituitary hormone deficiency, combined or isolated, 1, 613038 (3), Autosomal dominant, Autosomal recessive |
| PPARG | 5468 | ENSG00000132170 | {Diabetes, type 2}, 125853 (3), Autosomal dominant; Insulin resistance, severe, digenic, 604367 (3), Autosomal dominant; Lipodystrophy, familial partial, type 3, 604367 (3), Autosomal dominant; [Obesity, resistance to] (3); Obesity, severe, 601665 (3), Multifactorial, Autosomal dominant, Autosomal recessive; Carotid intimal medial thickness 1, 609338 (3) |
| PPOX | 5498 | ENSG00000143224 | Variegate porphyria, childhood-onset, 620483 (3), Autosomal recessive; Variegate porphyria, 176200 (3), Autosomal dominant |
| PPP1R17 | 10842 | ENSG00000106341 | {Hypercholesterolemia, susceptibility to}, 143890 (3), Autosomal dominant, Autosomal recessive |
| PPP2R3C | 55012 | ENSG00000092020 | Autosomal recessive |
| PRDX3 | 10935 | ENSG00000165672 | Spinocerebellar ataxia, autosomal recessive 32, 619862 (3), Autosomal recessive; Corneal dystrophy, punctiform and polychromatic pre-Descemet, 619871 (3), Autosomal dominant |
| PRLR | 5618 | ENSG00000113494 | Autosomal recessive |
| PROC | 5624 | ENSG00000115718 | Thrombophilia 3 due to protein C deficiency, autosomal dominant, 176860 (3), Autosomal dominant; Thrombophilia 3 due to protein C deficiency, autosomal recessive, 612304 (3), Autosomal recessive |
| PRODH | 5625 | ENSG00000100033 | recessive |
| PROM1 | 8842 | ENSG00000007062 | Stargardt disease 4, 603786 (3), Autosomal dominant; Cone-rod dystrophy 12, 612657 (3), Autosomal dominant, Autosomal recessive |
| PROS1 | 5627 | ENSG00000184500 | Thrombophilia 5 due to protein S deficiency, autosomal recessive, 614514 (3), Autosomal recessive; Thrombophilia 5 due to protein S deficiency, autosomal dominant, 612336 (3), Autosomal dominant |
| PRPH | 5630 | ENSG00000135406 | {Amyotrophic lateral sclerosis, susceptibility to}, 105400 (3), Autosomal dominant, Autosomal recessive |
| PRPH2 | 5961 | ENSG00000112619 | Autosomal dominant; Retinitis punctata albescens, 136880 (3), Autosomal dominant, Autosomal recessive; Leber congenital amaurosis 18, 608133 (3), Digenic dominant, Autosomal dominant, Autosomal recessive; Macular dystrophy, vitelliform, 3, 608161 (3), Autosomal dominant; Retinitis pigmentosa 7 and digenic form, 608133 (3), Digenic dominant, Autosomal dominant, Autosomal recessive |
| PRRX1 | 5396 | ENSG00000116132 | Agnathia-otocephaly complex, 202650 (3), Autosomal dominant, Autosomal recessive |
| PRX | 57716 | ENSG00000105227 | dominant, Autosomal recessive |
| PSAP | 5660 | ENSG00000197746 | Combined SAP deficiency, 611721 (3), Autosomal recessive; Krabbe disease, atypical, 611722 (3), Autosomal recessive; Metachromatic leukodystrophy due to SAP-b deficiency, 249900 (3), Autosomal recessive; Gaucher disease, atypical, 610539 (3); {Parkinson disease 24, autosomal dominant, susceptibility to}, 619491 (3), Autosomal dominant |
| PTH | 5741 | ENSG00000152266 | Hypoparathyroidism, familial isolated 1, 146200 (3), Autosomal dominant, Autosomal recessive |
| PTH1R | 5745 | ENSG00000160801 | recessive; Failure of tooth eruption, primary, 125350 (3), Autosomal dominant; Chondrodysplasia, Blomstrand type, 215045 (3), Autosomal recessive |
| PTPN22 | 26191 | ENSG00000134242 | {Rheumatoid arthritis, susceptibility to}, 180300 (3); {Systemic lupus erythematosus susceptibility to}, 152700 (3), Autosomal dominant; {Diabetes, type 1, susceptibility to}, 222100 (3), Autosomal recessive |
| PTPRQ | 374462 | ENSG00000139304 | Autosomal recessive |

| | | | |
|---|---|---|---|
| RAC2 | 5880 | ENSG00000128340 | Immunodeficiency 73A with defective neutrophil chemotaxix and leukocytosis, 608203 (3), Autosomal dominant; ?Immunodeficiency 73C with defective neutrophil chemotaxis and hypogammaglobulinemia, 618987 (3), Autosomal recessive; Immunodeficiency 73B with defective neutrophil chemotaxis and lymphopenia, 618986 (3), Autosomal dominant |
| RAD21 | 5885 | ENSG00000164754 | Cornelia de Lange syndrome 4, 614701 (3), Autosomal dominant; ?Mungan syndrome, 611376 (3), Autosomal recessive |
| RARB | 5915 | ENSG00000077092 | Microphthalmia, syndromic 12, 615524 (3), Autosomal dominant, Autosomal recessive |
| RAX2 | 84839 | ENSG00000173976 | Retinitis pigmentosa 95, 620102 (3), Autosomal recessive; Cone-rod dystrophy 11, 610381 (3), Autosomal dominant; ?Macular degeneration, age-related, 6, 613757 (3) |
| RBP4 | 5950 | ENSG00000138207 | Microphthalmia, isolated, with coloboma 10, 616428 (3), Autosomal dominant; Retinal dystrophy, iris coloboma, and comedogenic acne syndrome, 615147 (3), Autosomal recessive |
| RDH12 | 145226 | ENSG00000139988 | Leber congenital amaurosis 13, 612712 (3), Autosomal dominant, Autosomal recessive |
| RDH5 | 5959 | ENSG00000135437 | Fundus albipunctatus, 136880 (3), Autosomal dominant, Autosomal recessive |
| REEP1 | 65055 | ENSG00000068615 | autosomal dominant, 610250 (3), Autosomal dominant; ?Neuronopathy, distal hereditary motor, autosomal dominant 12, 614751 (3), Autosomal dominant |
| REEP2 | 51308 | ENSG00000132563 | Spastic paraplegia 72A, autosomal dominant, 615625 (3), Autosomal dominant; ?Spastic paraplegia 72B, autosomal recessive, 620606 (3), Autosomal recessive |
| RELN | 5649 | ENSG00000189056 | [illegible] Autosomal recessive |
| REN | 5972 | ENSG00000143839 | Renal tubular dysgenesis, 267430 (3), Autosomal recessive; [Hyperproreninemia] (3); Tubulointerstitial kidney disease, autosomal dominant, 4, 613092 (3), Autosomal dominant |
| RHAG | 6005 | ENSG00000112077 | [illegible] (3), Autosomal recessive |
| RHO | 6010 | ENSG00000163914 | autosomal dominant or recessive, 613731 (3), Autosomal dominant, Autosomal recessive; Retinitis punctata albescens, 136880 (3), Autosomal dominant, Autosomal recessive |
| RIPK1 | 8737 | ENSG00000137275 | Immunodeficiency 57 with autoinflammation, 618108 (3), Autosomal recessive; Autoinflammation with episodic fever and lymphadenopathy, 618852 (3), Autosomal dominant |
| RIPOR2 | 9750 | ENSG00000111913 | [illegible] Autosomal recessive |
| RLBP1 | 6017 | ENSG00000140522 | albescens, 136880 (3), Autosomal dominant, Autosomal recessive; Fundus albipunctatus, 136880 (3), Autosomal dominant, Autosomal recessive |
| RNF170 | 81790 | ENSG00000120925 | [illegible] 619686 (3), Autosomal recessive |
| RNF213 | 57674 | ENSG00000173821 | {Moyamoya disease 2, susceptibility to}, 607151 (3), Autosomal dominant, Autosomal recessive |
| ROBO1 | 6091 | ENSG00000169855 | Pituitary hormone deficiency, combined or isolated, 8, 620303 (3), Autosomal dominant; Neurooculorenal syndrome, 620305 (3), Autosomal recessive; ?Nystagmus 8, congenital, autosomal recessive, 257400 (3), Autosomal recessive |
| ROM1 | 6094 | ENSG00000149489 | Retinitis pigmentosa 7, digenic form, 608133 (3), Digenic dominant, Autosomal dominant, Autosomal recessive |
| ROR2 | 4920 | ENSG00000169071 | [illegible] recessive |
| RP1 | 6101 | ENSG00000104237 | Retinitis pigmentosa 1, 180100 (3), Autosomal dominant, Autosomal recessive |
| RP1L1 | 94137 | ENSG00000183638 | Occult macular dystrophy, 613587 (3), Autosomal dominant; Retinitis pigmentosa 88, 618826 (3), Autosomal recessive |
| RPE65 | 6121 | ENSG00000116745 | Retinitis pigmentosa 20, 613794 (3), Autosomal recessive; Retinitis pigmentosa 87 with choroidal involvement, 618697 (3), Autosomal dominant; Leber congenital amaurosis 2, 204100 (3), Autosomal recessive |
| RRM1 | 6240 | ENSG00000167325 | [illegible] dominant, Autosomal recessive |

| | | | |
|---|---|---|---|
| RRM2B | 50484 | ENSG00000048392 | syndrome 8A (encephalomyopathic type with renal tubulopathy), 612075 (3), Autosomal recessive; Rod-cone dystrophy, sensorineural deafness, and Fanconi-type renal dysfunction, 268315 (3), Autosomal recessive; Progressive external ophthalmoplegia with mitochondrial DNA deletions, autosomal dominant 5, 613077 (3), Autosomal dominant |
| RTEL1 | 51750 | ENSG00000258366 | Dyskeratosis congenita, autosomal dominant 4, 615190 (3), Autosomal dominant, Autosomal recessive; Dyskeratosis congenita, autosomal recessive 5, 615190 (3), Autosomal dominant, Autosomal recessive; Pulmonary fibrosis and/or bone marrow failure syndrome, telomere-related, 3, 616373 (3), Autosomal dominant |
| RYR1 | 6261 | ENSG00000196218 | Congenital myopathy 1B, autosomal recessive, 255320 (3), Autosomal recessive; Congenital myopathy 1A, autosomal dominant, with susceptibility to malignant hyperthermia, 117000 (3), Autosomal dominant; King-Denborough syndrome, 619542 (3), Autosomal dominant; {Malignant hyperthermia susceptibility 1}, 145600 (3), Autosomal dominant |
| SAG | 6295 | ENSG00000130561 | Retinitis pigmentosa 47, autosomal recessive, 613758 (3), Autosomal recessive; Retinitis pigmentosa 96, autosomal dominant, 620228 (3), Autosomal dominant; Oguchi disease-1, 258100 (3), Autosomal recessive |
| SAMD9 | 54809 | ENSG00000205413 | Tumoral calcinosis, familial, normophosphatemic, 610455 (3), Autosomal recessive; Monosomy 7 myelodysplasia and leukemia syndrome 2, 619041 (3), Autosomal dominant; MIRAGE syndrome, 617053 (3), Autosomal dominant |
| SAMHD1 | 25939 | ENSG00000101347 | ?Chilblain lupus 2, 614415 (3), Autosomal dominant; Aicardi-Goutieres syndrome 5, 612952 (3), Autosomal recessive |
| SASH1 | 23328 | ENSG00000111961 | Dyschromatosis universalis hereditaria 1, 127500 (3), Autosomal dominant; ?Cancer, alopecia, pigment dyscrasia, onychodystrophy, and keratoderma, 618373 (3), Autosomal recessive |
| SCN1B | 6324 | ENSG00000105711 | Generalized epilepsy with febrile seizures plus, type 1, 604233 (3), Autosomal dominant; Developmental and epileptic encephalopathy 52, 617350 (3), Autosomal recessive; Cardiac conduction defect, nonspecific, 612838 (3); Atrial fibrillation, familial, 13, 615377 (3), Autosomal dominant; Brugada syndrome 5, 612838 (3) |
| SCN4A | 6329 | ENSG00000007314 | Congenital myopathy 22B, severe fetal, 620369 (3), Autosomal recessive; Hypokalemic periodic paralysis, type 2, 613345 (3), Autosomal dominant; Myotonia congenita, atypical, acetazolamide-responsive, 608390 (3), Autosomal dominant; Myasthenic syndrome, congenital, 16, 614198 (3), Autosomal recessive; Congenital myopathy 22A, classic, 620351 (3), Autosomal recessive |
| SCN5A | 6331 | ENSG00000183873 | Cardiomyopathy, dilated, 1E, 601154 (3), Autosomal dominant; Heart block, nonprogressive, 113900 (3), Autosomal dominant; Long QT syndrome 3, 603830 (3), Autosomal dominant; Sick sinus syndrome 1, 608567 (3), Autosomal recessive; Brugada syndrome 1, 601144 (3), Autosomal dominant; Atrial fibrillation, familial, 10, 614022 (3), Autosomal dominant; {Sudden infant death syndrome, susceptibility to}, 272120 (3), Autosomal recessive |
| SCN9A | 6335 | ENSG00000169432 | Erythermalgia, primary, 133020 (3), Autosomal dominant; Insensitivity to pain, congenital, 243000 (3), Autosomal recessive; Small fiber neuropathy, 133020 (3), Autosomal dominant; Paroxysmal extreme pain disorder, 167400 (3), Autosomal dominant; Neuropathy, hereditary sensory and autonomic, type IID, 243000 (3), Autosomal recessive |
| SCNN1A | 6337 | ENSG00000111319 | Pseudohypoaldosteronism, type IB1, autosomal recessive, 264350 (3), Autosomal recessive; ?Liddle syndrome 3, 618126 (3), Autosomal dominant; Bronchiectasis with or without elevated sweat chloride 2, 613021 (3), Autosomal dominant |
| SCNN1B | 6338 | ENSG00000168447 | Bronchiectasis with or without elevated sweat chloride 1, 211400 (3), Autosomal dominant; Pseudohypoaldosteronism, type IB2, autosomal recessive, 620125 (3), Autosomal recessive; Liddle syndrome 1, 177200 (3), Autosomal dominant |
| SCNN1G | 6340 | ENSG00000166828 | Bronchiectasis with or without elevated sweat chloride 3, 613071 (3), Autosomal dominant; Pseudohypoaldosteronism, type IB3, autosomal recessive, 620126 (3), Autosomal recessive; Liddle syndrome 2, 618114 (3), Autosomal dominant |
| SCO2 | 9997 | ENSG00000284194 | [illegible] recessive |
| SDC3 | 9672 | ENSG00000162512 | {Obesity, association with}, 601665 (3), Multifactorial, Autosomal dominant, Autosomal recessive |
| SDHA | 6389 | ENSG00000073578 | Cardiomyopathy, dilated, 1GG, 613642 (3), Autosomal recessive; Mitochondrial complex II deficiency, nuclear type 1, 252011 (3), Autosomal recessive; Neurodegeneration with ataxia and late-onset optic atrophy, 619259 (3), Autosomal dominant; Pheochromocytoma/paraganglioma syndrome 5, 614165 (3), Autosomal dominant |

| | | | |
|---|---|---|---|
| SDHB | 6390 | ENSG00000117118 | Pheochromocytoma/paraganglioma syndrome 4, 115310 (3), Autosomal dominant; Mitochondrial complex II deficiency, nuclear type 4, 619224 (3), Autosomal recessive; Gastrointestinal stromal tumor, 606764 (3), Autosomal dominant, Isolated cases; Paraganglioma and gastric stromal sarcoma, 606864 (3) |
| SDHD | 6392 | ENSG00000204370 | Pheochromocytoma/paraganglioma syndrome 1, 168000 (3), Autosomal dominant; Paraganglioma and gastric stromal sarcoma, 606864 (3); Mitochondrial complex II deficiency, nuclear type 3, 619167 (3), Autosomal recessive |
| SEC23A | 10484 | ENSG00000100934 | Craniolenticulosutural dysplasia, 607812 (3), Autosomal dominant, Autosomal recessive |
| SEC23B | 10483 | ENSG00000101310 | [illegible] recessive |
| SERPINA6 | 866 | ENSG00000170099 | Corticosteroid-binding globulin deficiency, 611489 (3), Autosomal dominant, Autosomal recessive |
| SERPINC1 | 462 | ENSG00000117601 | Thrombophilia 7 due to antithrombin III deficiency, 613118 (3), Autosomal dominant, Autosomal recessive |
| SERPINE1 | 5054 | ENSG00000106366 | Plasminogen activator inhibitor-1 deficiency, 613329 (3), Autosomal dominant, Autosomal recessive; {Transcription of plasminogen activator inhibitor, modulator of} (3) |
| SERPING1 | 710 | ENSG00000149131 | Angioedema, hereditary, 1 and 2, 106100 (3), Autosomal dominant, Autosomal recessive; Complement component 4, partial deficiency of, 120790 (3), Autosomal dominant |
| SETX | 23064 | ENSG00000107290 | Spinocerebellar ataxia, autosomal recessive, with axonal neuropathy 2, 606002 (3), Autosomal recessive; Amyotrophic lateral sclerosis 4, juvenile, 602433 (3), Autosomal dominant |
| SFTPA1 | 653509 | ENSG00000122852 | Interstitial lung disease 1, 619611 (3), Autosomal dominant, Autosomal recessive |
| SH3TC2 | 79628 | ENSG00000169247 | [illegible] (3), Autosomal dominant |
| SLC12A2 | 6558 | ENSG00000064651 | Kilquist syndrome, 619080 (3), Autosomal recessive; Delpire-McNeill syndrome, 619083 (3), Autosomal dominant; Deafness, autosomal dominant 78, 619081 (3), Autosomal dominant |
| SLC12A5 | 57468 | ENSG00000124140 | {Epilepsy, idiopathic generalized, susceptibility to, 14}, 616685 (3), Autosomal dominant; Developmental and epileptic encephalopathy 34, 616645 (3), Autosomal recessive |
| SLC12A6 | 9990 | ENSG00000140199 | Agenesis of the corpus callosum with peripheral neuropathy, 218000 (3), Autosomal recessive; Charcot-Marie-Tooth disease, axonal, type 2II, 620068 (3), Autosomal dominant |
| SLC16A1 | 6566 | ENSG00000155380 | Hyperinsulinemic hypoglycemia, familial, 7, 610021 (3), Autosomal dominant; Erythrocyte lactate transporter defect, 245340 (3), Autosomal dominant; Monocarboxylate transporter 1 deficiency, 616095 (3), Autosomal dominant, Autosomal recessive |
| SLC25A4 | 291 | ENSG00000151729 | Mitochondrial DNA depletion syndrome 12B (cardiomyopathic type) AR, 615418 (3), Autosomal recessive; Progressive external ophthalmoplegia with mitochondrial DNA deletions, autosomal dominant 2, 609283 (3), Autosomal dominant; Mitochondrial DNA depletion syndrome 12A (cardiomyopathic type) AD, 617184 (3), Autosomal dominant |
| SLC26A8 | 116369 | ENSG00000112053 | Spermatogenic failure 3, 606766 (3), Autosomal dominant, Autosomal recessive |
| SLC2A1 | 6513 | ENSG00000117394 | [illegible] dominant, Autosomal recessive; Stomatin-deficient cryohydrocytosis with neurologic defects, 608885 (3), Autosomal dominant; {Epilepsy, idiopathic generalized, susceptibility to, 12}, 614847 (3), Autosomal dominant; GLUT1 deficiency syndrome 2, childhood onset, 612126 (3), Autosomal dominant |
| SLC2A2 | 6514 | ENSG00000163581 | [illegible] dominant |
| SLC2A9 | 56606 | ENSG00000109667 | {Uric acid concentration, serum, QTL 2}, 612076 (3), Autosomal dominant, Autosomal recessive; Hypouricemia, renal, 2, 612076 (3), Autosomal dominant, Autosomal recessive |
| SLC33A1 | 9197 | ENSG00000169359 | [illegible] Autosomal recessive |
| SLC34A1 | 6569 | ENSG00000131183 | ?Fanconi renotubular syndrome 2, 613388 (3), Autosomal recessive; Hypercalcemia, infantile, 2, 616963 (3), Autosomal recessive; Nephrolithiasis/osteoporosis, hypophosphatemic, 1, 612286 (3), Autosomal dominant |
| SLC36A2 | 153201 | ENSG00000186335 | [Iminoglycinuria], 242600 (3), Digenic recessive, Autosomal recessive; [Hyperglycinuria], 138500 (3), Autosomal dominant |

| | | | |
|---|---|---|---|
| SLC37A4 | 2542 | ENSG00000137700 | Glycogen storage disease Ib, 232220 (3), Autosomal recessive; Congenital disorder of glycosylation, type IIw, 619525 (3), Autosomal dominant; Glycogen storage disease Ic, 232240 (3), Autosomal recessive |
| SLC39A14 | 23516 | ENSG00000104635 | [illegible] recessive |
| SLC3A1 | 6519 | ENSG00000138079 | Cystinuria, 220100 (3), Autosomal dominant, Autosomal recessive |
| SLC4A1 | 6521 | ENSG00000004939 | dominant; [Blood group, Waldner], 112010 (3); Spherocytosis, type 4, 612653 (3), Autosomal dominant; [Blood group, Froese], 601551 (3); Distal renal tubular acidosis 4 with hemolytic anemia, 611590 (3), Autosomal recessive; {Malaria, resistance to}, 611162 (3); Cryohydrocytosis, 185020 (3), Autosomal dominant; Ovalocytosis, SA type, 166900 (3), Autosomal dominant; [Blood group, Diego], 110500 (3) |
| SLC5A2 | 6524 | ENSG00000140675 | Renal glucosuria, 233100 (3), Autosomal dominant, Autosomal recessive |
| SLC5A7 | 60482 | ENSG00000115665 | Neuronopathy, distal hereditary motor, autosomal dominant 7, 158580 (3), Autosomal dominant; Myasthenic syndrome, congenital, 20, presynaptic, 617143 (3), Autosomal recessive |
| SLC6A5 | 9152 | ENSG00000165970 | Hyperekplexia 3, 614618 (3), Autosomal dominant, Autosomal recessive |
| SLC7A9 | 11136 | ENSG00000021488 | Cystinuria, 220100 (3), Autosomal dominant, Autosomal recessive |
| SLCO2A1 | 6578 | ENSG00000174640 | Hypertrophic osteoarthropathy, primary, autosomal dominant, 167100 (3), Autosomal dominant; Hypertrophic osteoarthropathy, primary, autosomal recessive 2, 614441 (3), Autosomal recessive |
| SOD1 | 6647 | ENSG00000142168 | Spastic tetraplegia and axial hypotonia, progressive, 618598 (3), Autosomal recessive; Amyotrophic lateral sclerosis 1, 105400 (3), Autosomal dominant, Autosomal recessive |
| SOHLH1 | 402381 | ENSG00000165643 | Ovarian dysgenesis 5, 617690 (3), Autosomal recessive; Spermatogenic failure 32, 618115 (3), Autosomal dominant |
| SOST | 50964 | ENSG00000167941 | [illegible] dominant |
| SOX18 | 54345 | ENSG00000203883 | Hypotrichosis-lymphedema-telangiectasia syndrome, 607823 (3), Autosomal recessive; Hypotrichosis-lymphedema-telangiectasia-renal defect syndrome, 137940 (3), Autosomal dominant |
| SPG7 | 6687 | ENSG00000197912 | Spastic paraplegia 7, autosomal recessive, 607259 (3), Autosomal dominant, Autosomal recessive |
| SPINK1 | 6690 | ENSG00000164266 | Autosomal dominant; {Fibrocalculous pancreatic diabetes, susceptibility to}, 608189 (3), Autosomal dominant, Autosomal recessive |
| SPR | 6697 | ENSG00000116096 | Dystonia, dopa-responsive, due to sepiapterin reductase deficiency, 612716 (3), ?Autosomal dominant, Autosomal recessive |
| SPTA1 | 6708 | ENSG00000163554 | Spherocytosis, type 3, 270970 (3), Autosomal recessive; Elliptocytosis-2, 130600 (3), Autosomal dominant; Pyropoikilocytosis, 266140 (3), Autosomal recessive |
| SPTBN2 | 6712 | ENSG00000173898 | [illegible] Autosomal recessive |
| SPTSSA | 171546 | ENSG00000165389 | Spastic paraplegia 90A, autosomal dominant, 620416 (3), Autosomal dominant; ?Spastic paraplegia 90B, autosomal recessive, 620417 (3), Autosomal dominant |
| SQSTM1 | 8878 | ENSG00000161011 | Neurodegeneration with ataxia, dystonia, and gaze palsy, childhood-onset, 617145 (3), Autosomal recessive; Frontotemporal dementia and/or amyotrophic lateral sclerosis 3, 616437 (3), Autosomal dominant; Myopathy, distal, with rimmed vacuoles, 617158 (3), Autosomal dominant; Paget disease of bone 3, 167250 (3), Autosomal dominant |
| STAT1 | 6772 | ENSG00000115415 | Immunodeficiency 31C, chronic mucocutaneous candidiasis, autosomal dominant, 614162 (3), Autosomal dominant; Immunodeficiency 31A, mycobacteriosis, autosomal dominant, 614892 (3), Autosomal dominant; Immunodeficiency 31B, mycobacterial and viral infections, autosomal recessive, 613796 (3), Autosomal recessive |
| STAT5B | 6777 | ENSG00000173757 | hormone insensitivity with immune dysregulation 2, autosomal dominant, 618985 (3), Autosomal dominant; Leukemia, acute promyelocytic, somatic, 102578 (3) |
| STIM1 | 6786 | ENSG00000167323 | Myopathy, tubular aggregate, 1, 160565 (3), Autosomal dominant; Stormorken syndrome, 185070 (3), Autosomal dominant; Immunodeficiency 10, 612783 (3), Autosomal recessive |

| | | | |
|---|---|---|---|
| STT3A | 3703 | ENSG00000134910 | Congenital disorder of glycosylation, type Iw, autosomal dominant, 619714 (3), Autosomal dominant; Congenital disorder of glycosylation, type Iw, autosomal recessive, 615596 (3), Autosomal recessive |
| STUB1 | 10273 | ENSG00000103266 | Autosomal recessive |
| STXBP1 | 6812 | ENSG00000136854 | Developmental and epileptic encephalopathy 4, 612164 (3), Autosomal dominant, Autosomal recessive |
| SUFU | 51684 | ENSG00000107882 | recessive; Basal cell nevus syndrome 2, 620343 (3); {Medulloblastoma}, 155255 (3), Somatic mutation, Autosomal dominant, Autosomal recessive |
| SYNE1 | 23345 | ENSG00000131018 | autosomal dominant, 612998 (3), Autosomal dominant; Spinocerebellar ataxia, autosomal recessive 8, 610743 (3), Autosomal recessive |
| SYT2 | 127833 | ENSG00000143858 | Myasthenic syndrome, congenital, 7A, presynaptic, and distal motor neuropathy, autosomal dominant, 616040 (3), Autosomal dominant; Myasthenic syndrome, congenital, 7B, presynaptic, autosomal recessive, 619461 (3), Autosomal recessive |
| TBC1D24 | 57465 | ENSG00000162065 | Deafness, autosomal recessive 86, 614617 (3), Autosomal recessive; Epilepsy, rolandic, with paroxysmal exercise-induce dystonia and writer's cramp, 608105 (3), Autosomal recessive; Myoclonic epilepsy, infantile, familial, 605021 (3), Autosomal recessive; Deafness, autosomal dominant 65, 616044 (3), Autosomal dominant; Developmental and epileptic encephalopathy 16, 615338 (3), Autosomal recessive; DOORS syndrome, 220500 (3), Autosomal recessive |
| TBX4 | 9496 | ENSG00000121075 | Ischiocoxopodopatellar syndrome with or without pulmonary arterial hypertension, 147891 (3), Autosomal dominant; Amelia, posterior, with pelvic and pulmonary hypoplasia syndrome, 601360 (3), Autosomal recessive |
| TBX6 | 6911 | ENSG00000149922 | Spondylocostal dysostosis 5, 122600 (3), Autosomal dominant, Autosomal recessive |
| TBXT | 6862 | ENSG00000164458 | (3), Autosomal dominant |
| TCAP | 8557 | ENSG00000173991 | Cardiomyopathy, hypertrophic, 25, 607487 (3), Autosomal dominant; Muscular dystrophy, limb-girdle, autosomal recessive 7, 601954 (3), Autosomal recessive |
| TCF12 | 6938 | ENSG00000140262 | Craniosynostosis 3, 615314 (3), Autosomal dominant; Hypogonadotropic hypogonadism 26 with or without anosmia, 619718 (3), Autosomal dominant, Autosomal recessive |
| TCF3 | 6929 | ENSG00000071564 | Agammaglobulinemia 8B, autosomal recessive, 619824 (3), Autosomal recessive; Agammaglobulinemia 8A, autosomal dominant, 616941 (3), Autosomal dominant |
| TECTA | 7007 | ENSG00000109927 | Autosomal recessive |
| TERT | 7015 | ENSG00000164362 | Dyskeratosis congenita, autosomal dominant 2, 613989 (3), Autosomal dominant, Autosomal recessive; Dyskeratosis congenita, autosomal recessive 4, 613989 (3), Autosomal dominant, Autosomal recessive; Pulmonary fibrosis and/or bone marrow failure syndrome, telomere-related, 1, 614742 (3), Autosomal dominant; {Melanoma, cutaneous malignant, 9}, 615134 (3), Autosomal dominant; {Leukemia, acute myeloid}, 601626 (3), Somatic mutation, Autosomal dominant |
| TET3 | 200424 | ENSG00000187605 | Beck-Fahrner syndrome, 618798 (3), Autosomal dominant, Autosomal recessive |
| TFG | 10342 | ENSG00000114354 | ?Spastic paraplegia 57, autosomal recessive, 615658 (3), Autosomal recessive; Hereditary motor and sensory neuropathy, Okinawa type, 604484 (3), Autosomal dominant |
| TGFB1 | 7040 | ENSG00000105329 | Inflammatory bowel disease, immunodeficiency, and encephalopathy, 618213 (3), Autosomal recessive; Camurati-Engelmann disease, 131300 (3), Autosomal dominant; {Cystic fibrosis lung disease, modifier of}, 219700 (3), Autosomal recessive |
| THPO | 7066 | ENSG00000090534 | Thrombocythemia 1, 187950 (3), Autosomal dominant; Thrombocytopenia 9, 620478 (3), Autosomal dominant; Amegakaryocytic thrombocytopenia, congenital, 2, 620481 (3), Autosomal recessive |
| THRB | 7068 | ENSG00000151090 | Thyroid hormone resistance, autosomal recessive, 274300 (3), Autosomal recessive; Thyroid hormone resistance, 188570 (3), Autosomal dominant; Thyroid hormone resistance, selective pituitary, 145650 (3), Autosomal dominant |
| THSD1 | 55901 | ENSG00000136114 | recessive |
| TIA1 | 7072 | ENSG00000116001 | Welander distal myopathy, 604454 (3), Autosomal dominant, Autosomal recessive; Amyotrophic lateral sclerosis 26 with or without frontotemporal dementia, 619133 (3), Autosomal dominant |

| | | | |
|---|---|---|---|
| TICAM1 | 148022 | ENSG00000127666 | recessive |
| TLR3 | 7098 | ENSG00000164342 | dominant, Autosomal recessive |
| TMC1 | 117531 | ENSG00000165091 | Autosomal recessive |
| TNFRSF11A | 8792 | ENSG00000141655 | Osteopetrosis, autosomal recessive 7, 612301 (3), Autosomal recessive; {Paget disease of bone 2, early-onset}, 602080 (3), Autosomal dominant; Osteolysis, familial expansile, 174810 (3), Autosomal dominant |
| TNFRSF13B | 23495 | ENSG00000240505 | 2, 609529 (3) |
| TNNI3 | 7137 | ENSG00000129991 | ?Cardiomyopathy, dilated, 2A, 611880 (3), Autosomal recessive; Cardiomyopathy, hypertrophic, 7, 613690 (3), Autosomal dominant; Cardiomyopathy, familial restrictive, 1, 115210 (3), Autosomal dominant; Cardiomyopathy, dilated, 1FF, 613286 (3) |
| TNNT1 | 7138 | ENSG00000105048 | severe infantile, 605355 (3), Autosomal recessive; Nemaline myopathy 5B, autosomal recessive, childhood-onset, 620386 (3), Autosomal recessive |
| TNXB | 7148 | ENSG00000168477 | dominant |
| TOR1A | 1861 | ENSG00000136827 | 128100 (3), Autosomal dominant |
| TPM3 | 7170 | ENSG00000143549 | Congenital myopathy 4A, autosomal dominant, 255310 (3), Autosomal dominant; Congenital myopathy 4B, autosomal recessive, 609284 (3), Autosomal recessive |
| TREX1 | 11277 | ENSG00000213689 | Vasculopathy, retinal, with cerebral leukoencephalopathy and systemic manifestations, 192315 (3), Autosomal dominant; Aicardi-Goutieres syndrome 1, dominant and recessive, 225750 (3), Autosomal dominant, Autosomal recessive; {Systemic lupus erythematosus, susceptibility to}, 152700 (3), Autosomal dominant; Chilblain lupus, 610448 (3), Autosomal dominant |
| TSHR | 7253 | ENSG00000165409 | Hyperthyroidism, familial gestational, 603373 (3), Autosomal dominant; Hyperthyroidism, nonautoimmune, 609152 (3), Autosomal dominant; Thyroid adenoma, hyperfunctioning, somatic (3); Hypothyroidism, congenital, nongoitrous, 1, 275200 (3), Autosomal recessive; Thyroid carcinoma with thyrotoxicosis, somatic (3) |
| TTC21B | 79809 | ENSG00000123607 | Short-rib thoracic dysplasia 4 with or without polydactyly, 613819 (3), Autosomal recessive; Nephronophthisis 12, 613820 (3), Autosomal dominant, Autosomal recessive |
| TTN | 7273 | ENSG00000155657 | Muscular dystrophy, limb-girdle, autosomal recessive 10, 608807 (3), Autosomal recessive; Cardiomyopathy, familial hypertrophic, 9, 613765 (3), Autosomal dominant; Congenital myopathy 5 with cardiomyopathy, 611705 (3), Autosomal recessive; Tibial muscular dystrophy, tardive, 600334 (3), Autosomal dominant; Cardiomyopathy, dilated, 1G, 604145 (3), Autosomal dominant; Myopathy, myofibrillar, 9, with early respiratory failure, 603689 (3), Autosomal dominant |
| TUBB8 | 347688 | ENSG00000261456 | Oocyte/zygote/embryo maturation arrest 2, 616780 (3), Autosomal dominant, Autosomal recessive |
| TWIST2 | 117581 | ENSG00000233608 | Ablepharon-macrostomia syndrome, 200110 (3), Autosomal dominant; Barber-Say syndrome, 209885 (3), Autosomal dominant; Focal facial dermal dysplasia 3, Setleis type, 227260 (3), Autosomal recessive |
| TWNK | 56652 | ENSG00000107815 | ophthalmoplegia with mitochondrial DNA deletions, autosomal dominant 3, 609286 (3), Autosomal dominant; Perrault syndrome 5, 616138 (3), Autosomal recessive |
| TYR | 7299 | ENSG00000077498 | blue/green eyes], 601800 (3), Autosomal dominant; {Melanoma, cutaneous malignant, susceptibility to, 8}, 601800 (3), Autosomal dominant; Albinism, oculocutaneous, type IB, 606952 (3), Autosomal recessive; Albinism, oculocutaneous, type IA, 203100 (3), Autosomal recessive |
| UCHL1 | 7345 | ENSG00000154277 | {?Parkinson disease 5, susceptibility to}, 613643 (3), Autosomal dominant; Spastic paraplegia 79A, autosomal dominant, 620221 (3), Autosomal dominant; Spastic paraplegia 79B, autosomal recessive, 615491 (3), Autosomal recessive |
| UCP3 | 7352 | ENSG00000175564 | {Obesity, severe, and type II diabetes}, 601665 (3), Multifactorial, Autosomal dominant, Autosomal recessive |
| UFSP2 | 55325 | ENSG00000109775 | ?Hip dysplasia, Beukes type, 142669 (3), Autosomal dominant; Spondyloepimetaphyseal dysplasia, Di Rocco type, 617974 (3), Autosomal dominant; Developmental and epileptic encephalopathy 106, 620028 (3), Autosomal recessive |

| | | | |
|---|---|---|---|
| UGT1A1 | 54658 | ENSG00000241635 | Crigler-Najjar syndrome, type I, 218800 (3), Autosomal recessive; [Bilirubin, serum level of, QTL1], 601816 (3); Hyperbilirubinemia, familial transient neonatal, 237900 (3), Autosomal dominant, Autosomal recessive; Crigler-Najjar syndrome, type II, 606785 (3), Autosomal recessive; [Gilbert syndrome], 143500 (3), Autosomal recessive |
| UNC45B | 146862 | ENSG00000141161 | ?Cataract 43, 616279 (3), Autosomal dominant; Myofibrillar myopathy 11, 619178 (3), Autosomal recessive |
| UROD | 7389 | ENSG00000126088 | Porphyria, hepatoerythropoietic, 176100 (3), Autosomal dominant, Autosomal recessive; Porphyria cutanea tarda, 176100 (3), Autosomal dominant, Autosomal recessive |
| VAMP1 | 6843 | ENSG00000139190 | Autosomal dominant |
| VHL | 7428 | ENSG00000134086 | syndrome, 193300 (3), Autosomal dominant; Renal cell carcinoma, somatic, 144700 (3); Pheochromocytoma, 171300 (3), Autosomal dominant |
| VKORC1 | 79001 | ENSG00000167397 | 122700 (3), Autosomal dominant |
| VWF | 7450 | ENSG00000110799 | von Willebrand disease, type 1, 193400 (3), Autosomal dominant; von Willebrand disease, types 2A, 2B, 2M, and 2N, 613554 (3), Autosomal dominant, Autosomal recessive; von Willebrand disease, type 3, 277480 (3), Autosomal recessive |
| WARS1 | 7453 | ENSG00000140105 | Neuronopathy, distal hereditary motor, autosomal dominant 9, 617721 (3), Autosomal dominant; Neurodevelopmental disorder with microcephaly and speech delay, with or without brain abnormalities, 620317 (3), Autosomal recessive |
| WASHC5 | 9897 | ENSG00000164961 | Autosomal dominant |
| WDR11 | 55717 | ENSG00000120008 | Intellectual developmental disorder, autosomal recessive 78, 620237 (3), Autosomal recessive; Hypogonadotropic hypogonadism 14 with or without anosmia, 614858 (3), Autosomal dominant |
| WFS1 | 7466 | ENSG00000109501 | Deafness, autosomal dominant 6/14/38, 600965 (3), Autosomal dominant; ?Cataract 41, 116400 (3), Autosomal dominant; Wolfram-like syndrome, autosomal dominant, 614296 (3), Autosomal dominant; {Diabetes mellitus, noninsulin-dependent, association with}, 125853 (3), Autosomal dominant; Wolfram syndrome 1, 222300 (3), Autosomal recessive |
| WNK1 | 65125 | ENSG00000060237 | Neuropathy, hereditary sensory and autonomic, type II, 201300 (3), Autosomal recessive; Pseudohypoaldosteronism, type IIC, 614492 (3), Autosomal dominant |
| WNT1 | 7471 | ENSG00000125084 | {Osteoporosis, early-onset, susceptibility to, autosomal dominant}, 615221 (3), Autosomal dominant; Osteogenesis imperfecta, type XV, 615220 (3), Autosomal recessive |
| WNT10A | 80326 | ENSG00000135925 | Schopf-Schulz-Passarge syndrome, 224750 (3), Autosomal recessive; Tooth agenesis, selective, 4, 150400 (3), Autosomal dominant, Autosomal recessive; Ectodermal dysplasia 16 (odontoonychodermal dysplasia), 257980 (3), Autosomal recessive |
| WNT10B | 7480 | ENSG00000169884 | recessive |
| WNT4 | 54361 | ENSG00000162552 | dominant |
| YARS1 | 8565 | ENSG00000134684 | Infantile-onset multisystem neurologic, endocrine, and pancreatic disease 2, 619418 (3), Autosomal recessive; Charcot-Marie-Tooth disease, dominant intermediate C, 608323 (3), Autosomal dominant |
| ZFP57 | 346171 | ENSG00000204644 | Diabetes mellitus, transient neonatal 1, 601410 (3), Autosomal dominant, Autosomal recessive |
| ZNF408 | 79797 | ENSG00000175213 | Retinitis pigmentosa 72, 616469 (3), Autosomal recessive; ?Exudative vitreoretinopathy 6, 616468 (3), Autosomal dominant |
| ZNF423 | 23090 | ENSG00000102935 | dominant, Autosomal recessive |
